# Computational design of a modular protein sense/response system

**DOI:** 10.1101/648485

**Authors:** Anum A. Glasgow, Yao-Ming Huang, Daniel J. Mandell, Michael Thompson, Ryan Ritterson, Amanda L. Loshbaugh, Jenna Pellegrino, Cody Krivacic, Roland A. Pache, Kyle A. Barlow, Noah Ollikainen, Deborah Jeon, Mark J. S. Kelly, James S. Fraser, Tanja Kortemme

**Author notes:** Novozymes A/S, 2880 Bagsværd, Denmark (Roland A. Pache). A.A.G., Y.-M.H. and D.J.M contributed equally.

## Abstract

Sensing and responding to signals is a fundamental ability of living systems, but despite remarkable progress in computational design of new protein structures, there is no general approach for engineering arbitrary new protein sensors. Here we describe a generalizable computational strategy for designing sensor/actuator proteins by building binding sites *de novo* into heterodimeric protein-protein interfaces and coupling ligand sensing to modular actuation *via* split reporters. Using this approach, we designed protein sensors that respond to farnesyl pyrophosphate, a metabolic intermediate in the production of valuable compounds. The sensors are functional *in vitro* and in cells, and the crystal structure of the engineered binding site matches the design model with atomic accuracy. Our computational design strategy opens broad avenues to link biological outputs to new signals.

**One Sentence Summary:** An engineering strategy to design modular synthetic signaling systems that respond to new small molecule inputs.

## MAIN TEXT

In the last two decades there have been impressive demonstrations of computational protein design to create diverse new protein structures spanning helical(*1-5*), alpha-beta(*6-8*) and beta-sheet(*9, 10*) folds. In contrast, progress in our ability to computationally design arbitrary protein function *de novo* lags far behind, with relatively few examples that often required screening of many design variants followed by subsequent experimental optimization(*11, 12*). Moreover, many advanced functions present in nature have not yet been realized by computational protein design. One such unsolved challenge is the *de novo* design of small molecule sensor/actuators in which ligand binding by a protein directly controls changes in downstream functions, a key aspect of cellular signal transduction(*13*).

Fundamentally, sensing and responding to a small molecule signal requires both recognition of the target and linking target recognition to an output response. Exciting recent progress has been made with the design of proteins recognizing new ligands(*10, 11, 14-16*). A general solution to the second problem, coupling ligand recognition to diverse output responses, has remained challenging. Existing approaches have used a ligand that fluoresces upon binding(*10*), engineered the sensor components to be unstable and hence inactive in the absence of the ligand(*14, 17*), or repurposed an allosteric transcription factor(*18*). These strategies constrain the input signals or output responses that can be used, since they require fluorescent ligands, tuning of the energetic balance between ligand binding and protein stabilization, or coupling to a transcriptional output while preserving allosteric mechanisms.

Here we describe a new computational strategy to engineer protein complexes that can sense a small molecule and respond directly using different biological outputs, creating modular sensor/response systems. Distinct from prior work(*10, 11, 14, 15*) that reengineered existing binding sites or placed ligands into preformed cavities, we build small molecule recognition sites *de novo* into heterodimeric protein-protein interfaces, to create new and programmable chemically induced dimerization systems (CIDs). This strategy is inspired by naturally occurring and reengineered CID systems(*19*) that have been widely used but are limited to a small number of existing or similar input molecules. We reasoned that computationally designed synthetic CIDs would similarly link binding of a target small molecule to modular cellular responses through genetically encodable fusions of each sensor half protein to a split reporter (**Fig. 1A**), but would respond to new, user-defined inputs.

**Fig. 1.**
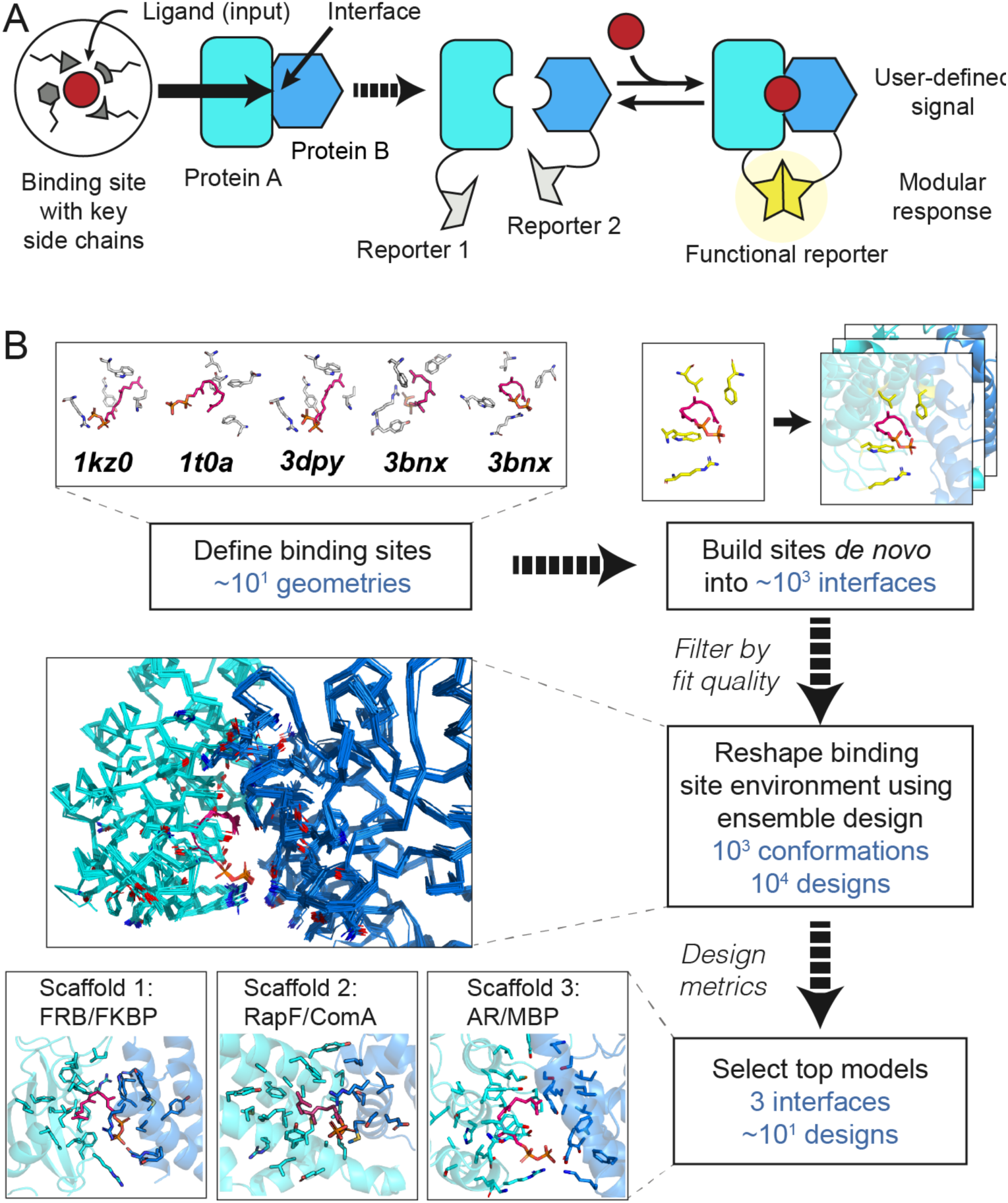
Computational design. (**A**) Cartoon of the design strategy. A small-molecule binding site is built *de novo* into protein-protein interfaces (left) to create new, chemically inducible dimerization systems (CIDs, right). Linking the designed sensor proteins to split reporters yields modular CID system where different reporter outputs can be coupled to user-defined small molecule input signals. (**B**) Steps in the design of a new CID system sensing FPP: (i) binding site geometries with key interacting side chains (“motif residues”) are selected from FPP-binding proteins (pdb codes indicated), (ii) these sites are built computationally into a large number of protein-protein interfaces (“scaffolds”) and (iii) binding sites with feasible geometries are reshaped and optimized by flexible backbone design (shown is a conformational ensemble for a single sequence). (iv) Top designs from 3 different scaffolds (bottom) were selected for experimental tests (**Fig. 2**).

To demonstrate this strategy, we chose farnesyl pyrophosphate (FPP) as the target ligand. FPP is an attractive target because it is a toxic intermediate in a commonly-engineered terpenoid biosynthesis pathway for the production of valuable terpenoid compounds, including the anti-malarial drug artemisinin(*20*). Our computational strategy (**Fig. 1B, Supplemental Methods**) proceeds in four main steps: (i) defining the geometries of minimal FPP binding sites comprised of 3-4 side chains (termed “motif residues”) that form key hydrophobic and hydrogen bonding interactions with the target ligand; (ii) modeling these geometries into a dataset of heterodimeric protein-protein interfaces (termed “scaffolds”) and computationally screening for coarsely compatible scaffolds(*21*); (iii) accommodating the *de novo* built binding sites in these scaffolds using new flexible backbone design methods not previously tested in forward-engineering applications(*22-24*) (“reshaping”); and (iv) ranking individual designs for testing according to several design metrics including ligand binding energy predicted using the Rosetta force field(*25*) and ligand burial.

Starting with 5 FPP binding site geometries and up to 3462 heterodimeric scaffolds, we selected the most highly ranked designs across three engineered scaffolds for a first round of experimental testing (**Fig. 1B, Supplemental Methods**): the FKBP/FRB complex originally responsive to rapamycin(*26*) (1 design), a complex between the bacterial signaling proteins RapF and ComA (*27*) (4 designs) and a synthetic complex between maltose binding protein (MBP) and an ankyrin repeat (AR) protein(*28*) (4 designs) (**Fig. 2A, Table S1, Fig. S1**). While the ligand was placed into the original rapamycin binding site in FKBP/FRB, binding sites in the other two complexes were modeled *de novo*.

**Fig. 2.**
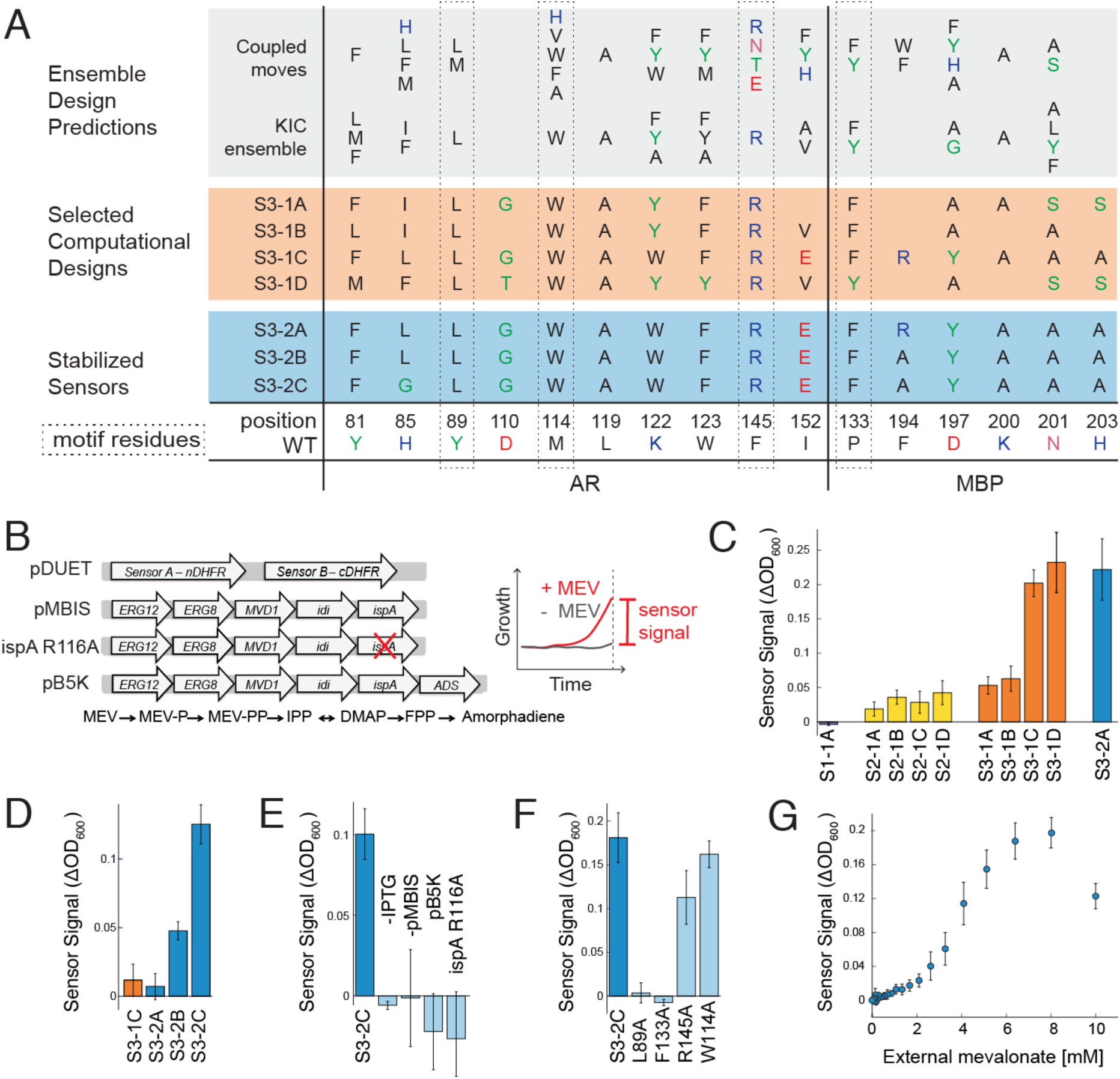
Sensor function in bacteria. **(A)** Designed sequences at key positions for scaffold 3 (see **Table S1, Fig. S1** for all designs and **Appendix 4** for complete sequences). Grey shading: preferred residues from flexible backbone reshaping (**Fig. 1B**). Orange shading: individual computational designs selected based on ligand burial (S3-1A), consensus (S3-1B), optimized ligand packing (S3-1C) and predicted ligand binding score (S3-1D). Blue shading: sensors stabilized by 2 additional mutations from single site saturation mutagenesis (note that designs S3-2B and S3-2C also contained 2 mutations from error-prone PCR that were not in the designed FPP binding site, see **Fig. S1**). **(B)** Constructs (left) used in the split mDHFR reporter assay (right). Cells co-express sensor proteins (pDUET) linked to the split mDHFR reporter with an engineered pathway of 5 enzymes to convert mevalonate (MEV) into FPP (pMBIS)(*20*): mevalonate kinase from *S. cerevisiae* (*ERG12*); phosphomevalonate kinase from *S. cerevisiae* (*ERG8*); mevalonate pyrophosphate decarboxylase from *S. cerevisiae* (*MVD*); IPP isomerase from *E. coli* (*idi*); and FPP synthase from *E. coli* (*ispA*). Plasmid *ispA* R116A contains a single point mutation in the fifth enzyme of the pathway that reduces catalytic activity of *ispA* by 13-fold(*35*). Plasmid pB5K adds a sixth enzyme to the pathway (amorphadiene synthase, ADS) that converts FPP to amorphadiene(*20*). Cells expressing functional sensors confer a growth advantage in the split mDHFR assay through FPP-driven dimerization of the sensor proteins and resulting complementation of functional mDHFR. Sensor signal is quantified as change in OD_600_ in the presence and absence of mevalonate. **(C)** Sensor signal in the split mDHFR assay for designs based on scaffold 1 (FKBP-FRB12, purple bar), scaffold 2 (RapF-ComA, yellow bars) and scaffold 3 (AR-MBP, orange bars). Computational designs S3-1C and S3-1D are strongly responsive to FPP. Sensor S3-2A, which was identified from library 2 and has 2 mutations distal from the designed FPP binding site (**Table S1**), is shown for comparison (blue bar). **(D)** Stability-enhancing mutations in S3-2B and S3-2C improve sensor signal over S3-1C and S3-2A under more stringent conditions (trimethoprim concentration increased to 6 µM compared to 1 µM in panel C). **(E)** Sensor signal for S3-2C is dependent on expression of the sensor proteins (- IPTG), expression of the FPP production pathway (-pMBIS), and specific to FPP (conversion of FPP to amorphadiene by co-expression of ADS in pB5K, or decrease of FPP production by the R116A mutation in *ispA*). **(F)** Mutation of three of the four motif residues to alanine decreases the sensor signal in response to FPP. **(G)** Dependence of the S3-2C sensor signal on concentration of the FPP precursor mevalonate added extracellularly. The decreased growth at high mevalonate concentrations (>8 mM) is likely due to FPP toxicity. Error bars are standard deviation from at least 4 biological replicates and 8 technical replicates for each biological replicate.

To test these computationally designed FPP sensors, we genetically fused the engineered sensor proteins to a well-studied split reporter, the enzyme murine dihydrofolate reductase (mDHFR(*29*), **Fig. 2B, Appendix 1**), and expressed the fusion constructs in *E. coli*. We reasoned that functional sensors should exhibit increased growth due to split mDHFR complementation in the presence of FPP under conditions where the endogenous *E. coli* DHFR protein was specifically inhibited by trimethoprim. Since FPP does not efficiently enter *E. coli*, we added its metabolic precursor, mevalonate, to the growth medium and co-expressed an engineered pathway of 5 enzymes(*20*) (**Fig. 2B**) to produce FPP from mevalonate in the cells. We then monitored sensor function as change in growth in the presence or absence of mevalonate under otherwise identical conditions (**Fig. 2B, Supplemental Methods**). In the following, we denote designs by their scaffold (S1, S2, S3), design generation (1, 2, 3) and consecutive letter (A, B, etc.; for details see **Table S1, Fig. S1**).

While 7 of the 9 selected designs showed only a small (S2-1A, B, C, D; S3-1A, B) or no signal (S1-1A), two designs (S3-1C, D) displayed a robust signal response to FPP (**Fig. 2C, Fig. S2**). Both designs resulted from the MBP-AR scaffold (S3, **Fig. 2A**). For this scaffold, we also generated two libraries: library 1 based on our ensemble design predictions (**Fig. 2A, Table S2**), and library 2 using error-prone PCR starting from design S3-1C. We screened 3×10^5^ members of each library and selected 1536 clones from each library after enrichment for growth via split mDHFR complementation. The selected clones were then subjected to an array-based colony-printing assay (**Supplemental Methods, Fig. S3**). From this assay, we selected 36 hits from which we confirmed 27 sequences (7 from library 1 and 20 from library 2) by individual growth assays (**Fig. S4, Fig. S5**). One of the most active designs across both library screens (S3-2A) was a variant of design S3-1C containing two additional mutations distal from the designed FPP binding site introduced by error-prone PCR. Interestingly, this variant displayed essentially equal activity as the original S3-1C design when tested under identical conditions (**Fig. 2C, Table S1, Fig. S2**). These results demonstrate that library screening or error-prone PCR were not necessary to identify functional sensors; instead, we obtained functional sensors directly *via* computational protein design. However, library 1 provided additional active sequences resulting from sequence tolerance predicted in the ensemble design simulations (**Fig. 2A, Table S2, Fig. S4**).

To further characterize the identified best design, S3-2A (**Table S1**), we performed single site saturation mutagenesis at 11 positions (both previously designed and additional second shell positions). We tested the resulting mutants with the growth-based split mDHFR reporter under more stringent conditions by increasing the trimethoprim concentration (**Table S3, Fig. S6**). While the original, computationally chosen amino acid at most positions appeared to be optimal under these conditions, we saw considerable improvements for mutations at two positions, R194A (design S3-2B) and R194A / L85G (design S3-2C, **Fig. 2A**). Designs S3-2B and S3-2C displayed increasing responses to the presence of mevalonate at higher trimethoprim concentrations (**Fig. 2D**). For the most active design, S3-2C, we confirmed that the sensor signal was dependent on expression of the sensor proteins (**Fig. 2E**, -IPTG, **Appendix 1**) and the presence of the metabolic pathway that converts added mevalonate to FPP (**Fig. 2E**, -pMBIS, **Appendix 1**). To test for specificity for FPP, we confirmed that the sensor signal was absent when preventing the accumulation of FPP by either inactivating the fifth enzyme in the pathway by a single point mutation (**Fig. 2B, E**, ispA R116A, **Appendix 1**) or adding a sixth enzyme that converts FPP to amorphadiene (**Fig. 2B, E**, pB5K, **Appendix 1**). To test whether the sensor signal was dependent on the original four motif side chains, we mutated each individually to alanine and observed decreased sensitivity to the presence of mevalonate for three of the four motif side chains (L89, F133, R145 but not W114; **Fig. 2F, Appendix 1**). Finally, we tested whether the sensor signal of design S3-2C was dependent on the concentration of FPP, using increasing concentrations of mevalonate added extracellularly as a proxy (**Fig. 2G**). Interestingly, while the sensor signal initially increased with increasing concentrations of mevalonate, as expected, the signal decreased again at the highest mevalonate concentration tested. This behavior is likely due to the toxicity of FPP known to decrease growth at this mevalonate concentration (*20*). We confirmed a consistent dependency of the sensor signal on both sensor expression (by adding different amounts of the inducer IPTG) and mevalonate addition to the growth medium for seven of our designs (S3-1A, B, C, D; S3-2A, B, C; **Fig. S7**). Taken together, these results confirmed that sensor function in *E. coli* was specific to FPP produced *via* an engineered pathway, dependent on key residues in the engineered binding site, dose-dependent in *E. coli*, and sensitive to FPP concentrations in a relevant range (i.e. below the toxicity level).

To confirm biochemically that FPP increases the binding affinity of the AR-MBP complex as designed, we purified the designed AR and MBP proteins without attached reporters (**Supplemental Methods**; these constructs contained several previously published mutations to stabilize AR(*30*), which when tested in the split mDHFR reporter assay led to active sensor S3-2D, **Table S1, Fig. S8, Appendix 2**). We determined the apparent binding affinity of the designed AR and MBP proteins comprising the S3-2D sensor (**Fig. 3A, Table S1, Fig. S1**) in the absence and presence of 200 µM FPP using biolayer interferometry with streptavidin-biotin coupling (**Fig. 3B, Fig. S9, Supplemental Methods**). The presence of FPP led to a greater than 100-fold stabilization of the interaction between the AR and MBP proteins comprising sensor S3-2D (from >200 µM to 2.1 µM, **Fig. 3C)**. Binding of FPP to the designed AR component of S3-2D alone was weak and binding of FPP to the designed MBP component of S3-2D alone not detectable (**Fig. 3D**). Taken together, these results confirm *in vitro* with purified components that design S3-2D functions as a CID system responding to FPP.

**Fig. 3.**
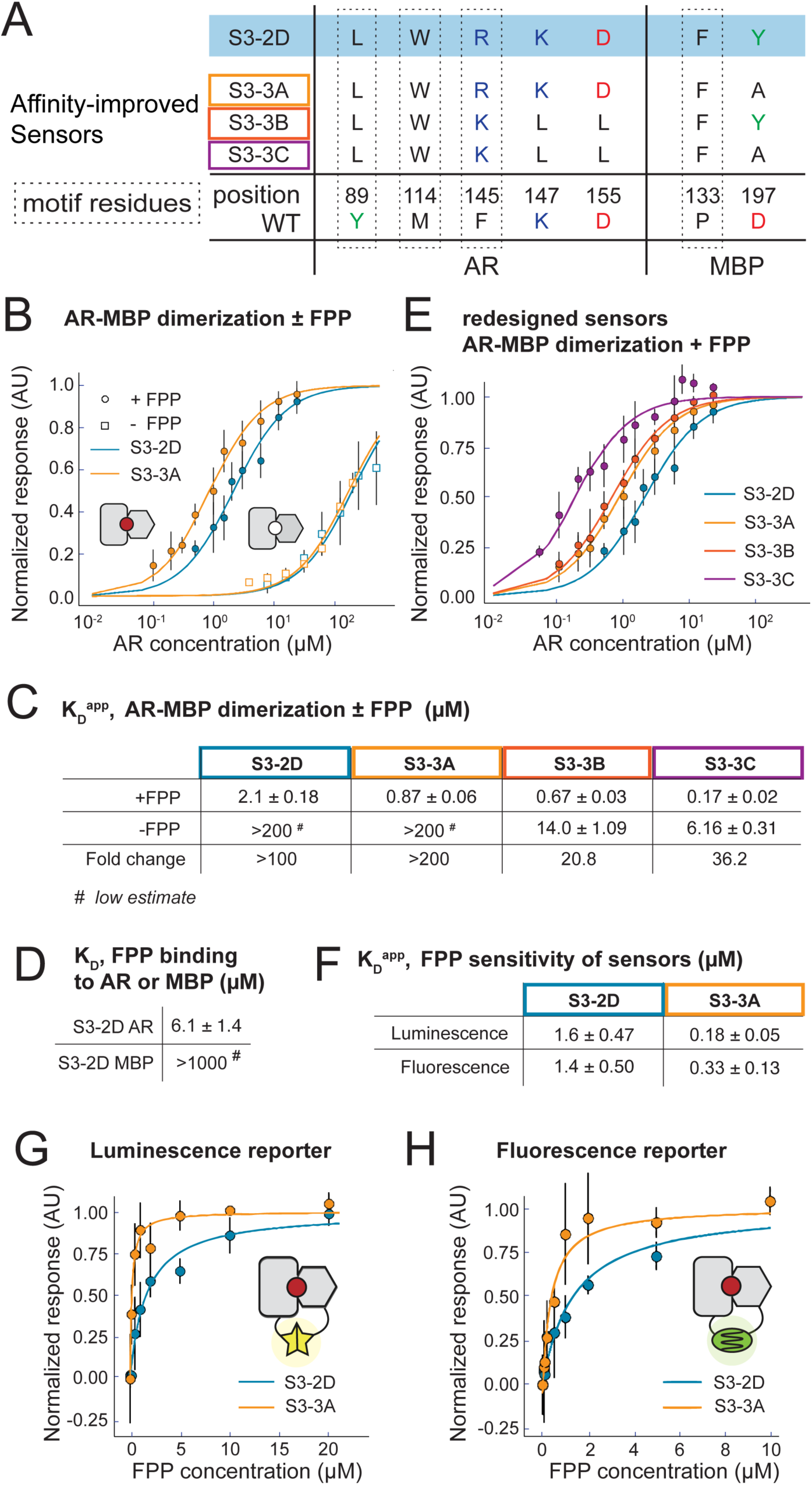
Sensor characterization *in vitro* and output modularity. **(A)** Sequence changes in sensor constructs tested *in vitro*. Motif residues are also shown. The starting construct, S3-2D (blue), is identical to S3-2C in the engineered FPP binding site but contains additional previously published stabilizing mutations in AR(*30*) (shown in **Table S1**). Variants S3-3A-C contain mutations in S3-2D computationally designed to improve the stability of the ternary S3-2D complex with FPP. **(B-H)** *In vitro* binding measurements from biolayer interferometry (BLI) using purified protein (panels **B-E**) or FPP titrations with sensors expressed by *in vitro* transcription / translation (TxTl) (panels **F-H**). **(B)** Apparent AR interaction with immobilized MBP in the presence (closed circles) or absence (open squares) of 200 µM FPP, comparing designs S3-2D (blue) and S3-3A containing the Y197A mutation (orange). (**C**) Summary of BLI results for apparent AR-MBP dimerization with and without FPP. (**D**) Summary of BLI results for FPP binding to the individual designed AR and MBP proteins comprising design S3-2D (**Table S1**). (**E**) Apparent AR interaction with immobilized MBP for a variant derived from local computational design of the FPP binding site using the S3-2D crystal structure as the input, with (purple, S3-3C) or without (red, S3-3B) the Y197A mutation; design S3-3C further shifted the apparent AR-MBP affinity to 170 nM but reduced sensor dynamic range (panel **C**, right). **(F)** Apparent affinity of the S3-2D and S3-3A sensors for FPP using two different reporters in TxTl experiments: nanoBiT, a split luciferase system(*32*), and ddGFP, a dimerization-dependent fluorescent protein pair(*31*). (**G, H**) FPP titration experiments in TxTl with the luminescence reporter (**G**) and the fluorescence reporter (**H**). Error bars are standard deviations for *n* ≥ 3.

To determine whether FPP is recognized in the *de novo* engineered binding site as predicted by the design model, we determined a 2.2 Å resolution crystal structure of the ternary complex of FPP bound in the engineered AR-MBP interface (**Supplemental Methods**; **Table S4**). The crystal structure of the bound complex is in excellent overall agreement with the design model (**Fig. 4A-C**). Despite twinning in the crystals, examining unbiased omit maps allowed modeling of unexplained density in the engineered binding site as FPP (**Fig. 4B, Fig. S10**) and confirmed the side chain conformations in the designed binding pocket (**Fig. 4C, D**). Overall, in a 10 Å shell around FPP in the binding pocket, the Cα root mean squared deviation (rmsd) between the model and the structure is 0.53 Å and the all heavy atom rmsd is 1.13 Å. While crystals formed only in the presence of FPP, only one of the two complexes in the asymmetric unit contained FPP in the binding site (**Fig. S11**). This behavior allowed us to compare *apo* and *holo* states of the complex. The majority of the designed side chains are in identical conformations in the FPP-bound *holo* and FPP-minus *apo* states (**Fig. 4E**), suggesting favorable pre-organization of the designed binding site. An exception is W114 on AR that is partly disordered in the apo state (**Fig. S11**), providing a potential explanation for why a W114A mutation is not as detrimental for sensor activity (**Fig. 2F**) as expected based on the observed packing interactions between W114 and FPP in the *holo* state. A second slight deviation between the model and the crystal structure appeared to be caused by potential steric clashes of the engineered Y197 on MBP with the modeled FPP conformer, which led to re-arrangements in the FPP structure and a rotamer change in another designed residue on MBP, F133 (**Fig. 4D**). Interestingly, many of the original models from computational design favored a smaller alanine side chain at this position (**Fig. 2A**). These observations led to the prediction that a Y197A mutation might stabilize the ternary complex, and indeed design S3-3A containing the Y197A mutation showed an increased (>200 fold) stabilization of the complex with FPP, with an apparent dissociation constant of the designed AR and MBP proteins comprising sensor S3-3A in the presence of 200 µM FPP of 870 ± 60 nM (**Fig. 3B, C**). We also confirmed that design S3-3A (**Table S1**) is active in *E. coli* (**Fig. S12**). To further improve the design based on the crystal structure of design S3-2D, we employed an additional round of flexible backbone design using the Rosetta “CoupledMoves” method(*24*) starting from the FPP-bound crystal structure. These simulations suggested 3 additional mutations leading to design S3-3B: R145K, K147L, D155L (**Fig. 3A**). These mutations, when combined with the Y197A mutation (design S3-3C), enhanced the apparent binding affinity of the designed AR and MBP proteins comprising sensor S3-3C in the presence of 200 µM FPP to 170 ± 20 nM (**Fig. 3C, E**), but also strengthened the binding affinity of the protein-protein dimer in the absence of FPP to 6.2 ± 0.3 µM (**Fig. S13**). Taken together, the crystal structure confirmed the engineered *de novo* binding site at atomic accuracy and provided key insights leading to further computational predictions that improved the apparent binding affinity of the sensor proteins in the presence of FPP to the nanomolar range (sensor S3-3A).

**Fig. 4.**
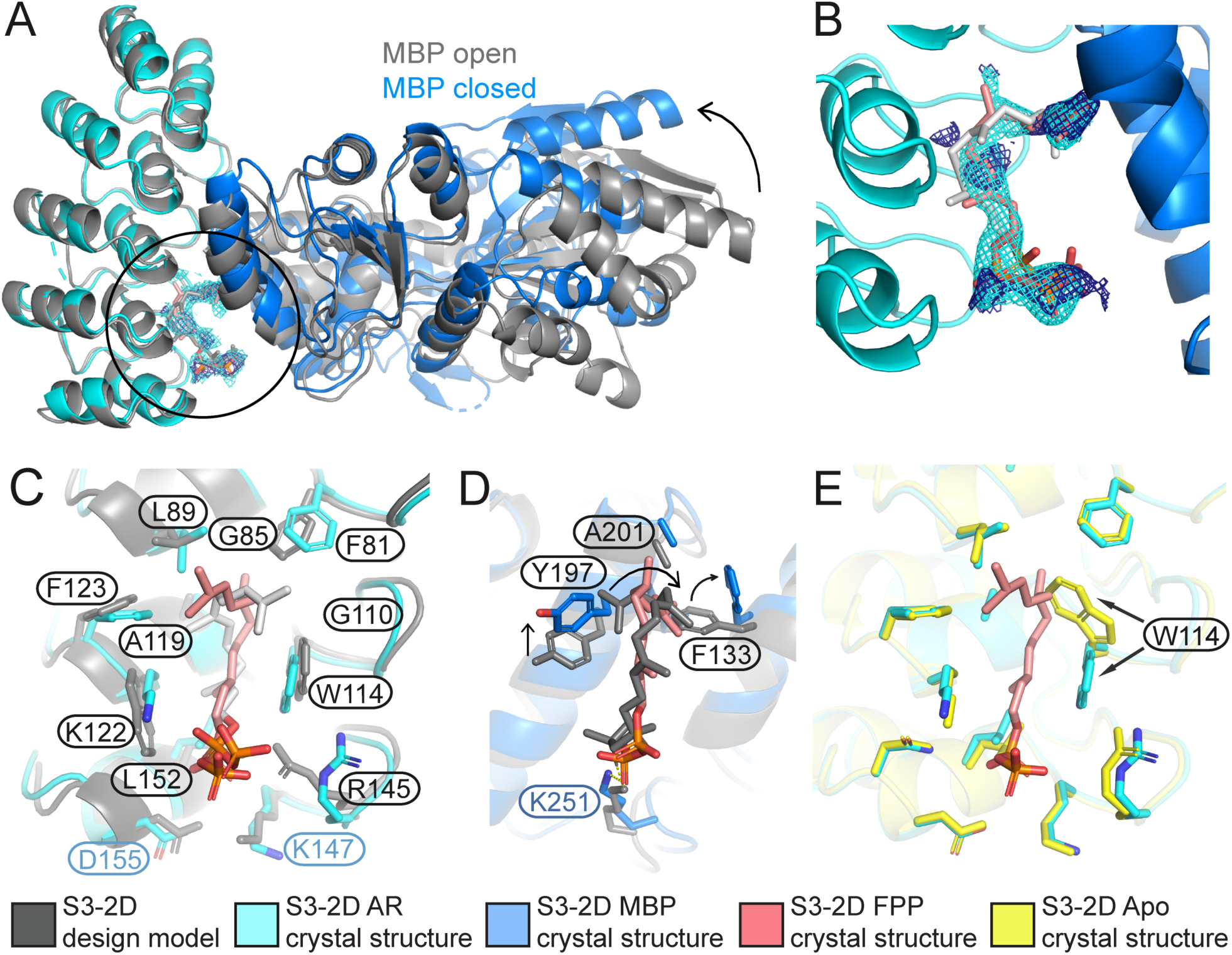
The S3-2D crystal structure matches the computational design with atomic accuracy. **(A)** Overlay of the design model (grey) with the crystal structure (designed AR: cyan, designed MBP: blue, FPP: pink) showing FPP binding in the computationally designed binding site at the AR-MBP interface (circle). Note that the designs crystallized in the open MBP conformation while MBP was in the closed conformation in the original scaffold on which the model was based, leading to a difference in rigid-body orientation (arrow) of one lobe of MBP distal to the FPP binding site. **(B)** FPP overlaid with *2F*_*o*_*-F*_*c*_ electron density map (1.2σ, cyan) and ligand omit map (0.8σ, dark blue). Strong density peaks were present in both maps for the phosphates and several anchoring hydrophobic groups. **(C)** Open-book representation of FPP binding site on AR, showing close match of designed side chain conformations. **(D)** Open-book representation of FPP binding site on MBP, indicating a clash between the position of MBP Y197 in the crystal structure (blue) and the designed FPP orientation in the model (grey), causing slight rearrangements of FPP and F133 (arrows). **(E)** Alignment of the holo (cyan) and apo (yellow) structures of S3-2D, showing overall agreement with the exception of the side chain of W114 (arrows). In panels (**C-E**), residues are labeled black when designed and green/blue when present in the original scaffold complex.

A main advantage of our CID design strategy is the ability to link an engineered sensor, whose input is specific to a user-defined small molecule signal, to a modular output that can in principle be chosen from many available split reporters (**Fig. 1A**). To test this concept, we linked the engineered CID sensors S3-2D and S3-3A to two additional outputs, a dimerization-dependent fluorescent protein(*31*) and split luciferase(*32*) (**Fig. 3G, H, Appendix 3**). We tested input-output responses with the two different reporters using an *in vitro* transcription-translation system (TxTl) (*33*) in which FPP can be added at defined concentrations to the assay extract, in contrast to the cell-based split mDHFR assay. The TxTl assay revealed a nanomolar FPP sensitivity (K_D_^app^) for our best sensor S3-3A (**Fig. 3F**) that is essentially identical for both reporters (180 ± 50 and 330 ± 130 nM by luminescence and fluorescence detection, respectively, **Fig. 3G, H**), and additionally confirms the improvements in design S3-3A containing the Y197A mutation over design S3-2D (the K_D_^app^ for S3-2D was 1.6 ± 0.5 µM and 1.4 ± 0.5 µM for the luminescence and fluorescence reporters, respectively, **Fig. 3F, G, H**). These results show that our CID sensor design strategy is compatible with modular outputs.

Taken together, our results demonstrate the first computational *de novo* design at atomic accuracy of a functional three-part biological sensing system, achieved by reprogramming protein-protein interfaces to respond to a new small molecule ligand. The designed sensor/actuators are conceptually similar to naturally occurring CID systems that allow diverse organisms to respond to changes in their intra-and extracellular environments. We reached an apparent sensitivity to the input ligand FPP in the nanomolar range by computational design without the need for extensive experimental screening. The resulting suite of FPP biosensors are (i) active *in vitro* (**Fig. 3E, Fig. S13**), (ii) functional over a relevant dynamic range in cells that produce FPP from metabolic precursors (**Fig. 2G**), and (iii) have modular compatibility with several reporter outputs (**Fig. 2B, C, Fig. 3G, H**).

While protein-protein interfaces are sometimes considered undruggable, our method demonstrates that small molecule binding sites can be built into these interfaces *de novo*. A prior computational analysis suggests that the appearance of pockets around artificially generated protein-protein interfaces may be an intrinsic geometric feature of protein structure(*34*), lending support to the idea that our approach is extensible to many other ligands and interfaces. The design method presented here hence introduces a generalizable way to create new molecular interactions and output responses with unique specificities that can be used in diverse biological contexts and that respond to user-defined molecular signals.

## Acknowledgments

We would like to thank: Jay Keasling and Fuzhong Zhang for advice on FPP production in microbes and pathway constructs; Emzo de los Santos, Zach Sun, Vincent Noireux, Richard Murray for TxTl advice and reagents; Spencer Alford and Robert Campbell’s lab for ddFP constructs; Aditya Anand, Victor Ruiz, Ben Adler and Alison Maxwell for contributions to computational design and characterization; Shane O’Connor for developing a database for design models; and members of the Kortemme lab for discussion.

## Author contributions

DJM and TK conceived the idea for the project; DJM developed and performed the majority of the computational design with contributions from AAG, RP, KB, NO, and JP; AAG and YMH designed the experimental approach and performed the majority of the experimental characterization, with contributions from RR, AL, CK, DJ and MJSK. MT and JSF determined the crystal structure and MJSK, JSF and TK provided guidance, mentorship and resources. AAG and TK wrote the manuscript with contributions from all authors.

## Data and materials availability

All relevant data are available in the main text or the supplementary materials. Upon publication, constructs will be made available via Addgene.

