## Supplementary Material for "Computational design of a modular protein sense/response system"

**This PDF file includes:**

Materials and Methods
Figs. S1 to S13
Tables S1 to S4
Appendices 1-4
References

**Supplementary Figures**

**Figure S1:** **Sequence alignments of designed proteins (Table S1).**
**Figure S2:** **Growth +/- 5 mM mevalonate and change in growth for data shown in Figure 2C.**
**Figure S3:** **Example data from library screening using a colony-printing assay.**
**Figure S4:** **Sequence alignments of hits from the computationally designed S3 library.**
**Figure S5:** **Sequence alignments of hits from the error-prone PCR S3 library.**
**Figure S6:** **Data from the single-site saturation mutagenesis screen.**
**Figure S7:** **Dependency of the sensor signal on IPTG and mevalonate concentrations.**
**Figure S8:** **Growth +/- mevalonate and change in growth for design S3-2D.**
**Figure S9:** **BLI binding assay representative data and fits.**
**Figure S10:** **Electron density maps supporting placement of the FPP ligand.**
**Figure S11:** **Electron density supporting ligand-induced rearrangement of the FPP binding site.**
**Figure S12:** **Comparison of S3-2D and S3-3A (with Y197A mutation) in the split mDHFR assay.**
**Figure S13:** **BLI binding data +/- FPP for designs S3-2D, S3-3A, S3-3B, S3-3C.**

**Supplementary Tables**

**Table S1:** **Summary of computational designs.**
**Table S2:** **Computationally designed library.**
**Table S3:** **Single-site saturation mutagenesis positions and results.**
**Table S4:** **X-ray data reduction and model refinement.**

**Appendix 1:** **Plasmids / constructs for split mDHFR reporter assay.**
**Appendix 2:** **Plasmids / constructs for *in vitro* binding experiments and crystallography.**
**Appendix 3:** **Plasmids / constructs for TxTl reporter assay.**
**Appendix 4:** **Gene and vector DNA sequences.**

**Supplementary References**

**Materials and Methods**

1. Computational design methods

Computational methods that resulted in the successful sensor design for scaffold 3 (AR-MBP) are as described in detail first(1). Differences in the design protocol that resulted in designs for scaffold 1 (FKBP12-FRB) and scaffold 2 (RapF-ComA) are described further below.

1.1 Selection of protein-protein interface scaffold set. To identify protein interface scaffolds suitable for accommodating small molecule binding sites, we searched the PDB for heterodimeric complexes with ≤95% sequence identity solved at ≤2.8 Å resolution between chains of 75 to 300 residues that were expressed in *E. coli*. This search resulted in 612 structures that were filtered to remove HETATM records and multiple densities (only the first densities listed were kept). Selenomethionines were converted to methionines.

1.2 Definition of FPP binding site geometries. We identified 29 X-ray structures of FPP–protein complexes at ≤2.8 Å resolution in the PDB to serve as templates for binding site geometries. We visually inspected each protein-FPP complex to identify cases where 4 residues could define an encompassing portion of the FPP binding surface. 18 structures were discarded because FPP bound in complex with an inhibitor or other small molecule, forming a binding site that cannot easily be reproduced by amino acid side-chains. Other cases were discarded because the binding site was formed by small contributions from too many residues to define a suitable binding site geometry. Ultimately, 4 template binding site geometries ("motifs") were selected for subsequent matching and design:

| PDB | Protein | Motif residues |
|---|---|---|
| 1kzo | Protein farnesyltransferase | chain B: R291, Y251, W303.<br>chain C: I10. |
| 1t0a | 2C-Methyl-D-Erythritol-2,4-cyclodiphosphate Synthase | chain A: I101, F9.<br>chain B: F9. chain C: F9. |
| 3bnx | Aristolochene synthase | chain A: R314, W308, L184, F153. |
| 3dpy | Protein farnesyltransferase | chain B: R291, Y251, W303.<br>chain C: I2008. |

Note that for PDB templates 1kzo and 3dpy, one of the motif residues comes from a co-associated peptide substrate, and the binding site geometry from 1t0a contains residues from a homotrimeric interface. In all cases non-polar hydrogens were added to FPP, and a single polar hydrogen was placed on the O5 oxygen.

1.3 Building binding sites *de novo* into protein-protein interfaces. We scanned the interface scaffold set for backbones that may accommodate the small molecule target and binding site geometry using a geometric matching procedure(*2*). For each binding site geometry, the relationship between the motif side-chains and the target is uniquely defined by 6 geometric constraints. The matching algorithm scans the first motif residue constraints across a set of scaffold residue positions (here, all positions with Ca atoms within 15 Å of the other chain). At each position, the motif residue is placed into rotameric conformations from the Dunbrack backbone-dependent rotamer library(*3*). For each side-chain conformation, the small molecule target is placed relative to the motif residue using the geometric constraints defined from the template binding site geometry. Conformations that place the target without introducing steric clashes between the motif side-chain, the target, and the scaffold backbone are recorded. The process is iterated for the remaining motif residues, comparing the clash-free target positions to those from the previous motif residues using an efficient geometric hashing technique. Motif side-chain conformations that place the target within the same geometric 'bin' as positions from the previous motif side-chains are recorded as ‘hits’. At the conclusion, cases where the target is placed into the same geometric bin for all motif residues are called ‘matches’. Only matches where at least one motif side-chain is placed on a different chain than the remaining motif side-chains (i.e., across the scaffold interface) are considered further. Many matches may be found for a set of scaffold interface residue positions, corresponding to highly similar conformations for the motif residue side-chains and the target that fall into the same geometric bin. One such match is randomly selected for design, which is termed a ‘unique match’. The numbers of unique matches arising from each template binding site geometry for FPP are shown below:

| Template PDB | Motif residues | Number of unique matches |
|---|---|---|
| 1kzo | chain B: R291, Y251, W303.<br>chain C: I10. | 79 |
| 1t0a | chain A: I101, F9.<br>chain B: F9. chain C: F9. | 371 |
| 3bnx | chain A: R314, W308, L184, F153. | 370 |
| 3dpy | chain B: R291, Y251, W303.<br>chain C: I2008. | 43 |

The quality and quantity of matching results are tuned by a number of parameters. To slightly relax the angle and torsion constraints, we sampled 5 degrees above and below the values computed from the template binding site geometry. There are also Euclidean and Euler parameters that determine the bin size for geometric hashing, which we set to 2.0 Å and 20.0 degrees, respectively. A bump tolerance parameter allowed for some steric overlap – to be resolved in the design stage – which we set to 0.6. We also allowed motif residues to be matched by other residue types with similar side-chain moieties. The following groups of residues were allowed to be matched by any residue in the group: “DE”, “LVI”, “FYW”, and “ST”. We tuned the matching parameters to produce a reasonable number of unique matches for design (on the order of several hundred). Evaluating matches directly is not necessarily informative, since a good match (one that faithfully reproduces the template binding site geometry) may yield poor designs due to an inability of the ‘second shell’ residues to accommodate the binding site geometry and target, while less precise matches may yield more promising designs after being subjected to rigid body optimization of the ligand and backbone relaxation of the scaffold. Thus, all unique matches are passed on to design.

A number of parameters control Rosetta-specific matching options, including the number of side-chain conformations sampled. The command line options used for Rosetta revision r35441 were as follows:

match.linuxgccrelease -database minirosetta_database -s 1SVX.pdb -match:lig_name LG1 -match:grid_boundary 1SVX.gridlig -match:scaffold_active_site_residues 1SVX.pos -match:geometric_constraint_file 3bnx.cst -extra_res_fa 3bnx_LG.fa.params -output_matches_per_group 10 -ex1 -ex2 -extrachi_cutoff 0 -euclid_bin_size 2.0 -euler_bin_size 20.0 -bump_tolerance 0.6 -match:output_format PDB -match:consolidate_matches -match:output_matchres_only

<u>1.4 Initial designs and analysis.</u> The matching algorithm places the motif residues and target into a scaffold interface while avoiding clashes with the backbone. However, the procedure will likely introduce unfavorable interactions with the residues surrounding the motif, or ‘second shell’ residues. In order to accommodate the target and binding motif, we applied a protocol that iterates between rigid body optimization of the target(*4*) and sequence design of the second shell residues(*5, 6*). In the design step, all residues with a Ca atom within 6.0 Å of any ligand heavy atom were designable (they could change residue type), as well as any residue with a Ca atom within 8.0 Å of any ligand heavy atom that also has a Cb atom closer to the ligand than the Ca

atom. Additionally, all residues with a Ca atom within 10.0 Å of any ligand heavy atom are subject to repacking (Metropolis Monte Carlo optimization of side-chain conformations) together with any residue with a Ca atom within 12.0 Å of the ligand with a Cb atom that is closer to the ligand than the Ca atom. The protocol iterates rigid body optimization and sequence design 3 times, producing interfaces with more favorable interactions between the motif residues, target, and second shell residues.

The command line options used for Rosetta revision r36129 were as follows:

```
EnzdesFixBB.linuxgccrelease -database minirosetta_database -s 1BH9_R33Y94L120F121.pdb -
extra_res_fa 3bnx_LG.fa.params enzdes:detect_design_interface -enzdes:cut1 6.0 -enzdes:cut2
8.0 -enzdes:cut3 10.0 -enzdes:cut4 12.0 -enzdes:cst_opt -enzdes:cst_design -enzdes:cst_min -
enzdes:cstfile 3bnx.cst -enzdes:bb_min -enzdes:chi_min -enzdes:design_min_cycles 3 -ex1 -ex2
-use_input_sc  -nstruct 999 -enzdes:start_from_random_rb_conf
```

For each template binding site geometry we produced on the order of $10^4$ designs and created distributions over computed physicochemical properties. Four of these distributions from the 3bnx template binding site geometry are shown below:

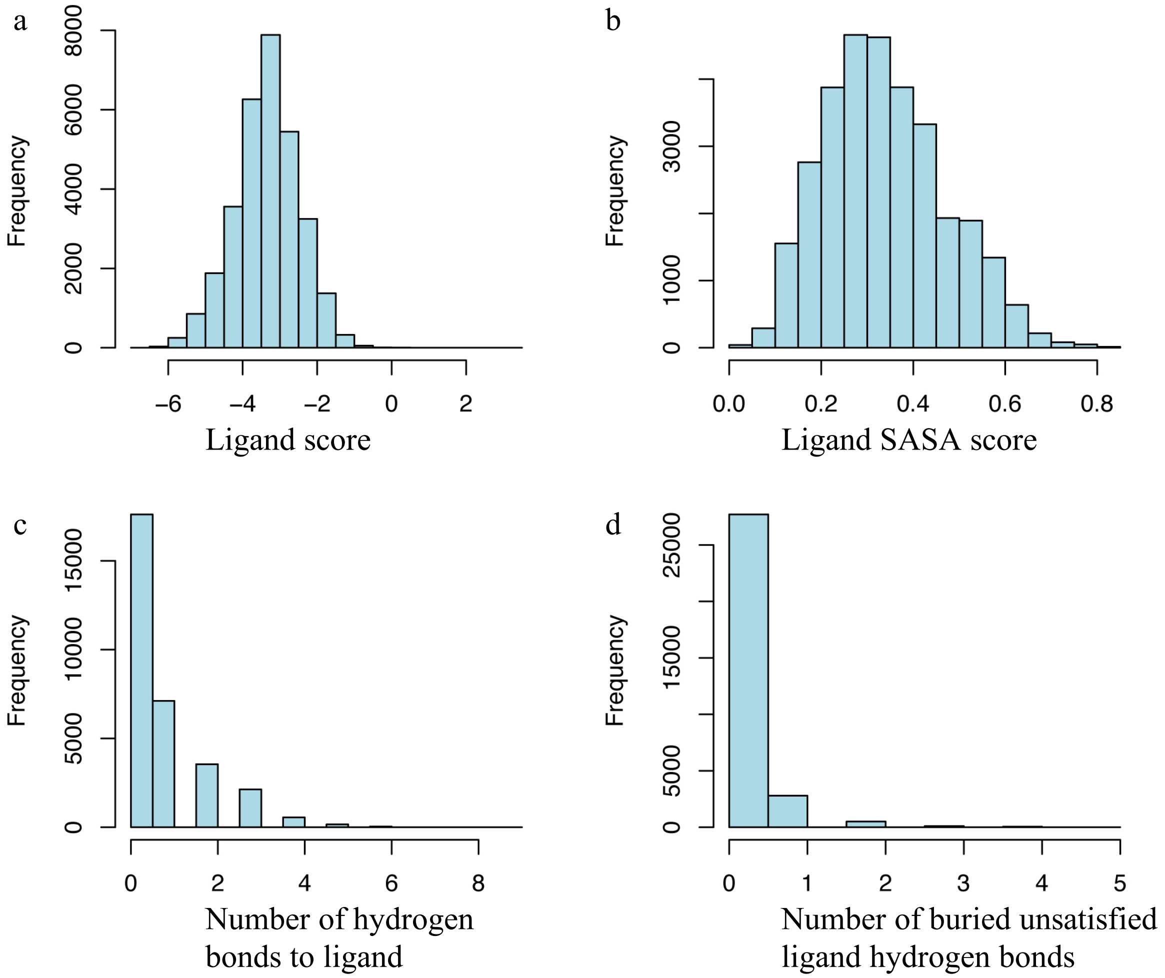

The ligand score (panel a) corresponds to the predicted binding energy between the ligand and scaffold interface. The ligand solvent accessible surface area (SASA, panel b) score measures the burial of the ligand from 0.0 (completely solvent exposed) to 1.0 (completely buried). The number of hydrogen bonds between the scaffold and the ligand (panel c) and the number of buried unsatisfied hydrogen bonds on the ligand (panel d) are also shown. For FPP, visual inspection of representative members of the distributions across all designs suggested the following filter for selecting designs for further refinement: ligand score < -6.0, ligand SASA > 0.6, ligand hydrogen bonds > 1, unsatisfied buried ligand hydrogen bonds = 0. The number of designs passing the filter for all FPP binding site geometries is given below:

| Template PDB | Motif residues | Number of unique matches | Number of designs passing filter |
|---|---|---|---|
| 1kzo | chain B: R291, Y251, W303. chain C: I10 | 79 | 31 |
| 1t0a | chain A: I101, F9. chain B: F9. chain C: F9. | 371 | 0 (1 if SASA filter relaxed to 0.5) |
| 3bnx | chain A: R314, W308, L184, F153. | 370 | 81 |
| 3dpy | chain B: R291, Y251, W303. chain C: I2008. | 43 | 0 (2 if SASA filter relaxed to 0.5) |

Passing designs were then further filtered to remove interface scaffolds imposing additional challenges such as cases with small molecules crystallized at the predicted target binding site and complexes that were purified from inclusion bodies.

Ultimately, a design passing all filters with the best ligand score on an interface scaffolds, the complex (PDB 1svx) between an engineered ankyrin repeat protein (AR) and maltose binding protein (MBP) with a 4-residue binding site geometry from template 3bnx, was selected for refinement by flexible backbone ensemble design.

1.5 Flexible backbone ensemble design. To model the conformational adjustments that could occur in concert with sequence mutations, matched scaffold designs were subject to kinematic closure (KIC(*7*)) over their entire backbones producing conformational ensembles. In brief, KIC generates backbone conformations of segments in proteins by sampling backbone phi/psi torsion angles for *n*-6 degrees of freedom ("non-pivot" torsions) in a selected segment, and then solving the remaining 6 "pivot" degrees of freedom analytically to close the loop. To generate a protein backbone ensemble, different segment start and end points are sampled throughput the protein. Here, we generated near-native conformational ensembles with KIC (200 conformations with 0.9 Å average rmsd to the X-ray structure) using a modified protocol(*8*) compared to the published *de* *novo* loop reconstruction method(*7*). The ensemble generation protocol skips the low-resolution centroid stage and fixes the temperature at 1.2 *k*T. Further, to focus sampling on near-native conformations, non-pivot torsions are sampled within a vicinity of 3 degrees of the input value before each kinematic move, instead of sampling from the allowable Ramachandran space.

A second round of Rosetta sequence design was then applied across the ensembles to the side-chains surrounding the *de novo* built binding site in order to accommodate the motif residues and the small molecule target. Designing across a conformational ensemble, rather than a single backbone, can improve agreement between the geometry of the binding site built into the scaffold interface and the original geometry in the binding site template, and generates a diversity of predicted low-energy sequences.

The command line used to generate the ensemble with Rosetta r36129 was as follows:

loopmodel.linuxgccrelease -database minirosetta_database -loops:refine refine_kic -loops:max_kic_build_attempts 10000 -loops:input_pdb 1SVX_R134W103L78Y286__DE_19.pdb -loops:loop_file 1SVX_R134W103L78Y286__DE_19.loop -extra_res_fa 1SVX_R134W103L78Y286__DE_19_LG.fa.params -in:file:extra_res_cen 1SVX_R134W103L78Y286__DE_19_LG.cen.params -in:file:native 1SVX_R134W103L78Y286__DE_19.pdb -loops:kic_max_seglen 12 -loops:outer_cycles 1 -loops:refine_init_temp 1.2 -loops:refine_final_temp 1.2 -loops:vicinity_sampling -loops:vicinity_degree 3 -ex1 -ex2

A similar protocol for small molecule rigid body optimization and scaffold sequence design from section 1. 4 was applied to refine every member of the KIC ensemble. Additional rotamers were included (*via* the -ex3 -ex4 Rosetta command line flags), and the selection of residues to be redesigned, fixed, or modeled as wild-type was performed manually rather than selected by distance cutoffs (redesigned positions for the 1svx scaffold are shown in **Figure 2A**). Designs resulting from this step were further optimized for small molecule dependence on complex formation by requiring a ligand score < -6.0, ligand SASA > 0.7, ligand hydrogen bonds > 1, and unsatisfied buried ligand hydrogen bonds = 0. In addition to selecting single designs (see below), designing across a conformational ensemble, rather than a single backbone, produced a sequence library(*9*) (**Table S2**) to be assayed in *E. coli* for biosensor activity.

In addition to KIC ensemble design, we also used a second method for flexible backbone design: CoupledMoves(*10*). In contrast to KIC ensemble design, which first pre-generates an ensemble and then performs design on each member of the ensemble, CoupledMoves simultaneously moves the backbone and side chain of an amino acid (or rigid-body and conformer of the ligand) during design. In these simulations, we started from the model of design S3-1C (**Fig. 2A**) with FPP placed into the AR-MBP scaffold through matching with 4 motif residues (AR: L89, W114, R145; MBP: F133), followed by CoupledMoves design performed as described in (*10*), allowing the motif residues and the following additional residues to design: AR: Y81, H85, Y89, W112, M114, T115, H118, L119, K122, W123, F145, K147, I152, D155; MBP: E130, P133, F194, D197, K200, N201, K251. CoupledMoves design simulations were performed using Rosetta rotamer flags "-ex1 -ex2 -extrachi_cutoff 0 -use_input_sc" and were run at a constant temperature of 0.6 kT. The backbone moves used in these simulations were 3-residue "backrub" moves(*11*) centered on the amino acid being designed. Amino acid residues preferred in these simulations at key positions are shown in **Figure 2A**.

1.6 Design ranking and selection. We chose four computational designs for FPP sensors based on the AR/MBP (PDB 1svx) scaffold for testing:

S3-1A: Computational design ranked most highly by ligand burial

S3-1B: Computational design consensus sequence, based on most frequently selected amino acid residues in KIC ensemble design

S3-1C: Computational design with improved protein-ligand packing interactions

S3-1D: Computational design ranked most highly by predicted interaction affinity with the ligand

Models of these designs are shown below in two orientations. The orientation shown on the left was rotated 45° over the *x*-axis to attain the orientation shown on the right. AR: cyan, MBP: blue, FPP: magenta sticks, motif residues: sticks, designed residues: green spheres.

**_S3-1A_**

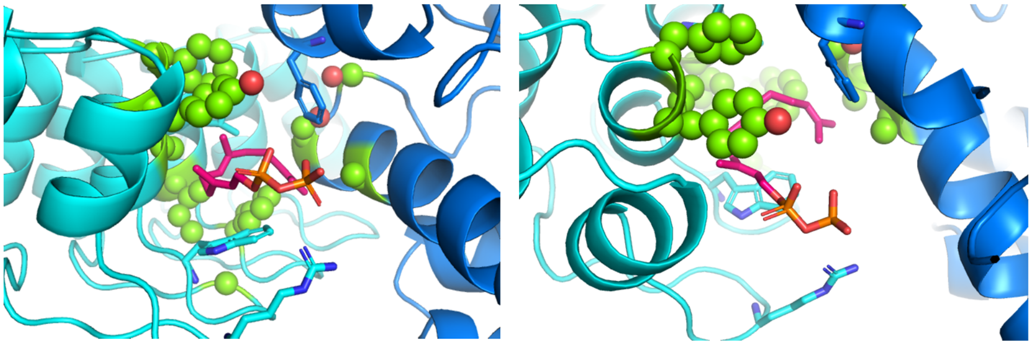

**_S3-1B_**

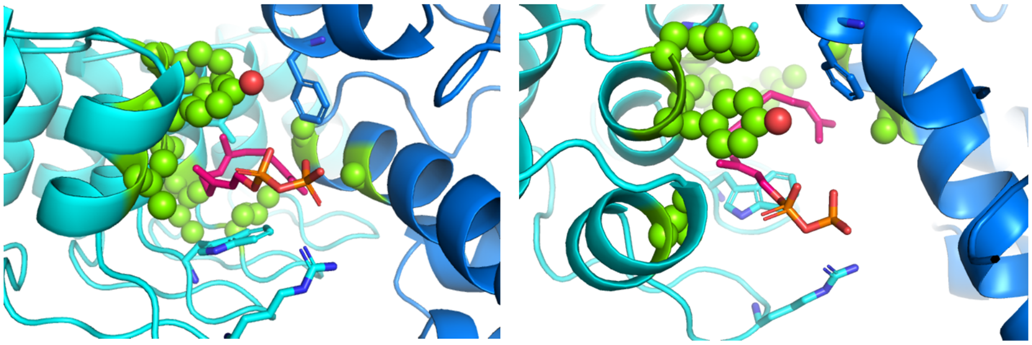

**_S3-1C_**

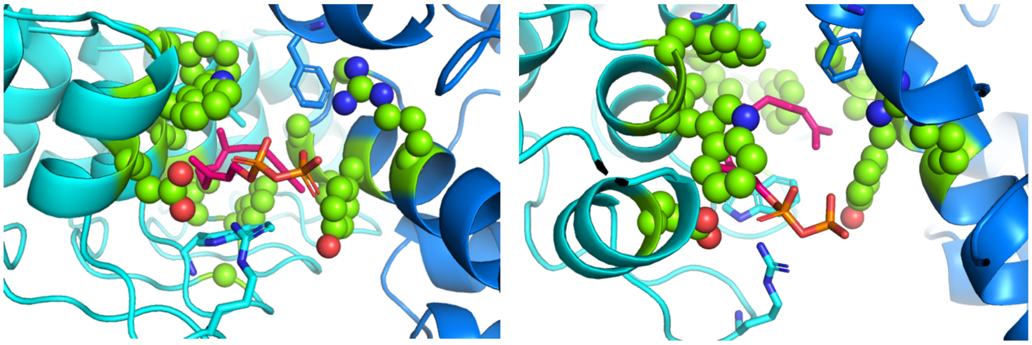

**_S3-1D_**

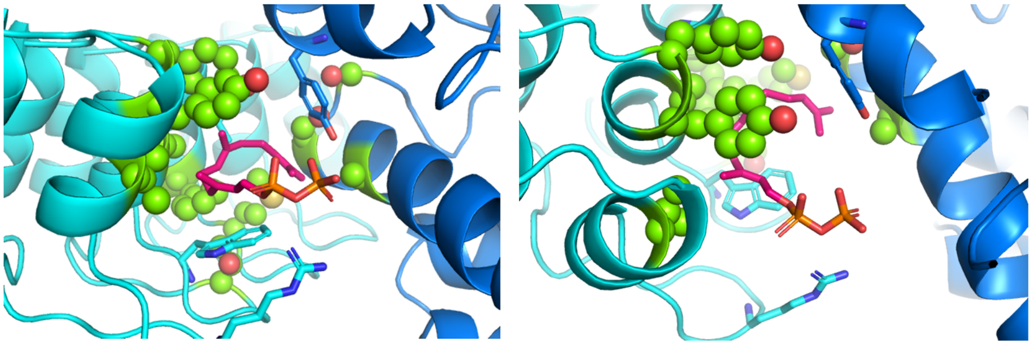

<u>1.7 Computational methods that resulted in designs on scaffolds 1 and 2</u>. Designs for scaffolds 1 and 2 resulted from the same overall protocol as described above, with the following modifications:

(i) We used an expanded protein-protein interface scaffold set consisting of 3462 heterodimers. Scaffolds were pre-relaxed using Rosetta FastRelax(*12*) with constraints to the starting coordinates for both backbone and side-chain atoms.

(ii) We used the following FPP binding site motifs:

| **Scaffold** | **Template PDB** | **Motif residues** |
|---|---|---|
| Scaffold 1 FRB/FKBP (3FAP.pdb) | 3bnx | F153, L184, W308, R314 |
| Scaffold 2 RapF/ComA (3ULQ.pdb) | 3bnx | F153, R314, Y315, F87 |

(ii) We did not use backbone ensembles to reshape the binding site environment around the motif residues built into the scaffold, but instead used Rosetta FastRelax to optimize both side chains and backbone in the binding site environment after the design step.

(iii) We selected designs for testing based on the top predicted ligand binding score.

For scaffold 1, we selected one design (S1-1A) that contained the following wild-type reversions: A28G, T36F, W46F, H48F, M59W, L193Y determined by visual inspection.

For scaffold 2, we selected 4 designs:

S2-1A: computationally designed sequence.

S2-1B: computationally designed with wild-type reversions: V21I, M404L and mutations of designed alanine residues: A28G, A32S, A59R.

S2-1C: as S2-1B with 2 additional alanine mutations to destabilize the wild-type protein-protein interaction: D23A, N418A.

S2-1D: as S2-1A with 2 additional mutations: W12F, A411M.

(iv) We used Rosetta version 54703 with the following command lines:

Building sites into scaffolds (matching):

```
match.linuxgccrelease -database ROSETTA_DATABASE -s INPUT_PDB
-match::lig_name LG1
-match::grid_boundary GRIDLIG_FILE
-match::scaffold_active_site_residues POSFILE
-match::geometric_constraint_file CST_FILE
-extra_res_fa FA_PARAMS_FILE
-output_matches_per_group 1
-match:consolidate_matches
-out:file:scorefile
-ex1 -ex2 -extrachi_cutoff 0 -use_input_sc
-euclid_bin_size 1.5 -euler_bin_size 15 -bump_tolerance 0.5
-out::path OUTDIR -match:output_format PDB
```

Design step:

```
enzyme_design.linuxgccrelease -database ROSETTA_DATABASE -s INPUT_PDB
-in:file:fullatom -out:file:o designs.score -extra_res_fa FA_PARAMS_FILE
-enzdes:cstfile CONSTRAINTS_FILE -overwrite -out:pdb_gz -nstruct 10
-enzdes:cst_design -enzdes:detect_design_interface
-enzdes:cut1 6.0 -enzdes:cut2 8.0 -enzdes:cut3 10.0 -enzdes:cut4 12.0
-enzdes:cst_opt -enzdes:cst_min -enzdes:bb_min -enzdes:chi_min
-enzdes:design_min_cycles 3 -ex1 -ex2 -extrachi_cutoff 0 -use_input_sc
-enzdes:start_from_random_rb_conf -enzdes:final_repack_without_ligand
-score:weights talaris2013_cst.wts
```

Fastrelax of designs:

```
relax.linuxgccrelease -database ROSETTA_DATABASE -s INPUT_PDB
-extra_res_fa FA_PARAMS_FILE -out:pdb_gz -out::path OUTDIR
-ignore_zero_occupancy false -relax:fast
-relax:constrain_relax_to_start_coords
-ex1 -ex2 -extrachi_cutoff 0 -use_input_sc
-score:weights talaris2013_cst.wts
-preserve_header -nstruct 10
```

Rescoring relaxed designs:

```
enzyme_design.linuxgccrelease -database ROSETTA_DATABASE -s INPUT_PDB
-in:file:fullatom -out:file:o designs.score -extra_res_fa FA_PARAMS_FILE
-enzdes:cstfile CONSTRAINTS_FILE -overwrite -out:pdb_gz
-enzdes:detect_design_interface -enzdes:cut1 0.0 -enzdes:cut2 0.0
-enzdes:cut3 10.0 -enzdes:cut4 12.0 -ex1 -ex2 -extrachi_cutoff 0
-use_input_sc -enzdes:no_unconstrained_repack -enzdes:lig_packer_weight 1.8
-enzdes:final_repack_without_ligand -score:weights talaris2013_cst.wts
```

1.8 Computational methods to further stabilize sensor S3-2D. Using the crystal structure for S3-2D as the input structure, we used the RosettaScripts platform to apply two successive cycles of CoupledMoves(*10*) to improve FPP-protein interactions and the binding pocket stability. The input structure was pre-relaxed using Rosetta FastRelax with constraints to the starting coordinates for both backbone and side-chain atoms. In each cycle of CoupledMoves, we chose a different group of neighboring residues in the binding pocket to be designed. The second cycle received the lowest-energy structure from the previous cycle as the input structure. Results were ranked according to total energy. Structures and sequences of top-ranked designs were manually inspected to choose which designs to experimentally test.

| **CoupledMoves cycle** | **Designed protein** | **Designable residues** |
|---|---|---|
| 1 | AR of S3-2D (AR-2.7) | 80-82, 84-86, 88-90, 118-122, 141-145, 150-155 |
| 2 | MBP of S3-2D (MBP-2.5) | 132-134, 196-202, 250-252 |

Rosetta commands (Rosetta 3.8, commit f3a3f038d9419c0d79a6030ea4dc16f62668d69f):

FastRelax of crystal structure:
relax.linuxgccrelease -database ROSETTA_DATABASE -in:file:s INPUT_PDB -in:file:fullatom -relax:constrain_relax_to_start_coords -extra_res_fa FA_PARAMS_FILE

Design flags:
```
-in:file:s INPUT_PDB
-extra_res_fa FA_PARAMS_FILE
-packing
	-ex1
	-ex1aro
	-extrachi_cutoff 0
	-ex2
-number_ligands 1
-coupled_moves
	-ntrials 10000
	-initial_repack false
	-min_pack true
```

```
	-ligand_mode true
	-ligand_weight 2.0
-nstruct 20
```

<u>RosettaScript to run CoupledMoves:</u>

```
<ROSETTASCRIPTS>
  <SCOREFXNS>
    <ScoreFunction name="ref15"/>
  </SCOREFXNS>

  <TASKOPERATIONS>
    <ReadResfile name="AR_resfile" filename=RESFILE_NAME_1/>
    <ReadResfile name="MBP_resfile" filename= RESFILE_NAME_2/>
  </TASKOPERATIONS>

  <MOVERS>
    <CoupledMovesProtocol name="coupled_moves_AR" task_operations="AR_resfile"/>
    <CoupledMovesProtocol name="coupled_moves_MBP" task_operations="MBP_resfile"/>
    <DumpPdb name="dump" fname="dump" tag_time="1"/>
    <FastRelax name="fastrelax" repeats="5"/>
  </MOVERS>

  <PROTOCOLS>
    <Add mover_name="coupled_moves_AR"/>
    <Add mover_name="dump"/>
    <Add mover_name="coupled_moves_MBP"/>
  </PROTOCOLS>
</ROSETTASCRIPTS>
```

2. Complementation assay with split murine dihydrofolate reductase (mDHFR).

2.1 Constructs and strains. We tested sensor function in *E. coli* strain DH10B using complementation of split mDHFR(*13*). The constructs used in this assay are listed in **Appendix 1** and the gene sequences are listed in **Appendix 4**. For all MBP genes tested in the mDHFR assay, we deleted the sequence corresponding to the last alpha helix in the structure (residues 354-370). This modification was made to bring the two sensor protein termini closer together for facile complementation of the attached split mDHFR constructs upon small-molecule mediated dimerization. A detailed protocol of the assay is given below.

2.2 Liquid culture assay. Fresh preparations were made of filter-sterilized 0.2 M isopropyl β-D-1-thiogalactopyranoside (IPTG) in sterile deionized water, 20% L-arabinose in sterile deionized water, 5 mg/ml trimethoprim (TMP) (Sigma) solution in methanol, and autoclaved M9 medium (0.1 mM $CaCl_2$, 1X M9 salts, 2 mM $MgSO_4$, 0.4% glucose, 0.2% Casaminoacids, 10 mg/ml leucine, 10 mg/ml isoleucine, 5 mg/ml thiamine, all filter-sterilized). One 4 ml overnight culture was grown at 37 °C and 225 rpm for each strain from freshly transformed plates in M9 medium with 37 μg/ml chloramphenicol and 50 μg/ml spectinomycin. To prepare media for the assay, the following was added to 49 ml M9 medium: 37 μg/ml chloramphenicol and 50 μg/ml spectinomycin, 60 μM IPTG (unless otherwise noted), 0.4% L-arabinose, and 6 μg/ml TMP (unless otherwise noted). All reagents were thawed completely, vortexed, and centrifuged prior to addition to the master mix to avoid precipitants. All ingredients were gently mixed in a Falcon tube, and then 4 ml of the master mix was aliquoted into new culture tubes. Each overnight culture was gently vortexed, checked for complete resolubilization of any chunks, diluted 1 to 100 in fresh medium, and poured into a reagent reservoir. Using a multichannel pipette, 200 µl of the diluted cultures were transferred to the wells of a transparent, flat-bottom 96-well plate (Costar), 8 wells at a time. A final concentration of 5 mM mevalonate (Sigma) (unless otherwise noted) was added to every other column via 1 to 1000 dilution of a 1 M stock solution in Millipore water. To prevent evaporation of medium, 50 µl mineral oil was gently pipetted on top of each well's contents. The plate was centrifuged for 30 seconds to pop bubbles and any remaining bubbles were popped manually with a pipette tip. The plate was placed in a shaker and grown at 35 °C and 200 rpm for 48 hours. After 48 hours of growth, cultures were mixed gently using a multichannel pipette set to 50 µl, avoiding disturbance of the mineral oil layer. Bubbles were removed using a Bunsen burner flame and/or pipette tips. Cell culture optical density (OD) was read over the whole plate, without a plate cap, at 600 nm using a Tecan Safire 2 microplate reader.

2.3 Library screening and saturation mutagenesis

2.3.1 Library construction. The computationally designed library (**Table S2**) was constructed using gene assembly mutagenesis(*14*). Briefly, multiple oligonucleotides were synthesized to assemble the design regions of AR and MBP, and 14 and 16 oligonucleotides were generated to cover the protein-protein interface of AR and MBP respectively. Computationally predicted mutations within the regions were introduced into the designed oligonucleotides using degenerate codons that code for a list of desired amino acids (**Table S2**). These synthesized oligonucleotides were combined and assembled using a two-step PCR to form the library of AR and MBP fragments, and both sets of the fragments were cloned sequentially into pCDFDuet to obtain the final library constructs.

2.3.2 Library screening. The library constructs were electroporated into DH10B *E. coli* and plated onto M9 minimal medium plates containing 100 μM IPTG, 0.4 % L-arabinose, 5 mM mevalonate and 6 μg/ml TMP. The plates were cultured for 10 days at room temperature. Clones that survived under TMP selection were potential active candidates that could form functional mDHFR and were enriched in this screening. A total of 3,072 clones were selected from this step and subjected to two-step confirmation screening – 1) colony-printing assay and 2) liquid culture assay.

In the colony-printing assay, freshly grown overnight cultures of each selected clone were 100X diluted and spotted onto two sets of M9 minimal medium plates, one with 5 mM mevalonate and one without. Both sets of the plates contained 100 μM IPTG, 0.4 % L-arabinose and 6 μg/ml TMP. The plates were cultured for 10 days at room temperature. Clones that showed increased growth (as indicated by the size of the colonies) with the addition of mevalonate after 10 days were selected and individually tested using the liquid culture assay as described above. The same conditions were kept in the liquid culture screening. 36 out of 3,072 candidates were selected from the colony-printing screen and 27 out of these 36 candidates were positive in the liquid culture validation.

2.3.3 Saturation mutagenesis. We performed two rounds of iterative saturation mutagenesis (ISM, **Table S3**). For each round, 11 positions (AR: position 85, 112, 118, 119, 122, 123, 152, 155; MBP: 194, 197, 251) around the designed binding site were allowed to mutate to any of 20 amino acids coded by NNK degenerate codon starting from S3-2A. Only one of 11 positions was allowed to mutate at a time (a.k.a. single-site saturation). The best clone from the first round of ISM (S3-2B) was selected as the starting sequence for the second round of ISM. A total of 384 clones and 480 clones were screened in the first and second round of ISM respectively. The top clones for each round were screened using the liquid culture assay as described above using 80 μM IPTG, 0.4 % L-arabinose, 5 mM mevalonate and 6 μg/ml TMP.

3. Protein purification

3.1 Constructs. We expressed and purified several S3 sensor proteins for further characterization. The expression plasmids are listed in **Appendix 2** and the gene sequences are listed in **Appendix 4**. For all MBP genes in expression plasmids, we included the sequence corresponding to the last alpha helix in the structure (residues 354-370).

3.2 Purification of ankyrin repeat (AR) proteins. Overnight cultures of BL21(DE3) pLysS cells (Novagen) harboring the pET47-6XHis-AR plasmid were grown with 37 μg/ml chloramphenicol and 50 μg/ml kanamycin for 14-18 hours at 37 °C and 225 rpm in lysogeny broth (LB) medium. Overnight cultures were diluted 1 to 50 in LB and grown at 37 °C and 225 rpm until the $OD_{600}$ reached approximately 0.6 (about two hours). Cultures were induced with 0.5 mM IPTG and further grown at 16 °C overnight. Cells were pelleted by centrifugation at 6,000 *g* for 20 minutes. Pellets from 1 L cell cultures were stored at -80 °C. Frozen cell pellets were thawed on ice for 20 minutes and resuspended by vortexing in 40 ml lysis buffer (25 mM Tris, 500 mM NaCl, pH 8.0). The following components were added to the cell resuspension: 5 mM $MgCl_2$, 1 mM $MnCl_2$, 100 μM $CaCl_2$, one protease inhibitor tablet (Thermo Fisher), and 5 μL 2000 units/ml DNaseI per ml lysis buffer. Cells were lysed by the addition of 10X BugBuster (Thermo Fisher) to a final concentration of 1X, mixed gently, and stored at room temperature for 5 minutes. Cell lysate was centrifuged at 27,000 *g* for 30 minutes at 4 °C. The supernatant was decanted into a fresh tube, and imidazole was added to 20 mM. The soluble lysate was further incubated on ice for 1 hour to allow any remaining precipitate to form, then centrifuged at 3,700 *g* for 10 minutes at 4 °C. The supernatant was transferred to a fresh tube, mixed with 3 ml Ni-NTA Slurry (Thermo Fisher) per 40 ml lysate, and mixed at 4 °C on a nutator for 1 hour. The lysate-slurry mix was centrifuged at 500 *g* for 5 minutes at 4 °C and the supernatant was immediately discarded. The pelleted resin was resuspended in two resin-volumes of cold wash buffer (25 mM Tris, 500 mM NaCl, 20 mM imidazole, pH 8.0) and applied to a $Ni^{2+}$-NTA gravity column, then the column was washed with two column volumes of cold wash buffer. Protein was eluted with 1.5 column volumes in 25 mM Tris, 500 mM NaCl, 250 mM imidazole, pH 8.0. The eluate was desalted into 25 mM Tris, 50 mM NaCl, pH 8.0 using an AKTA FPLC with a Hiprep 26/10 desalting column, then ion-exchanged using a Hitrap Q HP column into 25 mM Tris, 500 mM NaCl, pH 8.0 The resulting protein solution was concentrated down to 5 ml using 3 KDa MW cutoff spin concentrators (Sartorius), applied to a size-exclusion HiLoad 16/60 Superdex 75 column, and eluted in 50 mM Tris, 150 mM NaCl, pH 8.0. The eluate was concentrated to 1.5 ml and dialyzed three times against 1 L 25 mM Tris, 150 mM NaCl, pH 8.0, for two hours, overnight, and two hours, respectively. The final protein concentration was determined in the denatured state in 6 M guanidinium hydrochloride, 20 mM potassium phosphate solution and the purity was confirmed by Coomassie-stained SDS-PAGE to be > 95%.

3.3 Purification of maltose binding proteins (MBP). Methods for the growth and lysis of BL21(DE3) pLysS cells with pET28-MBP plasmids were similar to those described above for AR proteins, with the following modifications: (1) overnight cultures were subcultured with 0.2% glucose, and (2) lysis buffer was composed of 25 mM Tris, 150 mM NaCl, pH 8.0. Isolation of the soluble lysate was followed by incubation on ice for 1 hour, then centrifugation at 3,700 *g* for 10 minutes at 4 °C. The supernatant was decanted into a fresh tube, applied to an MBP affinity column (MBPtrap HP column) using an AKTA FPLC, and eluted in 50 mM Tris, 150 mM NaCl, 30 mM maltose, pH 8.0. The volume of the protein solution was reduced to 5 ml using 10 KDa MW cutoff spin concentrators (Millipore) and applied to a size-exclusion HiLoad 16/60 Superdex 75 column, and eluted in 50 mM Tris, 150 mM NaCl, pH 8.0. The size exclusion eluate volume was concentrated down to 1.5 ml and dialyzed three times against 1 L 25 mM Tris, 150 mM NaCl, pH 8.0, for two hours, overnight, and two hours, respectively. The final protein concentration was determined in the denatured state in 6 M guanidinium hydrochloride, 20 mM potassium phosphate solution and the purity was confirmed by Coomassie-stained SDS-PAGE to be >98%.

4. *In vitro* binding assays using bio-layer interferometry (BLI)

4.1 Constructs. We purified several S3 sensor protein pairs to determine their *in vitro* binding affinity in the presence and absence of FPP by BLI. In our BLI experiments, one protein is tethered to an optically transparent biosensor tip by a biotin-streptavidin interaction, and the other protein is present as the analyte in solution in the microplate. BLI expression plasmids are listed in **Appendix 2** and the gene sequences are listed in **Appendix 4**. Constructs for BLI experiments included the sequence corresponding to the last alpha helix in MBP (residues 354-370).

4.2 Biotinylation of avi-MBP. Avi-tagged MBP (**Appendix 2**) was prepared at 66 μM in 10 mM Tris buffer, pH 8.0. The BirA-500 biotin-protein ligase kit from Avidity was used as per the manufacturer's instructions, with a final avi-MBP concentration of 40 μM and 2.5 μg BirA enzyme per 10 nmol protein. The reaction was performed at 30 °C for 45 minutes, and then buffer-exchanged by spin concentration and 50-fold dilution three times in fresh 25 mM Tris, 150 mM NaCl buffer, pH 8.0, to remove excess biotin and reaction components. Biotinylated avi-MBP was diluted to 40 μM and stored at 4 °C for use in BLI measurements.

4.3 BLI. Affinity measurements between avi-tagged MBP, AR and FPP were performed at room temperature using an Octet RED96 system and streptavidin (SA)-coated biosensor tips (Pall ForteBio). Avi-MBP was diluted to 400 nM in HBS-P buffer (0.01 M HEPES pH 7.4, 0.15 M NaCl, 0.005% v/v Surfactant P20) to be used as the antigen. Antigen-bound SA-tips were washed, exposed to the analyte solutions during an association period, and then allowed to unbind from the analyte during a dissociation period. For experiments including FPP, the analyte solution was composed of 0-24 μM AR and 200 μM FPP. For experiments measuring the affinity of MBP for AR without FPP, the analyte solution was composed of 0-500 μM AR. The binding protocol was as follows: rinse tips in HBS-P buffer, 60 seconds; load tips with antigen, 300 seconds; establish baseline by rinsing tips in HBS-P buffer, 60 seconds; association with analyte, 60 seconds; dissociation in baseline wells, 600 seconds. Raw data was fit to 1:1 binding curves in Octet Data Analysis HT software using curve fitting kinetic analysis with local fitting. The theoretical equilibrium binding signal response data (R equilibrium) were normalized by the steady-state group maximum response ($R_{max}$) values, and the steady state affinity was determined using the Hill equation,

$$\theta = \frac{1}{\frac{K_A}{[L]}+1} ,$$

where θ is the fraction of MBP that is bound to AR, $K_A$ is the AR concentration at which half the MBP are occupied; and $[L]$ is the concentration of unbound AR. Non-cooperative binding kinetics are assumed. All binding curves fit the BLI data with $R^2$>0.90.

5. Cell-free transcription-translation (TxTl) assay

5.1 Constructs. Plasmids for TxTl experiments were constructed using standard BsaI/BsmBI Golden Gate cloning methods as described in Engler *et al.*(*15*) using our group's adaptation of the MoClo Yeast Tool Kit strategy as described in Lee *et al.*(*16*). For all MBP genes in TxTl plasmids, we included the sequence corresponding to the last alpha helix in the structure (residues 354-370). A comprehensive list of plasmids is available in **Appendix 3**.

5.2 TxTl protocol. Separate 30 µl TxTl reactions were prepared for each plasmid (expressing MBP or AR variants) as described in Sun *et al.*(*17*) and incubated for 8-16 hours at 29 °C. Each reaction also included 0.2 nM of the TxTl11 plasmid for RNA polymerase expression (**Appendix 3**) and 1 mM IPTG. The cell extract was prepared from Rosetta 2 cells (EMD Millipore). All TxTl reactions were mixed 1 to 1 with phosphate buffered saline/1% bovine serum albumin (PBS/BSA solution) for a total protein solution volume of 60 µl. A master mix was prepared by mixing 1 volume TxTl extract expressing an MBP variant, 1 volume TxTl extract expressing an AR variant, and 2 volumes PBS/BSA solution. Master mix aliquots of 27 µl were mixed with 3 µl farnesyl pyrophosphate (FPP) (Sigma) stock solutions in water with concentrations ranging from 1 µM to 2 mM. These 30 µl solutions were distributed in 9.5 µl aliquots in a white opaque 384 well plate for luminescence measurements or a black opaque 384 well plate for fluorescence measurements, centrifuged for 30 seconds to remove bubbles, and stored at room temperature for 10 minutes. For luminescence measurements, 1 ml plus 10 µl per well of Promega NanoGlo buffer was thawed on ice and mixed with a 50-fold dilution of furimazine substrate to make NanoGlo reagent, as described in Dixon *et al.*(*18*). The reagent was distributed in 9.5 µl aliquots to all wells containing

9.5 μl protein/FPP sample. After ten minutes, a SpectraMax L luminometer (Molecular Devices) was used to take luminescence readings using analog detection and injection, following priming of the injection lines with 1 ml NanoGlo reagent. Data were collected with the following machine settings: integration time, 1 s; PMT setting, photon counting; automix setting, classic with 5 s mix duration and 30 mm/s mix speed; M-injection, with injector volume 10 μl, post-injection delay 1 s; injection speed 320 μl/s, and no shaking after injection; and dark adapt set to off. For fluorescence measurements, a Tecan Safire 2 microplate reader was used to read ddGFP complementation from the top of the plate at excitation/emission wavelengths of 380 nm and 508 nm, with the gain set to 100.

5.3 Data fitting for *in vitro* FPP titration measurements. Raw background-subtracted luminescence and fluorescence data from TxTl assay measurements were averaged for each FPP concentration and fit to a modified three-parameter logistic nonlinear regression model of the Hill equation,

$$\hat{Y} = a + \frac{b-a}{1+\frac{c}{x}},$$

where $\hat{Y}$ is the expected response at FPP concentration $x$, $a$ is the minimum response when no FPP is present, $b$ is the maximum response, and $c$ is the FPP concentration at which half the protein sensors are bound, allowing for a fluorescent or luminescent output. Data and fit were normalized by subtracting the calculated minimum response and dividing by the difference of the calculated maximum and minimum responses.

6. Crystallography.

6.1 Constructs. The plasmids used to express S3-2D sensor proteins for crystallography are listed in **Appendix 2**. Their sequences are listed in **Appendix 4**. Methods for protein expression and purification are described in Section 3.

6.2 Crystallization. AR and MBP were each prepared at 340 μM and mixed 1:1 to obtain each protein at 170 μM in solution. FPP was prepared at 170 mM in 70% ethanol and diluted 100-fold in the protein solution for a final concentration of 1.7 mM FPP. We carried out initial crystallization trials in 15-well hanging drop format using EasyXtal crystallization plates (Qiagen) and a crystallization screen that was designed to explore the chemical space around the crystallization conditions reported by Binz *et al.* (*19*) for the co-crystal structure of their AR-MBP complex (PDBID:1SVX). Crystallization drops were prepared by mixing 1 μl of protein solution with 1 μl of the mother liquor, and sealing the drop inside a reservoir containing an additional 500 μl of the mother liquor solution. Thin, plate-like crystals were obtained in many of the conditions tested, with the largest single crystals growing from drops prepared using mother liquor containing 0.1 M Tris Buffer pH 8.7, 0.1 M sodium chloride, and 32% PEG-6000. The crystals could not be obtained without the addition of the FPP ligand, and we used SDS-PAGE to confirm that both proteins required to form the heterodimer were present in the crystals (after harvesting and washing in mother liquor without protein).

6.3 X-ray data collection and processing. Prior to X-ray data collection, crystals were dehydrated overnight in a solution containing 0.1 M Tris Buffer pH 8.7, 0.1 M sodium chloride, and 45%

PEG-6000. Next, the crystals were soaked in a solution containing 0.1 M Tris Buffer pH 8.7, 0.1 M sodium chloride, 42% PEG-6000, 7% ethanol, and 18 mM FPP for 30 minutes before they were harvested and flash-cooled in liquid nitrogen. The soaking steps were essential to improve the quality of the observed X-ray diffraction patterns, and to ensure adequate occupancy of the FPP ligand in the resulting electron density maps. No cryoprotectant was necessary due to the high concentration of PEG-6000 in the soaking solutions.

We collected single-crystal X-ray diffraction data on beamline 8.3.1 at the Advanced Light Source. The beamline was equipped with an ADSC Quantum 315r CCD detector, and the crystals were maintained at a cryogenic temperature (100 K) throughout the course of data collection.

We processed the X-ray data using the Xia2 system(*20*), which performed indexing, integration, and scaling with XDS and XSCALE(*21*), followed by merging with Pointless(*22*). A resolution cutoff (2.20 Å) was taken where the completeness of the data fell to a value of approximately 90%. Although other metrics of data quality (such as CC1/2 and <I/σI>) suggest that a more aggressive resolution cutoff would be acceptable, we were limited by physical constraints on the experimental geometry. Specifically, the plate-like morphology of the crystals resulted in the long axis of the unit cell always being perpendicular to the crystal rotation (phi) axis, which constrained the minimum sample-to-detector distance and the maximum Bragg angle that could be recorded on the detector without producing overlap between individual reflections. Further information regarding data collection and processing is presented in **Table S4**. The reduced diffraction data were analyzed with phenix.xtriage (http://www.ccp4.ac.uk/newsletters/newsletter43/articles/PHZ_RWGK_PDA.pdf) to check for crystal pathologies , which revealed the presence of pseudomerohedral twinning based on the results of the L-test (<|L|>=0.399, <L2>=0.225)(*23*).

6.4 Structure determination. We obtained initial phase information for calculation of electron density maps by molecular replacement using the program Phaser(*24*), as implemented in the PHENIX suite(*25*). We searched for each component of the heterodimer independently, using separate models of AR (PDBID: 1SVX, chain A) and the "closed" (ligand-bound) form of MBP (PDBID: 1FDQ). We explicitly did not use the design model for molecular replacement to avoid the introduction of model bias. Two complete copies of the heterodimer were found in the unit cell, consistent with an analysis of Matthews probabilities for the observed unit cell and molecular weight of the heterodimer(*26, 27*).

Next, we attempted to rebuild the missing or incorrect parts of the molecular replacement solution using the electron-density maps calculated from model phases, however, the presence of twinning compromised the quality of the initial maps. To improve the interpretability of the electron density maps, we carried out a round of atomic refinement using phenix.refine(*28*) in which we also refined the twin fraction (twin operator = *h,-k,-l*; refined twin fraction, $\boldsymbol{\alpha}$ = 0.49). The twin refinement improved the map quality substantially, allowing us to rebuild the missing or incorrect parts of the structure. Additional iterative steps of manual model rebuilding and atomic refinement were performed, and during this process we found evidence for several small molecule ligands in both $2mF_o\text{-}DF_c$ and $mF_o\text{-}DF_c$ electron density maps. Specifically, we identified an FPP molecule occupying one of the computationally-designed binding sites, as well as two maltose disaccharides (one bound to each copy of MBP). The presence of merohedral twinning can present challenges for map interpretation because the structure factor detwinning process can exacerbate the effects of model phase bias(*29*). Consequently, we took extra care with the placement of the FPP ligand into electron density features around the designed binding site. The initial placement of the ligand was motivated by the observation of several strong electron density peaks near a key lysine residue and in the hydrophobic binding cavity, which could be attributed to the pyrophosphate group and aliphatic tail of FPP, respectively. Refinement of the structure with FPP in the binding site resulted in a rearrangement of side chains in contact with the ligand, providing additional evidence that this binding site is occupied. Specifically, Trp114 of AR becomes ordered upon FPP binding, and Y197 in MBP rotates and displaces a small network of two neighboring water molecules (**Figure S11**). After adding the ligands, we performed additional refinement of atomic positions, atomic displacement parameters, and occupancies using non-crystallographic symmetry (NCS) and secondary structure restraints, a riding hydrogen model, and automatic weight optimization. The refined atomic B-factors of the FPP ligand are similar to those of the surrounding protein atoms. After refining the model to convergence, we further verified the presence of the FPP ligand in the model by repeating our final refinement starting from atomic coordinates without the ligand atoms. This "omit refinement" produced $2mF_o\text{-}DF_c$ and $mF_o\text{-}DF_c$ omit maps and allowed us to assess the effect of including the ligand. The omit maps showed density that was highly similar to the original maps that permitted initial placement of FPP (**Figure S10**). The absence of FPP in the second ligand binding site could be the result of distortions to the heterodimeric interface caused by crystallization, or by the dehydration procedure that was required to improve the diffraction resolution. All model building was performed using Coot(*30*) and refinement steps were performed with phenix.refine (v1.13-2998) within the PHENIX suite(*25, 28*). Restraints for the maltose and FPP ligands were calculated using phenix.elbow(*31*). The final model coordinates were deposited in the Protein Data Bank (PDB(*32*)) under accession code 6OB5. Further information regarding model building and refinement is presented in **Table S4**.

**Figure S1**

### Scaffold 1:

#### FKBP-12

```
            1                  20                  40                  60
            |                  |                   |                   |
FKBP12-WT   GVQVETISPGDGRTFPKRGQTCVVHYTGMLEDGKKFDSSRDRNKPFKFMLGKQEVIRGWEEGVAQMSVGQRAKLTISPD
FKBP12-1.1  GVQVETISPGDGRTFPKRGQTCVVHITGMLEDGKKFISSRDRNKPFKFMLGKQRVLRGWEEGVAQMSVGQRAKLTISPD

            80                 100
            |                   |
FKBP12-WT   YAYGATGHPGIIPPHATLVFDVELLKLE
FKBP12-1.1  YAFGATGHPGIIPPHATLVFDVELLKLE
```

#### FRB

```
        108                127                 147                 167
         |                  |                   |                   |
FRB-WT   VAILWHEMWHEGLEEASRLYFGERNVKGMFEVLEPLHAMMERGPQTLKETSFNQAYGRDLMEAQEWCRKYMKSGNVKDL
FRB-1.1  VAILWHEMWHEGAEEAARLYRGERNVKGMFEVLEPLHAMMERGPQTLKETSFNQAYGRDLMEAQEWCRKYMKSGNVKDL

        187
         |
FRB-WT   TQAWDLYYHVFRRIS
FRB-1.1  WQAMLLYAHVRDRIS
```

### Scaffold 2:

#### RapF

```
          1                  20                  40                  60
          |                  |                   |                   |
RapF-WT   SSSSIGEKINEWYMYIRRFSIPDAEYLRREIKQELDQMEEDQDLHLYYSLMEFRHNLMLEYLEPLEKMRIEEQPRLSDL
RapF-1.1  SSSSIGEKINEWYMYIRRFSIPDAAYLAFEIAQELDQMEEDQDLHLYYSLMLFRAYLMAEYLEPLEKMRIEEQPRLSDL
RapF-1.2  SSSSIGEKINEWYMYIRRFSVPDAAYLGFEISQELDQMEEDQDLHLYYSLMLFRAYLMREYLEPLEKMRIEEQPRLSDL
RapF-1.3  SSSSIGEKINEWYMYIRRFSVPAAAYLGFEISQELDQMEEDQDLHLYYSLMLFRAYLMREYLEPLEKMRIEEQPRLSDL
RapF-1.4  SSSSIGEKINEFYMYIRRFSIPDAAYLAFEIAQELDQMEEDQDLHLYYSLMLFRAYLMAEYLEPLEKMRIEEQPRLSDL

          80
          |
RapF-WT   LLEIDKK
RapF-1.1  LLEIDKK
RapF-1.2  LLEIDKK
RapF-1.3  LLEIDKK
RapF-1.4  LLEIDKK
```

#### ComA

```
          377                396                 416
          |                  |                   |
ComA-WT   VLTPRECLILQEVEKGFTNQEIADALHLSKRSIEYSLTSIFNKLNVGSRTEAVLIAKS
ComA-1.1  VLTPRECLILQEVEKGFTNQEIADALHLRKSAIEASLTSIFNKLNVGSRTEAVLIAKS
ComA-1.2  VLTPRECLILQEVEKGFTNQEIADALHMRKSAIEASLTSIFNKLNVGSRTEAVLIAKS
ComA-1.3  VLTPRECLILQEVEKGFTNQEIADALHMRKSAIEASLTSIFAKLNVGSRTEAVLIAKS
ComA-1.4  VLTPRECLILQEVEKGFTNQEIADALHLRKSAIEMSLTSIFNKLNVGSRTEAVLIAKS
```

### Scaffold 3:

#### MBP

```
        1                 20                  40                  60
        |                  |                   |                   |
MBP-WT  KIEEGKLVIWINGDKGYNGLAEVGKKFEKDTGIKVTVEHPDKLEEKFPQVAATGDGPDIIFWAHDRFGGYAQSGLLAEI
MBP-1.1 KIEEGKLVIWINGDKGYNGLAEVGKKFEKDTGIKVTVEHPDKLEEKFPQVAATGDGPDIIFWAHDRFGGYAQSGLLAEI
MBP-1.2 KIEEGKLVIWINGDKGYNGLAEVGKKFEKDTGIKVTVEHPDKLEEKFPQVAATGDGPDIIFWAHDRFGGYAQSGLLAEI
MBP-1.3 KIEEGKLVIWINGDKGYNGLAEVGKKFEKDTGIKVTVEHPDKLEEKFPQVAATGDGPDIIFWAHDRFGGYAQSGLLAEI
MBP-1.4 KIEEGKLVIWINGDKGYNGLAEVGKKFEKDTGIKVTVEHPDKLEEKFPQVAATGDGPDIIFWAHDRFGGYAQSGLLAEI
MBP-2.5 KIEEGKLVIWINGDKGYNGLAEVGKKFEKDTGIKVTVEHPDKLEEKFPQVAATGDGPDIIFWAHDRFGGYAQSGLLAEI
MBP-3.6 KIEEGKLVIWINGDKGYNGLAEVGKKFEKDTGIKVTVEHPDKLEEKFPQVAATGDGPDIIFWAHDRFGGYAQSGLLAEI

        80                 100                 120                 140
        |                   |                   |                   |
MBP-WT  TPDKAFQDKLYPFTWDAVRYNGKLIAYPIAVEALSLIYNKDLLPNPPKTWEEIPALDKELKAKGKSALMFNLQEPYFTW
MBP-1.1 TPDKAFQDKLYPFTWDAVRYNGKLIAYPIAVEALSLIYNKDLLPNPPKTWEEIFALDKELKAKGKSALMFNLQEPYFTW
MBP-1.2 TPDKAFQDKLYPFTWDAVRYNGKLIAYPIAVEALSLIYNKDLLPNPPKTWEEIFALDKELKAKGKSALMFNLQEPYFTW
MBP-1.3 TPDKAFQDKLYPFTWDAVRYNGKLIAYPIAVEALSLIYNKDLLPNPPKTWEEIFALDKELKAKGKSALMFNLQEPYFTW
MBP-1.4 TPDKAFQDKLYPFTWDAVRYNGKLIAYPIAVEALSLIYNKDLLPNPPKTWEEIFALDKELKAKGKSALMFNLQEPYFTW
MBP-2.5 TPDKAFQDKLYPFTWDAVRYNGKLIAYPIAVEALSLIYNKDLLPNPPKTWEEIFALDKELKAKGKSALMFNLQEPYFTW
MBP-3.6 TPDKAFQDKLYPFTWDAVRYNGKLIAYPIAVEALSLIYNKDLLPNPPKTWEEIFALDKELKAKGKSALMFNLQEPYFTW

         160                 180                 200                 220
         |                   |                   |                   |
MBP-WT  PLIAADGGYAFKYENGKYDIKDVGVDNAGAKAGLTFLVDLIKNKHMNADTDYSIAEAAFNKGETAMTINGPWAWSNIDT
MBP-1.1 PLIAADGGYAFKYENGKYDIKDVGVDNAGAKAGLTFLVALIAAKAMNADTDYSIAEAAFNKGETAMTINGPWAWSNIDT
MBP-1.2 PLIAADGGYAFKYENGKYDIKDVGVDNAGAKAGLTFLVALIKAKHMNADTDYSIAEAAFNKGETAMTINGPWAWSNIDT
MBP-1.3 PLIAADGGYAFKYENGKYDIKDVGVDNAGAKAGLTRLVYLIAAKAMNADTDYSIAEAAFNKGETAMTINGPWAWSNIDT
MBP-1.4 PLIAADGGYAFKYENGKYDIKDVGVDNAGAKAGLTFLVALIKAKAMNADTDYSIAEAAFNKGETAMTINGPWAWSNIDT
MBP-2.5 PLIAADGGYAFKYENGKYDIKDVGVDNAGAKAGLTALVYLIAAKAMNADTDYSIAEAAFNKGETAMTINGPWAWSNIDT
MBP-3.6 PLIAADGGYAFKYENGKYDIKDVGVDNAGAKAGLTALVALIAAKAMNADTDYSIAEAAFNKGETAMTINGPWAWSNIDT

          240                 260                 280                 300
          |                   |                   |                   |
MBP-WT  SKVNYGVTVLPTFKGQPSKPFVGVLSAGINAASPNKELAKEFLENYLLTDEGLEAVNKDKPLGAVALKSYEEELAKDPR
MBP-1.1 SKVNYGVTVLPTFKGQPSKPFVGVLSAGINAASPNKELAKEFLENYLLTDEGLEAVNKDKPLGAVALKSYEEELAKDPR
MBP-1.2 SKVNYGVTVLPTFKGQPSKPFVGVLSAGINAASPNKELAKEFLENYLLTDEGLEAVNKDKPLGAVALKSYEEELAKDPR
MBP-1.3 SKVNYGVTVLPTFKGQPSKPFVGVLSAGINAASPNKELAKEFLENYLLTDEGLEAVNKDKPLGAVALKSYEEELAKDPR
MBP-1.4 SKVNYGVTVLPTFKGQPSKPFVGVLSAGINAASPNKELAKEFLENYLLTDEGLEAVNKDKPLGAVALKSYEEELAKDPR
MBP-2.5 SKVNYGVTVLPTFKGQPSKPFVGVLSAGINAASPNKELAKEFLENYLLTDEGLEAVNKDKPLGAVALKSYEEELAKDPR
MBP-3.6 SKVNYGVTVLPTFKGQPSKPFVGVLSAGINAASPNKELAKEFLENYLLTDEGLEAVNKDKPLGAVALKSYEEELAKDPR

           320                 340                 360
           |                   |                   |
MBP-WT  IAATMENAQKGEIMPNIPQMSAFWYAVRTAVINAASGRQTVDEALKDAQTRITK
MBP-1.1 IAATMENAQKGEIMPNIPQMSAFWYAVRTAVINAASG
MBP-1.2 IAATMENAQKGEIMPNIPQMSAFWYAVRTAVINAASG
MBP-1.3 IAATMENAQKGEIMPNIPQMSAFWYAVRTAVINAASG
MBP-1.4 IAATMENAQKGEIMPNIPQMSAFWYAVRTAVINAASG
MBP-2.5 IAATMENAQKGEIMPNIPQMSAFWYAVRTAVINAASGRQTVDEALKDAQTRITK
MBP-3.6 IAATMENAQKGEIMPNIPQMSAFWYAVRTAVINAASGRQTVDEALKDAQTRITK
```

**AR**

```
        12                 31                  51                  71
        |                  |                   |                   |
AR-WT   SDLGRKLLEAARAGQDDEVRILMANGADVNAADNTGTTPLHLAAYSGHLEIVEVLLKHGADVDASDVFGYTPLHLAAYW
AR-1.1  SDLGRKLLEAARAGQDDEVRILMANGADVNAADNTGTTPLHLAAYSGHLEIVEVLLKHGADVDASDVFGFTPLILAALW
AR-1.2  SDLGRKLLEAARAGQDDEVRILMANGADVNAADNTGTTPLHLAAYSGHLEIVEVLLKHGADVDASDVFGLTPLILAALW
AR-1.3  SDLGRKLLEAARAGQDDEVRILMANGADVNAADNTGTTPLHLAAYSGHLEIVEVLLKHGADVDASDVFGFTPLLLAALW
AR-1.4  SDLGRKLLEAARAGQDDEVRILMANGADVNAADNTGTTPLHLAAYSGHLEIVEVLLKHGADVDASDVFGMTPLFLAALW
AR-2.5  SDLGRKLLEAARAGQDDEVRILMANGADVNAADNTGTTPLHLAAYSGHLEIVEVLLKHGADVDASDVFGFTPLLLAALW
AR-2.6  SDLGRKLLEAARAGQDDEVRILMANGADVNAADNTGTTPLHLAAYSGHLEIVEVLLKHGADVDASDVFGFTPLGLAALW
AR-2.7  SDLGRKLLEAARAGQDDEVRILMANGADVNAADNTGTTPLHLAAYSGHLEIVEVLLKHGADVDASDVFGFTPLGLAALW
AR-3.8  SDLGRKLLEAARAGQDDEVRILMANGADVNAADNTGTTPLHLAAYSGHLEIVEVLLKHGADVDASDVFGFTPLGLAALW

        91                  111                 131                 151
        |                   |                   |                   |
AR-WT   GHLEIVEVLLKNGADVNAMDSDGMTPLHLAAKWGYLEIVEVLLKHGADVNAQDKFGKTAFDISIDNGNEDLAEILQKLN
AR-1.1  GHLEIVEVLLKNGADVNAMGSDGWTPLHAAAYFGYLEIVEVLLKHGADVNAQDKRGKTAFDISIDNGNEDLAEILQKLN
AR-1.2  GHLEIVEVLLKNGADVNAMDSDGWTPLHAAAYFGYLEIVEVLLKHGADVNAQDKRGKTAFDVSIDNGNEDLAEILQKLN
AR-1.3  GHLEIVEVLLKNGADVNAMGSDGWTPLHAAAWFGYLEIVEVLLKHGADVNAQDKRGKTAFDESIDNGNEDLAEILQKLN
AR-1.4  GHLEIVEVLLKNGADVNAMTSDGWTPLHAAAYFGYLEIVEVLLKHGADVNAQDKRGKTAFDVSIDNGNEDLAEILQKLN
AR-2.5  GHLEIVEVLLKHGEDVNAMGSDGWTPLHAAAWFGYLEIVEVLLKHGADVNAQDKRGKTAFDESIDNGNEDLAEILQKLN
AR-2.6  GHLEIVEVLLKHGEDVNAMGSDGWTPLHAAAWFGYLEIVEVLLKHGADVNAQDKRGKTAFDESIDNGNEDLAEILQKLN
AR-2.7  GHLEIVEVLLKHGEDVNAMGSDGWTPLHAAAKFGYLEIVEVLLKHGADVNAQDKRGKTPFDLAIDNGNEDIAEVLQKAA
AR-3.8  GHLEIVEVLLKHGEDVNAMGSDGWTPLHAAAKFGYLEIVEVLLKHGADVNAQDKKGLTPFDLAILNGNEDIAEVLQKAA
```

**Fig. S1. Sequence alignments of designed proteins (Table S1).** Residues that are different from the wild-type scaffold are highlighted, and motif residues are highlighted in yellow and boxed. Computationally predicted mutations for original designs are in magenta, stability-enhancing mutations from Kramer *et al.*(*33*) are in dark gray, mutations from error-prone PCR are in light grey, mutations from saturation mutagenesis are in orange, computationally predicted stabilizing mutations are in purple, and the computationally predicted reversion mutation is in blue.

### Figure S2

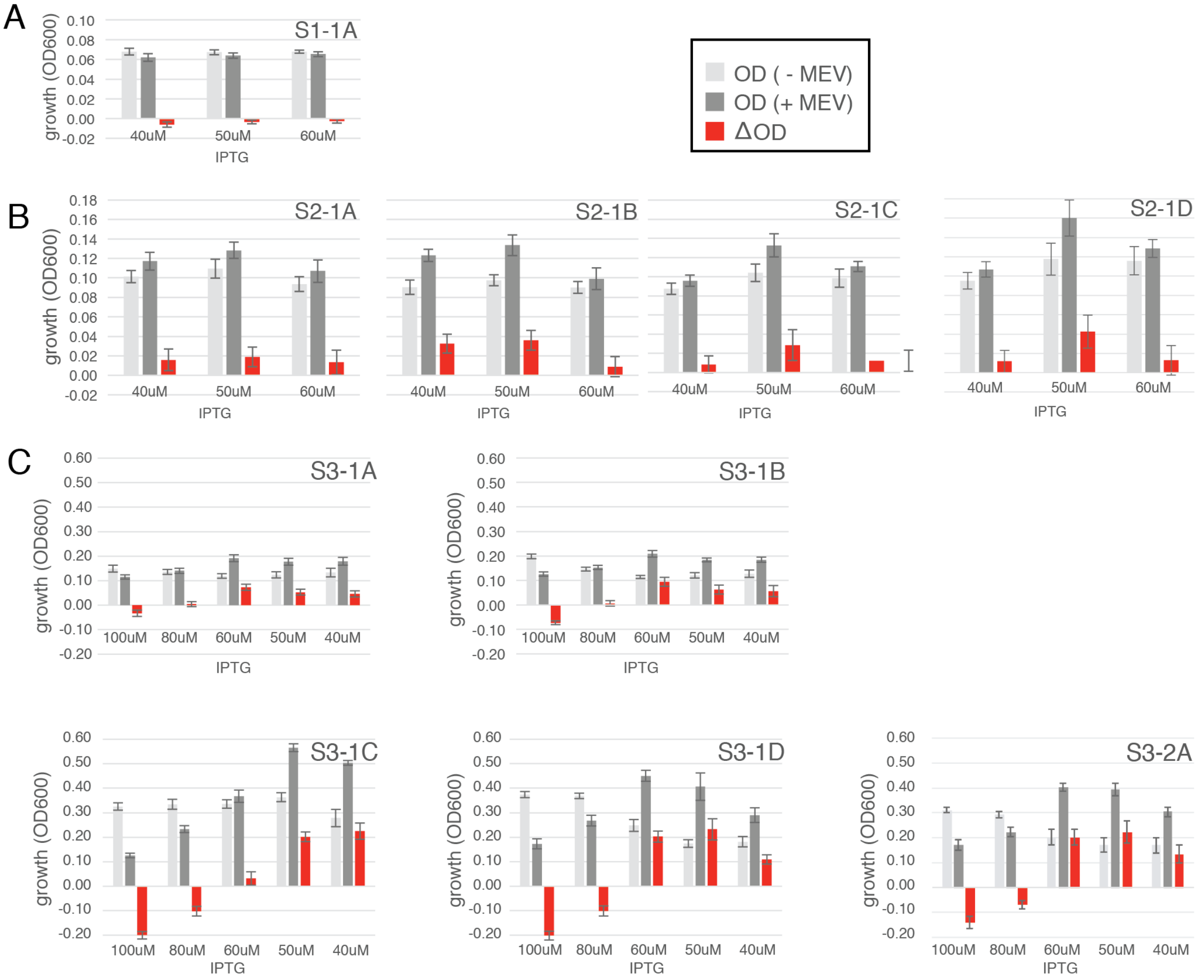

**Fig. S2. Growth +/- 5 mM mevalonate and change in growth for data shown in Figure 2C. (A)** Sensor design for scaffold 1 (FRB/FKBP). (**B**) Sensor designs for scaffold 2 (RapF/ComA). (**C**) Sensor designs for scaffold 3 (AR/MBP). Change in growth is dependent on IPTG concentration (panel C) as expected, where at high induction levels split mDHFR is complemented independent of mevalonate, and adding mevalonate leads to a growth disadvantage because of production of toxic metabolites (IPP, DMAP, and FPP). **Figure 2C** shows data at 50 µM IPTG. Experimental conditions: 0.4% L-arabinose, 1 µg/mL TMP, 35°C. Error bars are the standard deviation from at least 4 biological replicates and 8 replicates for each biological replicate.

Note that S3-2A, which contains two mutations introduced by error-prone PCR and was selected by library screening using the split mDHFR reporter, behaves similarly (except for a slight shift in the effect of IPTG concentration) to the original computational design S3-1C, which has an identical sequence without the two error-prone PCR mutations (**Fig. 2A**, **Fig. S5**).

**Figure S3**

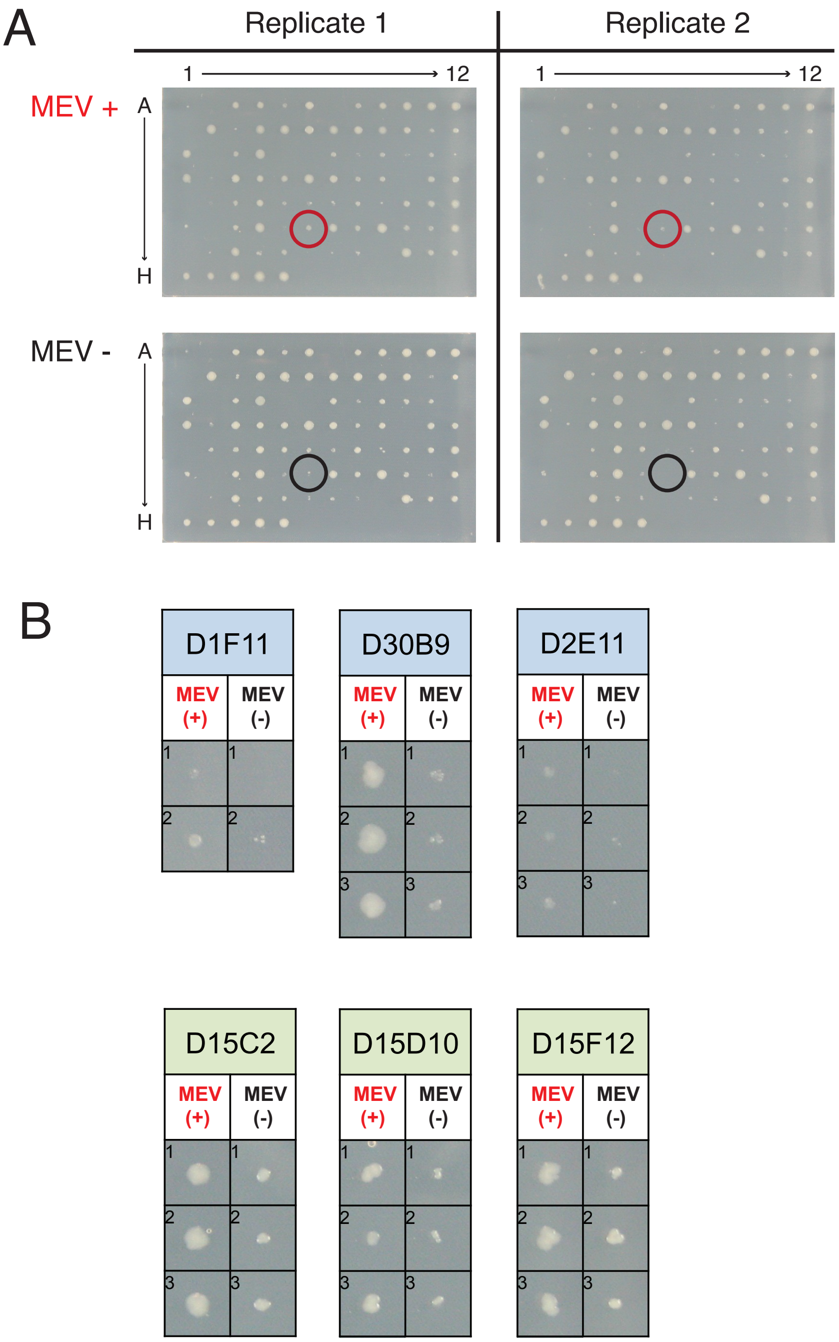

**Fig. S3. Example data from library screening using a colony-printing assay. (A)** Comparison of growth in the presence (top) and absence (bottom) of 5 mM mevalonate for two replicates. A colony growing better with mevalonate is circled in the replicates. (**B**) Examples of two to three replicates for library hits (concatenated from different plates). Experimental conditions: 0.4 % L-arabinose, 6 μg/mL TMP, 100 μM IPTG. Top row: hits from library 2; bottom row: hits from library 1.

**Figure S4**

**MBP**

```
           1                   20                  40                  60
           |                   |                   |                   |
MBP-WT     KIEEGKLVIWINGDKGYNGLAEVGKKFEKDTGIKVTVEHPDKLEEKFPQVAATGDGPDIIFWAHDRFGGYAQSGLLAEI
MBP-D2B1   KIEEGKLVIWINGDKGYNGLAEVGKKFEKDTGIKVTVEHPDKLEEKFPQVAATGDGPDIIFWAHDRFGGYAQSGLLAEI
MBP-D4A2   KIEEGKLVIWINGDKGYNGLAEVGKKFEKDTGIKVTVEHPDKLEEKFPQVAATGDGPDIIFWAHDRFGGYAQSGLLAEI
MBP-D11C1  KIEEGKLVIWINGDKGYNGLAEVGKKFEKDTGIKVTVEHPDKLEEKFPQVAATGDGPDIIFWAHDRFGGYAQSGLLAEI
MBP-D13E6  KIEEGKLVIWINGDKGYNGLAEVGKKFEKDTGIKVTVEHPDKLEEKFPQVAATGDGPDIIFWAHDRFGGYAQSGLLAEI
MBP-D15C2  KIEEGKLVIWINGDKGYNGLAEVGKKFEKDTGIKVTVEHPDKLEEKFPQVAATGDGPDIIFWAHDRFGGYAQSGLLAEI
MBP-D15D10 KIEEGKLVIWINGDKGYNGLAEVGKKFEKDTGIKVTVEHPDKLEEKFPQVAATGDGPDIIFWAHDRFGGYAQSGLLAEI
MBP-D15F12 KIEEGKLVIWINGDKGYNGLAEVGKKFEKDTGIKVTVEHPDKLEEKFPQVAATGDGPDIIFWAHDRFGGYAQSGLLAEI

           80                  100                 120                 140
           |                   |                   |                   |
MBP-WT     TPDKAFQDKLYPFTWDAVRYNGKLIAYPIAVEALSLIYNKDLLPNPPKTWEEIPALDKELKAKGKSALMFNLQEPYFTW
MBP-D2B1   TPDKAFQDKLYPFTWDAVRYNGKLIAYPIAVEALSLIYNKDLLPNPPKTWEEIHALDKELKAKGKSALMFNLQEPYFTW
MBP-D4A2   TPDKAFQDKLYPFTWDAVRYNGKLIAYPIAVEALSLIYNKDLLPNPPKTWEEIYALDKELKAKGKSALMFNLQEPYFTW
MBP-D11C1  TPDKAFQDKLYPFTWDAVRYNGKLIAYPIAVEALSLIYNKDLLPNPPKTWEEIHALDKELKAKGKSALMFNLQEPYFTW
MBP-D13E6  TPDKAFQDKLYPFTWDAVRYNGKLIAYPIAVEALSLIYNKDLLPNPPKTWEEIFALDKELKAKGKSALMFNLQEPYFTW
MBP-D15C2  TPDKAFQDKLYPFTWDAVRYNGKLIAYPIAVEALSLIYNKDLLPNPPKTWEEIHALDKELKAKGKSALMFNLQEPYFTW
MBP-D15D10 TPDKAFQDKLYPFTWDAVRYNGKLIAYPIAVEALSLIYNKDLLPNPPKTWEEIHALDKELKAKGKSALMFNLQEPYFTW
MBP-D15F12 TPDKAFQDKLYPFTWDAVRYNGKLIAYPIAVEALSLIYNKDLLPNPPKTWEEIHALDKELKAKGKSALMFNLQEPYFTW

            160                 180                 200                 220
            |                   |                   |                   |
MBP-WT     PLIAADGGYAFKYENGKYDIKDVGVDNAGAKAGLTFLVDLIKNKHMNADTDYSIAEAAFNKGETAMTINGPWAWSNIDT
MBP-D2B1   PLIAADGGYAFKYENGKYDIKDVGVDNAGAKAGLTFLVDLIKAKHMNADTDYSIAEAAFNKGETAMTINGPWAWSNIDT
MBP-D4A2   PLIAADGGYAFKYENGKYDIKDVGVDNAGAKAGLTFLVDLIKAKHMNADTDYSIAEAAFNKGETAMTINGPWAWSNIDT
MBP-D11C1  PLIAADGGYAFKYENGKYDIKDVGVDNAGAKAGLTFLVDLIKDKHMNADTDYSIAEAAFNKGETAMTINGPWAWSNIDT
MBP-D13E6  PLIAADGGYAFKYENGKYDIKDVGVDNAGAKAGLTFLVALIAAKAMNADTDYSIAEAAFNKGETAMTINGPWAWSNIDT
MBP-D15C2  PLIAADGGYAFKYENGKYDIKDVGVDNAGAKAGLTFLVDLIKNKHMNADTDYSIAEAAFNKGETAMTINGPWAWSNIDT
MBP-D15D10 PLIAADGGYAFKYENGKYDIKDVGVDNAGAKAGLTFLVGLIEDKHMNADTDYSIAEAAFNKGETAMTINGPWAWSNIDT
MBP-D15F12 PLIAADGGYAFKYENGKYDIKDVGVDNAGAKAGLAFLVDLIKDKHMNADTDYSIAEAAFNKGETAMTINGPWAWSNIDT

              240                 260                 280                 300
              |                   |                   |                   |
MBP-WT     SKVNYGVTVLPTFKGQPSKPFVGVLSAGINAASPNKELAKEFLENYLLTDEGLEAVNKDKPLGAVALKSYEEELAKDPR
MBP-D2B1   SKVNYGVTVLPTFKGQPSKPFVGVLSAGINAASPNKELAKEFLENYLLTDEGLEAVNKDKPLGAVALKSYEEELAKDPR
MBP-D4A2   SKVNYGVTVLPTFKGQPSKPFVGVLSAGINAASPNKELAKEFLENYLLTDEGLEAVNKDKPLGAVALKSYEEELAKDPR
MBP-D11C1  SKVNYGVTVLPTFKGQPSKPFVGVLSAGINAASPNKELAKEFLENYLLTDEGLEAVNKDKPLGAVALKSYEEELAKDPR
MBP-D13E6  SKVNYGVTVLPTFKGQPSKPFVGVLSAGINAASPNKELAKEFLENYLLTDEGLEAVNKDKPLGAVALKSYEEELAKDPR
MBP-D15C2  SKVNYGVTVLPTFKGQPSKPFVGVLSAGINAASPNKELAKEFLENYLLTDEGLEAVNKDKPLGAVALKSYEEELAKDPR
MBP-D15D10 SKVNYGVTVLPTFKGQPSKPFVGVLSAGINAASPNKELAKEFLENYLLTDEGLEAVNKDKPLGAVALKSYEEELAKDPR
MBP-D15F12 SKVNYGVTVLPTFKGQPSKPFVGVLSAGINAASPNKELAKEFLENYLLTDEGLEAVNKDKPLGAVALKSYEEELAKDPR

               320                 340
               |                   |
MBP-WT     IAATMENAQKGEIMPNIPQMSAFWYAVRTAVINAA
MBP-D2B1   IAATMENAQKGEIMPNIPQMSAFWYAVRTAVINAA
MBP-D4A2   IAATMENAQKGEIMPNIPQMSAFWYAVRTAVINAA
MBP-D11C1  IAATMENAQKGEIMPNIPQMSAFWYAVRTAVINAA
MBP-D13E6  IAATMENAQKGEIMPNIPQMSAFWYAVRTAVINAA
MBP-D15C2  IAATMENAQKGEIMPNIPQMSAFWYAVRTAVINAA
MBP-D15D10 IAATMENAQKGEIMPNIPQMSAFWYAVRTAVINAA
MBP-D15F12 IAATMENAQKGEIMPNIPQMSAFWYAVRTAVINAA
```

**AR**

```
             12                 31                  51                  71
             |                  |                   |                   |
AR-WT        SDLGRKLLEAARAGQDDEVRILMANGADVNAADNTGTTPLHLAAYSGHLEIVEVLLKHGADVDASDVFGYTPLHLAAYW
AR-D2B1      SDLGRKLLEAARAGQDDEVRILMANGADVNAADNTGTTPLHLAAYSGHLEIVEVLLKHGADVDASDVAGITPLILAAMW
AR-D4A2      SDLGRKLLEAARAGQDDEVRILMANGADVNAADNTGTTPLHLAAYSGHPEIVEVLLKHGADVDASDVAGLTPLFLAAMW
AR-D11C1     SDLGRKLLEAARAGQDDEVRILMANGADVNAADNTGTTPLHLAAYSGHLEIVEVLLKHGADVDASDVAGITPLFLAAMW
AR-D13E6     SDLGRKLLEAARAGQDDEVRILMANGADVNAADNTGTTPLHLAAYSGHLEIVEVLLKHGADVDASDVAGLTPLILAAFW
AR-D15C2     SDLGRKLLEAARAGQDDEVRILMANGADVNAADNTGTTPLHLAAYSGHLEIVEVLLKHGADVDASDVAGMTPLILAAFW
AR-D15D10    SDLGRKLLEAARAGQDDEVRILMANGADVNAADNTGTTPLHLAAYSGHLEIVEVLLKHGADVDASDVAGLTPLILAAMW
AR-D15F12    SDLGRKLLEAARAGQDDEVRILMANGADVNAADNTGTTPLHLAAYSGHLEIVEVLLKHGADVDASDVAGITPLILAAMW

              91                  111                 131                 151
              |                   |                   |                   |
AR-WT        GHLEIVEVLLKNGADVNAMDSDGMTPLHLAAKWGYLEIVEVLLKHGADVNAQDKFGKTAFDISIDNGNEDLAEILQKLN
AR-D2B1      GHLEIVEVLLKNGADVNAMDSDGWTPLHAAAYLGYLEIVEVLLKHGADVNAQDKRGKTAFDTSIDNGNEDLAEILQKLN
AR-D4A2      GHLEIVEVLLKNGADVNAMDSDGWTPLHAAAVLGYLEIVEVLLKHGADVNAQDKRGKTAFDVSIDNGNEDLAEILQKLN
AR-D11C1     GHLEIVEVLLKNGADVNAMDSDGWTPLHAAADAGYLEIVEVLLKHGADVNAQDKRGKTAFDTSIDNGNEDLAEILQKLN
AR-D13E6     GHLEIVEVLLKNGADVNAMDSDGWTPLHAAASYGYLEIVEVLLKHGADVNAQDKRGKTAFDISIDNGNEDLAEILQKLN
AR-D15C2     GHLEIVEVLLKNGADVNAMDSDGWTPLHAAAVVGYLEIVEVLLKHGADVNAQDKRGKTAFDTSIDNGNEDLAEILQKLN
AR-D15D10    GHLEIVEVLLKNGADVNAMDSDGWTPLHAAADEGYLEIVEVLLKHGADVNAQDKRGKTAFDTSIDNGNEDLAEILQKLN
AR-D15F12    GHLEIVEVLLKNGADVNAMDSDGWTPLHAAAYYGYLEIVEVLLKHGADVNAQDKRGKTAFDISIDNGNEDLAEILQKLN
```

**Fig. S4. Sequence alignments of hits from the computationally designed S3 library.** These variants were identified using a plate-based split mDHFR assay (**Fig. S3**) and validated by individual solution growth assays (Methods). Residues different from the WT scaffold are highlighted in magenta, and motif residues are highlighted in yellow and boxed.

### Figure S5

### MBP

```
           1                    20                    40                    60
MBP-WT     KIEEGKLVIWINGDKGYNGLAEVGKKFEKDTGIKVTVEHPDKLEEKFPQVAATGDGPDIIFWAHDRFGGYAQSGLLAEI
MBP-D1F11  KIEEGKLVIWINGDKGYNGLAEVGKKFEKDTGIKVTVEHPDKLEEKFPQVAATGDGPDIIFWAHDRFGGYAQSGLLAEI
MBP-D2E11  KIEEGKLVIWINGDKGYNGLAEVGKKFEKDTGIKVTVEHPDKLEEKFPQVAATGDGPDIIFWAHDRFGGYAQSGLLAEI
MBP-D23E3  KIEEGKLVIWINGDKGYNGLAEVGKKFEKDTGIKVTVEHPDKLEEKFPQVAATGDGPDIIFWAHDRFGGYAQSGLLAEI
MBP-D26B5  KIEEGKLVIWINGDKGYNGLAEVGKKFEKDTGIKVTVEHPDKLEEKFPQVAATGDGPDIIFWAHDRFGGYAQSGLLAEI
MBP-D28D9  KIEEGKLVIWINGDKGYNGLAEVGKKFEKDTGIKVTVEHPDKLEEKFPQVAATGDGPDIIFWAHDRFGGYAQSGLLAEI
MBP-D28H1  KIEEGKLVIWINGDKGYNGLAEVGKKFEKDTGIKVTVEHPDKLEEKFPQVAATGDGPDIIFWAHDRFGGYAQSGLLAEI
MBP-D29F4  KIEEGKLVIWINGDKGYNGLAEVGKKFEKDTGIKVTVEHPDKLEEKFPQVAATGDGPDIIFWAHDRFGGYAQSGLLAEI
MBP-D30B9  KIEEGKLVIWINGDKGYNGLAEVGKKFEKDTGIKVTVEHPDKLEEKFPQVAATGDGPDIIFWAHDRFGGYAQSGLLAEI
MBP-D30F9  KIEEGKLVIWINGDKGYNGLAEVGKKFEKDTGIKVTVEHPDKLEEKFPQVAATGDGPDIIFWAHDRFGGYAQSGLLAEI
MBP-D31D9  KIEEGKLVIWINGDKGYNGLAEVGKKFEKDTGIKVTVEHPDKLEEKFPQVAATGDGPDIIFWAHDRFGGYAQSGLLAEI
MBP-D31E8  KIEEGKLVIWINGDKGYNGLAEVGKKFEKDTGIKVTVEHPDKLEEKFPQVAATGDGPDIIFWAHDRFGGYAQSGLLAEI

           80                   100                   120                   140
MBP-WT     TPDKAFQDKLYPFTWDAVRYNGKLIAYPIAVEALSLIYNKDLLPNPPKTWEEIPALDKELKAKGKSALMFNLQEPYFTW
MBP-D1F11  TPDKAFQDKLYPFTWDAVRYNGKLIAYPIAVEALSLIYNKDLLPNPPKTWEEIFALDKELKAKGKSALMFNLQEPYFTW
MBP-D2E11  TPDKAFQDKLYPFTWDAVRYNGKLIAYPIAVEALSLIYNKDLLPNPPKTWEEIFALDKELKAKGKSALMFNLQEPYFTW
MBP-D23E3  TPDKAFQDKLYPFTWDAVRYNGKLIAYPIAVEALSLIYNKDLLPNPPKTWEEIFALDKELKAKGKSALMFNLQEPYFTW
MBP-D26B5  TPDKAFQDKLYPFTWDAVRYNGKLIAYPIAVEALSLIYNKDLLPNPPKTWEEIFALDKELKAKGKSALMFNLQEPYFTW
MBP-D28D9  TPDKAFQDKLYPFTWDAVRYNGKLIAYPIAVEALSLIYNKDLLPNPPKTWEEIFALDKELKAKGKSALMFNLQEPYFAW
MBP-D28H1  TPDKAFQDKLYPFTWDAVRYNGKLIAYPIAVEALSLIYNKDLLPNPPKTWEEIFALDKELKAKGKSALMFNLQEPYFTW
MBP-D29F4  TPDKAFQDKLYPFTWDAVRYNGKLIAYPIAVEALSLIYNKDLLPNPPKTWEEIFALDKELKAKGKSALMFNLQEPYFTW
MBP-D30B9  TPDKAFQDKLYPFTWDAVRYNGKLIAYPIAVEALSLIYNKDLLPNPPKTWEEIFALDKELKAKGKSALMFNLQEPYFTW
MBP-D30F9  TPDKAFQDKLYPFTWDAVRYNGKLIAYPIAVEALSLIYNKDLLPNPPKTWEEIFALDKELKAKGKSALMFNLQEPYFTW
MBP-D31D9  TPDKAFQDKLYPFTWDAVRYNGKLIAYPIAVEALSLIYNKDLLPNPPKTWEEIFALDKELKAKGKSALMFNLQEPYFTW
MBP-D31E8  TPDKAFQDKLYPFTWDAVRYNGKLIAYPIAVEALSLIYNKDLLPNPPKTWEEIFALDKELKAKGKSALMFNLQEPYFTW

           160                  180                   200                   220
MBP-WT     PLIAADGGYAFKYENGKYDIKDVGVDNAGAKAGLTFLVDLIKNKHMNADTDYSIAEAAFNKGETAMTINGPWAWSNIDT
MBP-D1F11  PLIAADGGYAFKYENGKYDIKDVGVDNAGAKAGLTRLVYLIAAKAMNADTDYSIAEAAFNKGETAMTINGPWAWSNIDT
MBP-D2E11  PLIAADGGYAFKYENGKYDIKDVGVDNAGAKAGLTRLVYLIAAKAMNADTDYSIAEAAFNKGETAMTINGPWAWSNIDT
MBP-D23E3  PLIAADGGYAFKYENGKYDIKDVGVDNAGAKAGLTRLVYLIAAKAMNADTDYSIAEAAFNKGETAMTINGPWAWSNIDT
MBP-D26B5  PLIAADGGYAFKYENGKYDIKDVGVDNAGAKAGLTRLVYLIAAKAMNADTDYSIAEAAFNKGETAMTINGPWAWSNIDT
MBP-D28D9  PLIAADGGYAFKYENGKYDIKDVGVDNAGAKAGLTRLVYLIAAKAMNADTDYSIAEAAFNKGETAMTINGPWAWSNIDT
MBP-D28H1  PLIAADGGYAFKYENGKYDIKDVGVDNAGAKAGLTRLVYLIAAKAMNADTDYSIAEAAFNKGETAMTINGPWAWSNIDT
MBP-D29F4  PLIAADGGYAFKYENGKYDIKDVGVDNAGAKAGLTRLVYLIAAKAMNADTDYSIAEAAFNKGETAMTINGPWAWSNIDT
MBP-D30B9  PLIAADGGYAFKYENGKYDIKDVGVDNAGAKAGLTFLVALIAAKAMNADTDYSIAEAAFNKGETAMTINGPWAWSNIDT
MBP-D30F9  PLIAADGGYAFKYENGKYDIKDVGVDNAGAKAGLTRLVYLIAAKAMNADTDYSIAEAAFNKGETAMTINGPWAWSNIDT
MBP-D31D9  PLIAADGGYAFKYENGKYDIKDVGVDNAGAKAGLTRLVYLIAAKAMNADTDYSIAEAAFNKGETAMTINGPWAWSNIDT
MBP-D31E8  PLIAADGGYAFKYENGKYDIKDVGVDNAGAKAGLTRLVYLIAAKAMNADTDYSIAEAAFNKGETAMTINGPWAWSNIDT

            240                  260                   280                   300
MBP-WT     SKVNYGVTVLPTFKGQPSKPFVGVLSAGINAASPNKELAKEFLENYLLTDEGLEAVNKDKPLGAVALKSYEEELAKDPR
MBP-D1F11  SKVNYGVTVLPTFKGQPSKPFVGVLSAGINAASPNKELAKEFLENYLLTDEGLEAVNKDKPLGAVALKSYEEELAKDPR
MBP-D2E11  SKVNYGVTVLPTFKGQPSKPFVGVLSAGINAASPNKELAKEFLENYLLTDEGLEAVNKDKPLGAVALKSYEEELAKDPR
MBP-D23E3  SKVNYGVTVLPTFKGQPSKPFVGVLSAGINAASPNKELAKEFLENYLLTDEGLEAVNKDKPLGAVALKSYEEELAKDPR
MBP-D26B5  SKVNYGVTVLPTFKGQPSKPFVGVLSAGINAASPNKELAKEFLENYLLTDEGLEAVNKDKPLGAVALKSYEEELAKDPR
MBP-D28D9  SKVNYGVTVLPTFKGQPSKPFVGVLSAGINAASPNKELAKEFLENYLLTDEGLEAVNKDKPLGAVALKSYEEELAKDPR
MBP-D28H1  SKVNYGVTVLPTFKGQPSKPFVGVLSAGINAASPNKELAKEFLENYLLTDEGLEAVNKDKPLGAVALKSYEEELAKDPR
MBP-D29F4  SKVNYGVTVLPTFKGQPSKPFVGVLSAGINAASPNKELAKEFLENYLLTDEGLEAVNKDKPLGAVALKSYEEELAKDPR
MBP-D30B9  SKVNYGVTVLPTFKGQPSKPFVGVLSAGINAASPNKELAKEFLENYLLTDEGLEAVNKDKPLGAVALKSYEEELAKDPR
MBP-D30F9  SKVNYGVTVLPTFKGQPSKPFVGVLSAGINAASPNKELAKEFLENYLLTDEGLEAVNKDKPLGAVALKSYEEELAKDPR
MBP-D31D9  SKVNYGVTVLPTFKGQPSKPFVGVLSAGINAASPNKELAKEFLENYLLTDEGLEAVNKDKPLGAVALKSYEEELAKDPR
MBP-D31E8  SKVNYGVTVLPTFKGQPSKPFVGVLSAGINAASPNKELAKEFLENYLLTDEGLEAVNKDKPLGAVALKSYEEELAKDPR

             320                  340
MBP-WT     IAATMENAQKGEIMPNIPQMSAFWYAVRTAVINAA
MBP-D1F11  IAATMENAQKGEIMPNIPQMSAFWYAVRTAVINAA
MBP-D2E11  IAATMENAQKGEIMPNIPQMSAFWYAVRTAVINAA
MBP-D23E3  IAATMENAQKGEIMPNIPQMSAFWYAVRTAVINAA
MBP-D26B5  IAATMENAQKGEIMPNIPQMSAFWYAVRTAVINAA
MBP-D28D9  IAATMENAQKGEIMPNIPQMSAFWYAVRTAVINAA
MBP-D28H1  IAATMENAQKGEIMPNIPQMSAFWYAVRTAVINAA
MBP-D29F4  IAATMENAQKGEIMPNIPQMSAFWYAVRTAVINAA
MBP-D30B9  IAATMENAQKGEIMPNIPQMSAFWYAVRTAVINAA
MBP-D30F9  IAATMENAQKGEIMPNIPQMSAFWYAVRTAVINAA
MBP-D31D9  IAATMENAQKGEIMPNIPQMSAFWYAVRTAVINAA
MBP-D31E8  IAATMENAQKGEIMPNIPQMSAFWYAVRTAVINAA
```

**AR**

```
          12                 31                  51                  71
          |                  |                   |                   |
AR-WT     SDLGRKLLEAARAGQDDEVRILMANGADVNAADNTGTTPLHLAAYSGHLEIVEVLLKHGADVDASDVFGYTPLHLAAYW
AR-D1F11  SDLGRKLLEAARAGQDDEVRILMANGADVNAADNTGTTPLHLAAYSGHLEIVEVLLKHGADVDASDVFGFTPLLLAALW
AR-D2E11  SDLGRKLLEAARAGQDDEVRILMANGADVNAADNTGTTPLHLAAYSGHLEIVEVLLKHGADVDASDVFGFTPLLLAALW
AR-D23E3  SDLGRKLLEAARAGQDDEVRILMANGADVNAADNTGTTPLHLAAYSGHLEIVEVLLKHGADVDASDVFGFTPLLLAALW
AR-D26B5  SDLGRKLLEAARAGQDDEVRILMANGADVNAADNTGTTPLHLAAYSGHLEIVEVLLKHGADVDASDVFGFTPLLLAALW
AR-D28D9  SDLGRKLLEAARAGQDDEVRILMANGADVNAADNTGTTPLHLAAYSGHLEIVEVLLKHGADVDASDVFGFTPLLLAALW
AR-D28H1  SDLGRKLLEAARAGQDDEVRILMANGADVNAADNTGTTPLHLAAYSGHLEIVEVLLKHGADVDASDVFGFTPLLLAALW
AR-D29F4  SDLGRKLLEAARAGQDDEVRILMANGADVNAADNTGTTPLHLAAYSGRLEIVEVLLKHGADVDASDVFGFTPLLLAALW
AR-D30B9  SDLGRKLLEAARAGQDDEVRILMANGADVNAADNTGTTPLHLAAYSGHLEIVEVLLKHGADVDASDVFGFTPLLLAALW
AR-D30F9  SDLGRKLLEAARAGQDDEVRILMANGADVNAADNTGTTPLHLAAYSGHLEIVEVLLKHGADVDASDVFGFTPLLLAALW
AR-D31D9  SDLGRKLLEAARAGQDDEVRILMANGADVNAADNTGTTPLHLAAYSGHLEIVEVLLKHGADVDASDVFGFTPLLLAALW
AR-D31E8  SDLGRKLLEAARAGQDDEVRILMANGADVNAADNTGTTPLHLAAYSGHLEIVEVLLKHGADVDASDVFGFTPLLLAALW

          91                  111                 131                 151
          |                   |                   |                   |
AR-WT     GHLEIVEVLLKNGADVNAMDSDGMTPLHLAAKWGYLEIVEVLLKHGADVNAQDKFGKTAFDISIDNGNEDLAEILQKLN
AR-D1F11  GHLEIVEVLLKHGEDVNAMGSDGWTPLHAAAWFGYLEIVEVLLKHGADVNAQDKRGKTAFDESIDNGNEDLAEILQKLN
AR-D2E11  GHLEIVEVLLKNGEDVNAMGSDGWTPLHAAAWFGYLEIVEVLLKHGADVNAQDKRGKTAFDESIDNGNEDLAEILQKLN
AR-D23E3  GHLEIVEVLLKNGADVNAMGSDGWTPPHTAAWFGYLEIVEVLLKHGADVNAQDKRGKTAFDESIDNGNEDLAEILQKLN
AR-D26B5  GHLEIVEVLLKNGADVNAMGSDGWTPLHAAAWFGYLGNVEVLLKHGADVNAQDKRGKTAFDESIDNGNEDLAEILQKLN
AR-D28D9  GHLEIVEVLLKNGADVNAMGSDGWTPLHAAAWFGYLEIEEVLLKLGADVNAQDKRGKTAFDESIDNGNEDLAEILQKLN
AR-D28H1  GHLEIVEVLLKNGADVNAMGSDGWTPLHAAAWFGYLEFVEVLLQHGADVNAQDKRGKTAFDESIDNGNEDLAEILQKLN
AR-D29F4  GHLEIVEVLLKNGTGVNAMGSDGWTPLHAAAWFGYLEIVEVLLKHGADVNAQDKRGKTAFDESIDNGNEDLAEILQKLN
AR-D30B9  GHLEIVEVLLKNGADVNAMGSDGWTPLHAAAWFGYLEFVEVLLKHGADVNAQDKRGKTAFDESIDNGNEDLAEILQKLN
AR-D30F9  GHLEIVEVLLKNGAVVNAMGSDGWTPLLAAAWFGYLEIVEVLLKHGADVNAQDKRGKTAFDESIDNGNEDLAEILQKLN
AR-D31D9  GHLEIVEVLLKNGADVNAMGSDGWTPLPAAAWFGYLEIVEDLLKHGADVNAQDKRGKTAFDESIDNGNEDLAEILQKLN
AR-D31E8  GHLEIVEVLLKNGADVNAMGSDGWTPPHAAAWFGYLEIVEVLLKHGADVNAQDKRGKTAFDESIDNGNEDLAEILQKLN
```

**Fig. S5. Sequence alignments of hits from the error-prone PCR S3 library.** These variants were identified using a plate-based split mDHFR assay (**Fig. S3**) and validated by individual solution growth assays (Methods). Residues different from the WT scaffold are highlighted in magenta, and motif residues are highlighted in yellow and boxed.

### Figure S6

A

| | 85 | 118 | 119 | 122 | 152 | 194 |
|---|---|---|---|---|---|---|
| | AR | | | | | MBP |
| **S3-2A** | L | H | A | W | E | R |
| **Round 1, starting from S3-2A** | | | | | | |
| S1B5 | L | H | A | V | E | R |
| S1C4 | L | L | A | W | E | R |
| S1D10 | L | P | A | W | E | R |
| S1E5 | L | N | A | W | E | R |
| S2F9 | L | H | A | L | E | R |
| S3H7 | L | H | A | W | E | F |
| S3H11 | L | H | A | W | E | G |
| **S3-2B** | L | H | A | W | E | A |
| **Round 2, starting from S2-2B** | | | | | | |
| S5B3 | L | T | A | W | E | A |
| S5B4 | L | I | A | W | E | A |
| S7G5 | L | H | P | W | E | A |
| S7H7 | L | H | R | W | E | A |
| **S3-2C** | G | H | A | W | E | A |

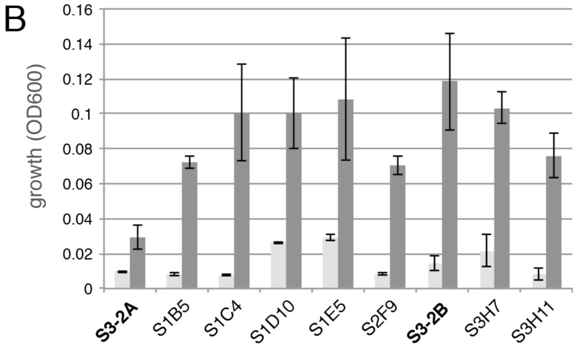

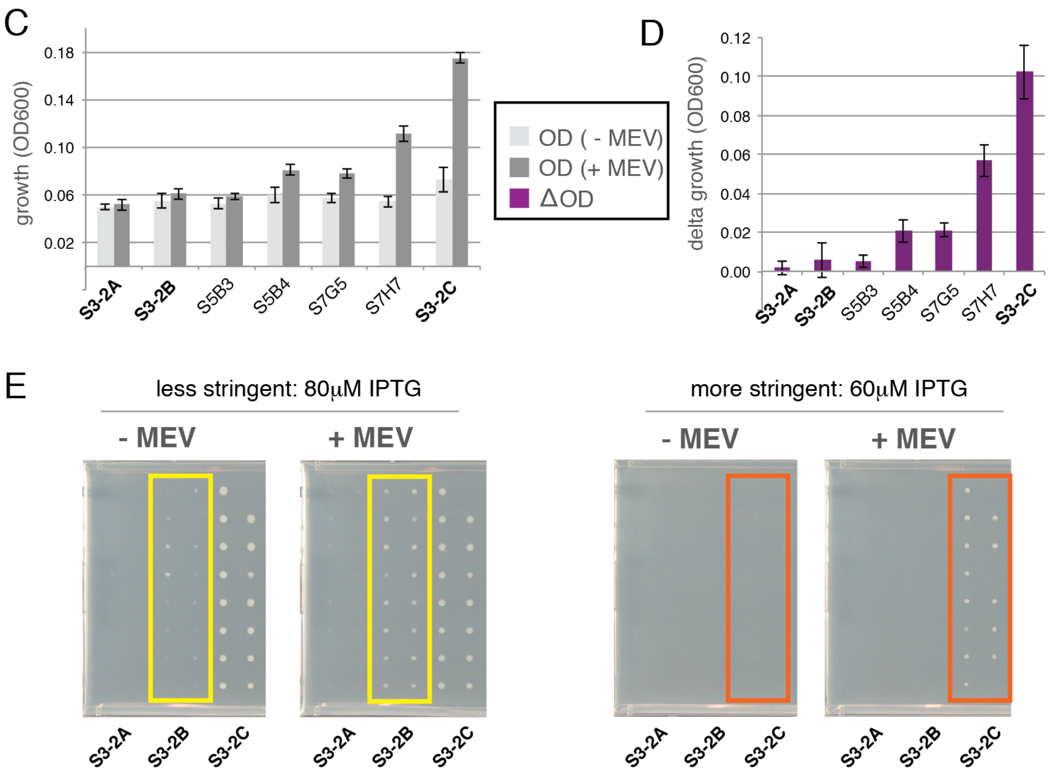

**Fig. S6. Data from the single-site saturation mutagenesis screen. (A)** Sequences of hits at screened positions. (**B**) Validation of hits from round 1 (starting from S3-2A) with the split mDHFR assay in liquid culture. (**C**) Validation of hits from round 2 (starting from S3-2B) with split mDHFR assay in liquid culture. (**D**) Sensor signal (change in growth as measured by $OD_{600}$) from round 2. (**E**) Split mDHFR plate assay showing that sensors S3-2B and S3-2C function under more stringent conditions (lower IPTG inducer concentration). Experimental conditions: Round 1 (panel B): cultures grown with 80 μM IPTG at 35 °C, *n*=3. Round 2: (panels C, D): cultures grown with 60 μM IPTG at 35 °C, *n*=4. Error bars reflect standard deviation.

### Figure S7

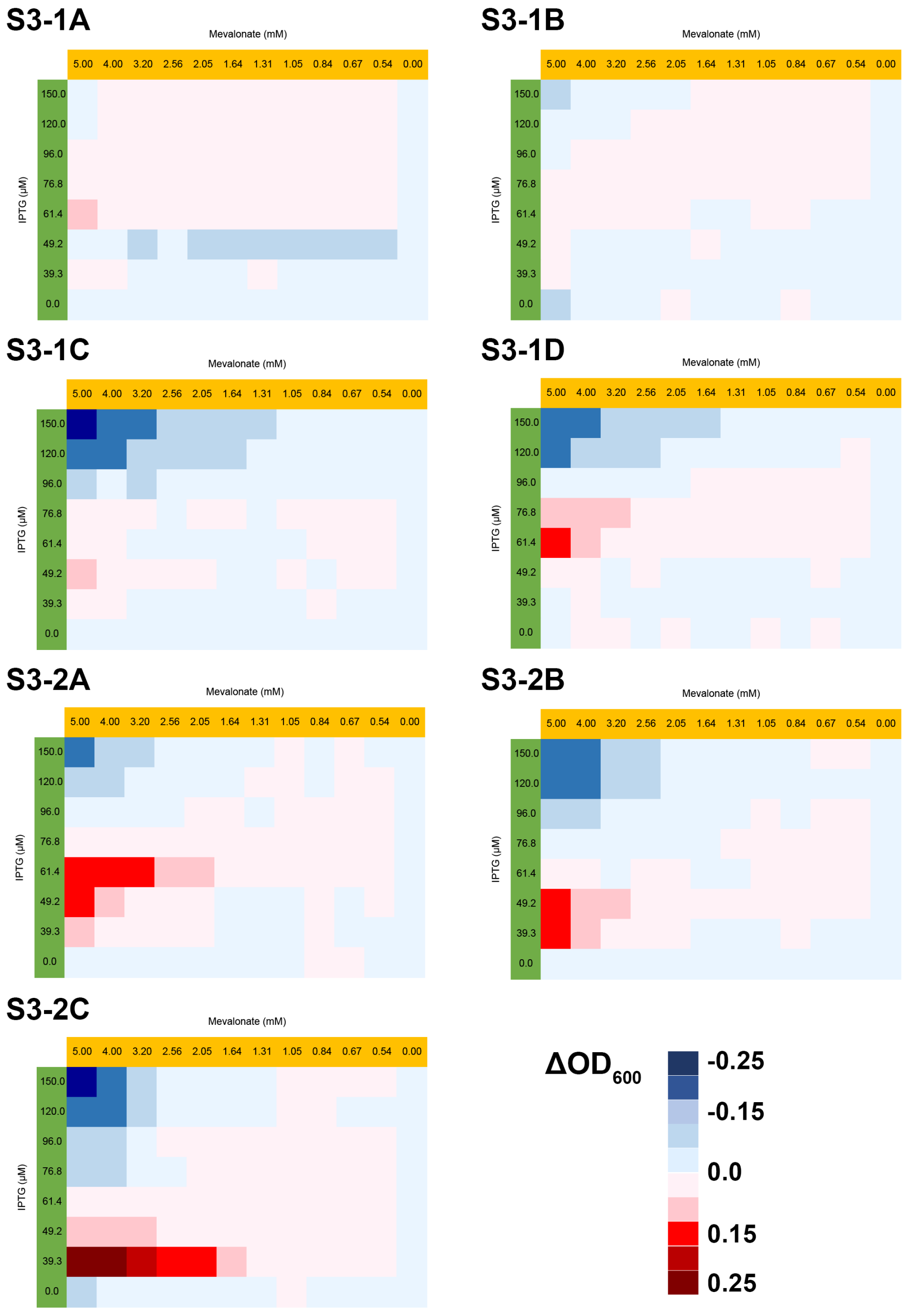

**Fig. S7. Dependency of the sensor signal on IPTG and mevalonate concentrations.** Data are from a split mDHFR assay in liquid culture. Experimental conditions: varied concentrations of IPTG and mevalonate, 0.4% L-arabinose, 1 μg/ml TMP, cultured at 35 °C. Data are averaged over 6 plates per design.

**Figure S8**

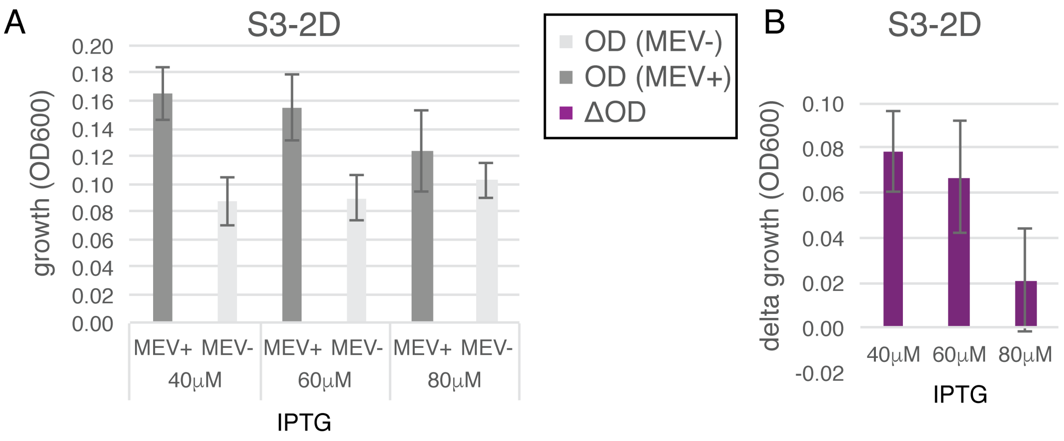

**Fig. S8. (A) Growth +/- mevalonate and (B) change in growth for design S3-2D.** Experimental conditions: 35 °C, 1 μg/ml TMP, 5 mM mevalonate. Error bars are standard deviation from at least 4 biological replicates and 8 replicates for each biological replicate.

**Figure S9**

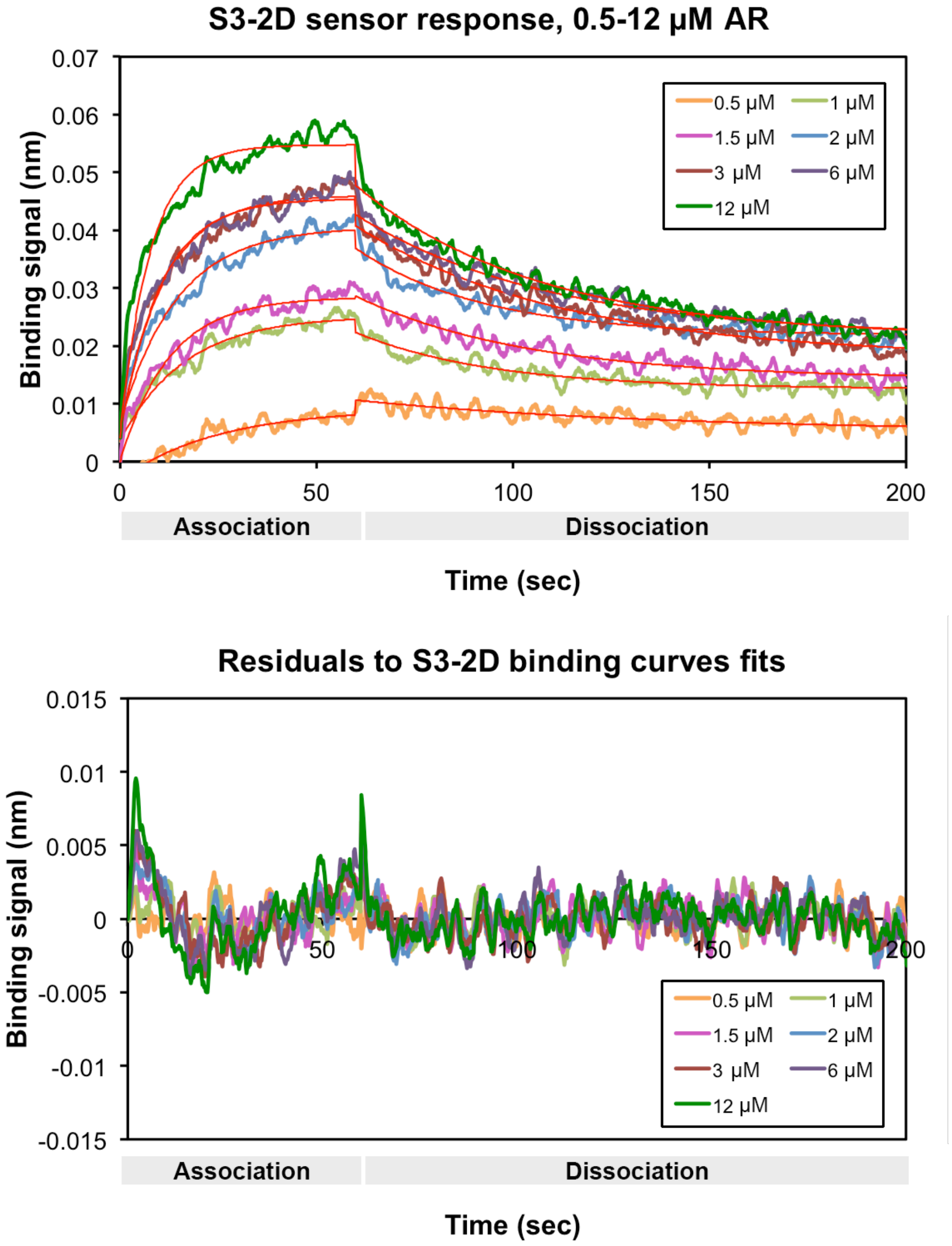

**Fig. S9. BLI binding assay representative data and fits.** The apparent $K_D$ of design S3-2D was measured using immobilized avi-MBP-2.5, titrated AR-2.7 at indicated concentrations (top), in the presence of 200 µM FPP. Red lines in the top plot indicate fit to the data and residuals are shown in the bottom plot.

**Figure S10**

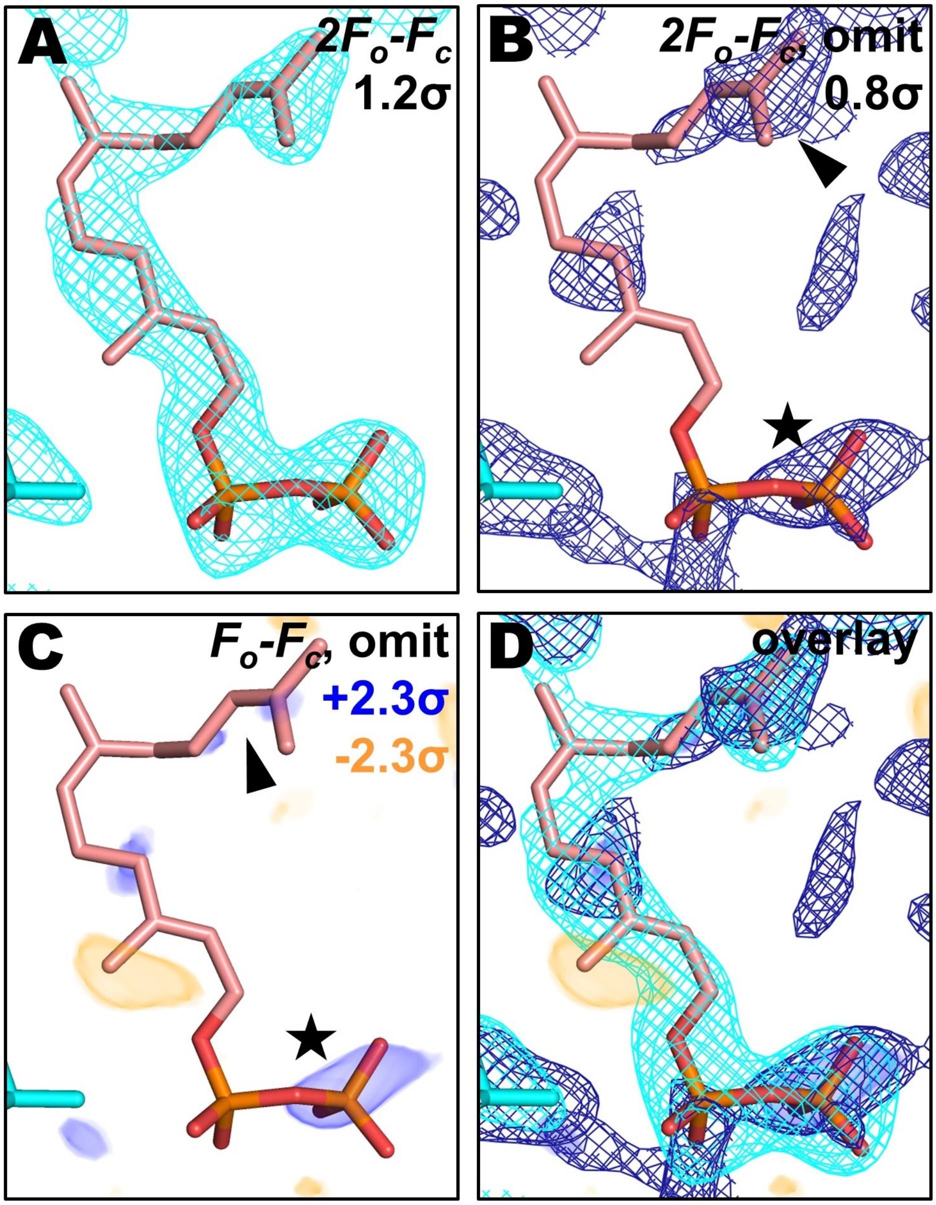

**Fig. S10. Electron density maps supporting placement of the FPP ligand.** The $2F_o$-$F_c$ electron density map created using the final model phases (**A**, cyan mesh) shows good agreement with the modeled ligand (RSCC=0.89). The $2F_o$-$F_c$ (**B**, dark blue mesh) and $F_o$-$F_c$ (**C**, blue/orange volumes) omit maps contain notable peaks corresponding to the pyrophosphate group (marked with a star) and the aliphatic tail (marked with an arrow), which resembled peaks in the $2F_o$-$F_c$ and $F_o$-$F_c$ maps that were used to initially identify the ligand. An overlay of the different electron density maps (**D**) shows the co-localization of these features in real space.

**Figure S11**

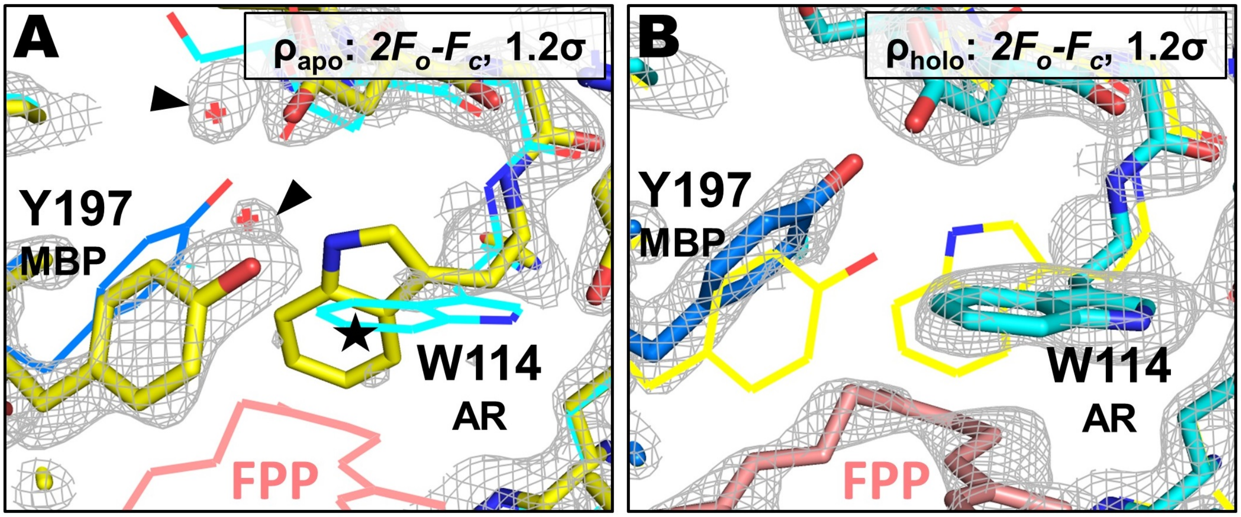

**Fig. S11. Electron density supporting ligand-induced rearrangement of the FPP binding site.** In the unoccupied FPP binding site (yellow models, density shown in panel (**A**), Y197 in the MBP monomer is rotated down into the binding pocket, and W114 of the AR monomer appears flexible, likely occupying multiple rotameric states (denoted by a star). When the binding site is occupied (MBP models in blue, AR models in cyan, density shown in panel (**B**), the Y197 side chain rotates away from the FPP molecule, displacing a small network of neighboring water molecules (indicated by the black arrows in panel A), and W114 becomes well-ordered with its indole group packed against the bound FPP molecule (pink).

**Figure S12**

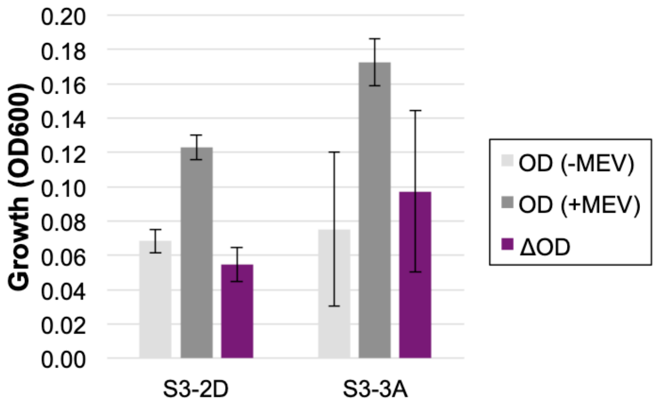

**Fig. S12. Comparison of S3-2D and S3-3A (Y197A mutation) in the split mDHFR assay.** Design S3-3A is an active sensor in *E. coli*. Experimental conditions: 35 °C, 60 µM IPTG, 6 µg/ml TMP, 0.4% L-arabinose. Error bars are standard deviations, *n*=8 for each design and +/- mevalonate condition.

**Figure S13**

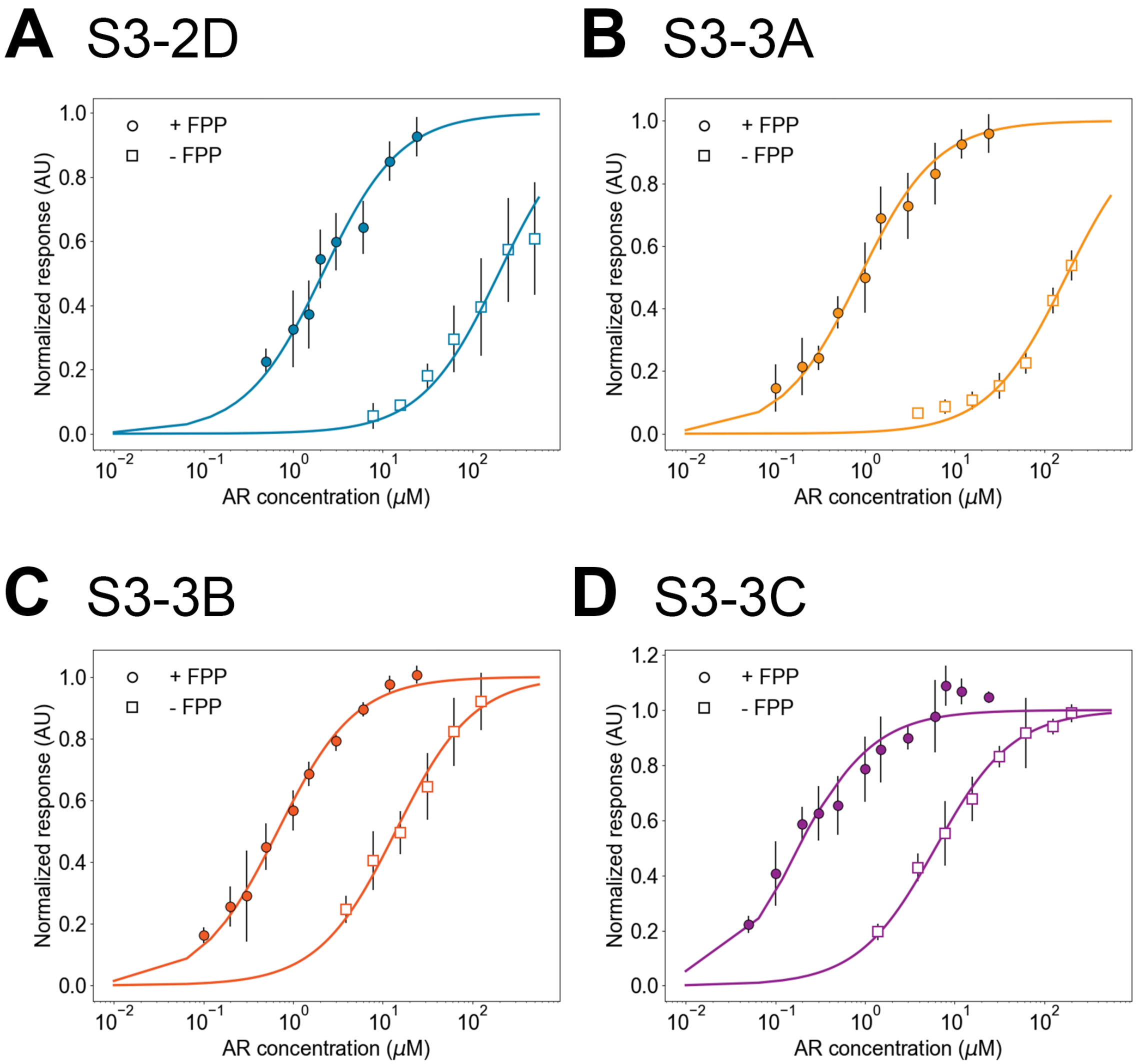

**Fig. S13. BLI binding data +/- FPP for designs S3-2D, S3-3A, S3-3B, S3-3C.** Experiments use immobilized MBP variants titrating in AR variants at the indicated concentrations. Data for the +FPP condition are also shown in **Figure 3E** and depicted here for comparison. Apparent $K_D$ values are given in **Figure 3C**. Error bars are standard deviations, $n \geq 3$ for each of +/- FPP conditions.

**Table S1. Summary of computational designs.** The design ID lists scaffold first (S1, S2, S3), followed by design round (1, 2, 3), followed by a letter (A, B, etc). The nomenclature for the individual sensor proteins lists protein, design round, and a consecutive number denoting the protein's variant.

| **Scaffold (WT PDB)** | **Design ID** | **Sensor Protein A** | **Sensor Protein B** | **Design / Engineering Method** |
|---|---|---|---|---|
| **Scaffold 1:** FKBP12 -FRB (3FAP.pdb) | S1-1A | FKBP12-1.1 | FRB-1.1 | Computational Design |
| **Scaffold 2:** RapF-ComA (3ULQ.pdb) | S2-1A | RapF-1.1 | ComA-1.1 | Computational Design |
| | S2-1B | RapF-1.2 | ComA-1.2 | Computational Design |
| | S2-1C | RapF-1.3 | ComA-1.3 | Computational Design |
| | S2-1D | RapF-1.4 | ComA-1.4 | Computational Design |
| **Scaffold 3:** AR-MBP (1SVX.pdb) | **Computational designs (group 1)** | | | |
| | S3-1A | AR-1.1 | MBP-1.1 | Computational Design, best ligand burial |
| | S3-1B | AR-1.2 | MBP-1.2 | Computational Design, consensus |
| | S3-1C | AR-1.3 | MBP-1.3 | Computational Design, optimized ligand packing |
| | S3-1D | AR-1.4 | MBP-1.4 | Computational Design, best ligand score |
| | **Computational designs with stability enhancing mutations (group 2)** | | | |
| | S3-2A | AR-2.5 | MBP-1.3 | Hit from error-prone PCR (identical to design S3-1C except for 2 mutations in AR: N102H, A104E) |
| | S3-2B | AR-2.5 | MBP-2.5 | S3-2A with stability enhancing mutation from saturation mutagenesis in MBP: R194A |
| | S3-2C | AR-2.6 | MBP-2.5 | S3-2A with stability enhancing mutations from saturation mutagenesis: MBP: R194A AR: L85G |
| | S3-2D | AR-2.7 | MBP-2.5 | S3-2C with published stabilizing mutations in AR*: W122K, E152L, A149P, S153A, L161I, I164V, L168A, N169A |
| | **Computational designs with designed affinity enhancing mutations (group 3)** | | | |
| | S3-3A | AR-2.7 | MBP-3.6 | S3-2D with affinity enhancing mutation in MBP from KIC ensemble design: Y197A |
| | S3-3B | AR-3.8 | MBP-2.5 | S3-2D with affinity enhancing mutations in AR from CoupledMoves: R145K, K147L, D155L |
| | S3-3C | AR-3.8 | MBP-3.6 | S3-2D with affinity enhancing mutations in AR and MBP: AR: R145K, K147L, D155L MBP: Y197A |

* from reference (*33*)

**Table S2. Computationally designed library.** We designed a sequence library with $2.4\times10^6$ members using degenerate codons (top) based on the dominant residues predicted computationally (bottom). The library was tested using the plate-based split mDHFR assay. Hits were defined as sensor sequences that grew better in the presence of mevalonate. Residues are shown with motif positions in red and wild-type residues shaded grey. Residues with minor frequencies in the flexible design predictions are shown as lower-case letters.

| Sequence position | 81 | 85 | 89 | 114 | 119 | 122 | 123 | 145 | 152 | 133 | 194 | 197 | 200 | 201 | 203 |
|---|---|---|---|---|---|---|---|---|---|---|---|---|---|---|---|
| Protein | AR | | | | | | | | | MBP | | | | | |
| Wild-type residue | Y | H | Y | M | L | K | W | F | I | P | F | D | K | N | H |
| Library codon | WTK | WTT | WTK | TGG | GCG | KHT | BHK | CGT | RYT | YWT | – | GVC | RMG | KHT | – |
| Library residues (plus F79A) | F<br>I<br>L<br>M | F<br>I | F<br>I<br>L<br>M | W | A | A<br>D<br>F<br>S<br>V<br>Y | F<br>Y<br>A<br>S<br>V<br>D<br>L<br>P<br>H<br>Q<br>E | R | A<br>I<br>T<br>V | F<br>H<br>L<br>Y | – | A<br>D<br>G | A<br>E<br>K<br>T | A<br>D<br>F<br>S<br>V<br>Y | – |
| Residues in library hits | I<br>L<br>M | F<br>I | F<br>M | W | A | D<br>S<br>V<br>Y | Y<br>A<br>V<br>L<br>E | R | I<br>T<br>V | F<br>Y<br>H | – | D<br>A<br>G | K<br>E<br>A | N<br>A<br>D | – |

**Computational predictions**

| Sequence position | 81 | 85 | 89 | 114 | 119 | 122 | 123 | 145 | 152 | 133 | 194 | 197 | 200 | 201 | 203 |
|---|---|---|---|---|---|---|---|---|---|---|---|---|---|---|---|
| Protein | AR | | | | | | | | | MBP | | | | | |
| Computational Design (Top-ranked individual Sequences) (+99: D,G,T) | F<br>I<br>L<br>M | F<br>I<br>L | L | W | A | Y<br>W | F<br>Y | R | I<br>V<br>E | F<br>Y | F<br>R | A<br>Y | A<br>K | A<br>S | A<br>H<br>S |
| Computational Predictions (KIC Ensemble) | a<br>F<br>i<br>L<br>M | a<br>F<br>I<br>m | L | W | A | A<br>F<br>l<br>Y | A<br>e<br>F<br>h<br>q<br>Y | R | A<br>V | F<br>Y | – | A<br>G | A | A<br>L<br>F<br>Y | – |
| Computational Predictions (CoupledMoves) | F | F<br>H<br>L<br>M | L<br>M | A<br>F<br>H<br>V<br>W | A | F<br>W<br>Y | F<br>M<br>Y | E<br>R<br>N<br>T | H<br>F<br>Y | F<br>Y | F<br>W | A<br>F<br>H<br>Y | A | A<br>S | – |

**Table S3: Single-site saturation mutagenesis positions and results.**

| Sequence position | 85 | 112 | 118 | 119 | 122 | 123 | 152 | 155 | 194 | 197 | 251 |
|---|---|---|---|---|---|---|---|---|---|---|---|
| Protein | AR | | | | | | | | MBP | | |
| Wild-type residue | H | D | H | L | K | W | I | D | F | D | K |
| Residues in library hits | F I | Not varied in comp. library design | Not varied in comp. library design | A | D S V Y | Y A V L E | I T V | Not varied in comp. library design | Not varied in comp. library design | D A G | Not varied in comp. library design |
| Residue in design S3-1D | F | D | H | A | Y | Y | V | D | F | A | K |
| Residue in design S3-1C | L | D | H | A | W | F | E | D | R | Y | K |
| Residue in design S3-2A | L | D | H | A | W | F | E | D | R | Y | K |
| Saturation mutagenesis hits round 1, starting from S3-2A | - | - | L N P | - | V L | - | - | - | A G F | - | - |
| Best from combination of hits round 1 (design S3-2B) | L | D | H | A | W | F | E | D | A | Y | K |
| Saturation mutagenesis hits round 2, starting from S3-2B | G | - | T I | P R | - | - | - | - | - | - | - |
| Best from round 2 (design S3-2C) | G | D | H | A | W | F | E | D | A | Y | K |

**Table S4: X-ray data reduction and model refinement.**

| | |
|---|---|
| Wavelength | 1.116Å |
| Resolution Range | 95.47-2.20 (2.24-2.20) |
| Unit Cell | a=44.57Å , b=190.92Å , c=55.48Å<br>**α**=**β**=**γ**=90° |
| Space Group | $P2_1$ |
| Unique Reflections | 45972 (7746) |
| Multiplicity | 3.7 (3.6) |
| Completeness | 98.5% (91.5%) |
| <I/σI> | 17.5 (8.3) |
| $CC_{1/2}$[a] | 0.997 (0.966) |
| $R_{pim}$[b] | 0.033 (0.090) |
| $R_{work}$[c] | 0.2071 (0.3448) |
| $R_{free}$[c] | 0.2596 (0.4761) |
| Total Refined Atoms | 8115 |
| Protein Residues | 1022 |
| Solvent Molecules | 141 |
| Refined Ligand Atoms | 70 |
| Average B-factor | 21.6Å$^2$ |
| $RMSD_{bonds}$ | 0.003Å |
| $RMSD_{angles}$ | 0.56° |
| Rama. Plot: | |
| Favored | 97.1% |
| Allowed | 2.9% |
| Outliers | 0.0% |
| Molprobity Clashscore[d] | 3.97 |
| PDB ID | 6OB5 |

a. Reference (*34*)
b. Reference (*20*)
c. Reference (*35*)
d. Reference (*36*)

**Appendix 1: Plasmids / constructs for split mDHFR reporter assays.** See **Appendix 4** for full DNA sequences for each gene and vector listed below.

**Designs with C-terminal split mDHFR**

| Construct | Vector background | Description |
|---|---|---|
| **DHFR1** | pCDFDuet* | FKBP12/FRB design S1-1A |
| **DHFR2** | pCDFDuet | RapF/ComA design S2-1A |
| **DHFR3** | pCDFDuet | RapF/ComA design S2-1B |
| **DHFR4** | pCDFDuet | RapF/ComA design S2-1C |
| **DHFR5** | pCDFDuet | RapF/ComA design S2-1D |
| **DHFR6** | pCDFDuet | AR/MBP design S3-1A |
| **DHFR7** | pCDFDuet | AR/MBP design S3-1B |
| **DHFR8** | pCDFDuet | AR/MBP design S3-1C |
| **DHFR9** | pCDFDuet | AR/MBP design S3-1D |
| **DHFR10** | pCDFDuet | AR/MBP Design S3-2A |
| **DHFR11** | pCDFDuet | AR/MBP Design S3-2B |
| **DHFR12** | pCDFDuet | AR/MBP Design S3-2C |
| **DHFR13** | pCDFDuet | AR/MBP Design S3-2D |
| **DHFR14** | pCDFDuet | AR/MBP Design S3-3A |

***** pCDFDuet (Novagen) is spectromycin/spectinomycin-resistant and has the CDF origin of replication. Each protein is fused to one of the subunits of split mDHFR. See **Appendix 3** for sequences of split mDHFR subunits.

**Mevalonate pathway constructs**

| Construct | Vector background | Description |
|---|---|---|
| **pMBIS** | pB8a** | *ERG12-ERG8-MVD1-idi-ispA* |
| **ispA R116A** | pB8a | pMBIS + loss of function in *ispA* |
| **pB5K** | pB8a | pMBIS + *ADS* |

****** pB8a is **chloramphenicol-resistant** and has the **pBBR1 origin of replication**.

**Motif residue alanine reversions with C-terminal split mDHFR**

| Construct | Vector background | Description |
|---|---|---|
| **DHFR15** | pCDFDuet | Design S3-2C AR L89A |
| **DHFR16** | pCDFDuet | Design S3-2C AR W114A |
| **DHFR17** | pCDFDuet | Design S3-2C AR R145A |
| **DHFR18** | pCDFDuet | Design S3-2C MBP F133A |

**Appendix 2: Plasmids / constructs for *in vitro* binding experiments and crystallography.** These constructs included the C-terminal alpha helix in MBP (residues 354-370). See **Appendix 4** for full DNA sequences for each gene and vector listed below.

**Constructs for bio-layer interferometry (Designs S3-2D, S3-3A, S3-3B and S3-3C)**

| **Construct** | **Vector background** | **Description*** |
|---|---|---|
| **BLI1** | pET28b-GG | AR-2.7 |
| **BLI2** | pET28b-GG | Avi-MBP-2.5 |
| **BLI3** | pET28b-GG | AR-3.8 |
| **BLI4** | pET28b-GG | Avi-MBP-3.6 |

***Avi tag sequence**
**DNA: GGTCTGAACGACATCTTCGAGGCTCAGAAAATCGAATGGCACGAA**
**Protein: GLNDIFEAQKIEWHE**

**Constructs for crystallography (Design S3-2D)**

| **Construct** | **Vector background** | **Description** |
|---|---|---|
| **CR1** | pET28a | MBP-2.5 |
| **CR2** | pET47b(+) | AR-2.7 |

**Appendix 3: Plasmids / constructs for TxTl reporter assays.** See **Appendix 4** for full DNA sequences for each gene, vector, and reporter listed below.

**Designs S3-2D and S3-3A constructs for TxTl cell-free reporter assay**

| Construct | Vector background* | Description** |
|---|---|---|
| **TxTl1** | TxTl T7 A1 | LgBIT-AR-2.7 |
| **TxTl2** | TxTl T7 A2 | MBP-2.5-SmBIT |
| **TxTl3** | TxTl T7 A2 | ddRFPb-AR-2.7 |
| **TxTl4** | TxTl T7 A2 | MBP-2.5-ddGFPa |
| **TxTl5** | TxTl T7 A2 | MBP-3.6-SmBIT |
| **TxTl6** | TxTl T7 A2 | MBP-3.6-ddGFPa |

* The TxTl T7 A1 vector has a kanamycin resistance cassette and a p15a origin of replication.
The TxTl T7 A2 vector has an ampicillin resistance cassette and a ColE1 origin of replication.

**LgBIT: 18 kDa large subunit of the engineered split nanoluciferase, NanoBiT
SmBIT: 1.3 kDa (11 amino acid peptide) small subunit of NanoBiT

ddRFPb: 26 kDa dimerization-dependent protein that fluoresces in complex with ddGFPa.
ddGFPa: 26 kDa dimerization-dependent protein that fluoresces in complex with ddRFPb.

ddRFPb + ddGFPa complex is excited at 493 and 380 nm, with emission peaks at 448 and 508 nm.

**Controls for TxTl cell-free reporter assay**

| Construct | Vector background | Description |
|---|---|---|
| **TxTl7** | TxTl T7 A1 | LgBIT-WT AR |
| **TxTl8** | TxTl T7 A2 | WT MBP-SmBIT |
| **TxTl9** | TxTl T7 A2 | ddRFPb-WT AR |
| **TxTl10** | TxTl T7 A2 | WT MBP-ddGFPa |

**Additional construct**

| Construct | Vector background*** | Description |
|---|---|---|
| **TxTl11** | TxTl T500 A3 | T7 RNA polymerase expression |

***The TxTl T500 A3 vector has an ampicillin resistance cassette and a p15a origin of replication. The target gene is expressed from the Pr promoter from lambda phage.

### Appendix 4: Gene and vector DNA sequences.

#### pCDFDuet

CGCTGACGTCGGTACCCTCGAGTCTGGTAAAGAAACCGCTGCTGCGAAATTTGAACGCCAGCACATGGAC
TCGTCTACTAGCGCAGCTTAATTAACCTAGGCTGCTGCCACCGCTGAGCAATAACTAGCATAACCCCTTG
GGGCCTCTAAACGGGTCTTGAGGGGTTTTTTGCTGAAACCTCAGGCATTTGAGAAGCACACGGTCACACT
GCTTCCGGTAGTCAATAAACCGGTAAACCAGCAATAGACATAAGCGGCTATTTAACGACCCTGCCCTGAA
CCGACGACCGGGTCATCGTGGCCGGATCTTGCGGCCCCTCGGCTTGAACGAATTGTTAGACATTATTTGC
CGACTACCTTGGTGATCTCGCCTTTCACGTAGTGGACAAATTCTTCCAACTGATCTGCGCGCGAGGCCAA
GCGATCTTCTTCTTGTCCAAGATAAGCCTGTCTAGCTTCAAGTATGACGGGCTGATACTGGGCCGGCAGG
CGCTCCATTGCCCAGTCGGCAGCGACATCCTTCGGCGCGATTTTGCCGGTTACTGCGCTGTACCAAATGC
GGGACAACGTAAGCACTACATTTCGCTCATCGCCAGCCCAGTCGGGCGGCGAGTTCCATAGCGTTAAGGT
TTCATTTAGCGCCTCAAATAGATCCTGTTCAGGAACCGGATCAAAGAGTTCCTCCGCCGCTGGACCTACC
AAGGCAACGCTATGTTCTCTTGCTTTTGTCAGCAAGATAGCCAGATCAATGTCGATCGTGGCTGGCTCGA
AGATACCTGCAAGAATGTCATTGCGCTGCCATTCTCCAAATTGCAGTTCGCGCTTAGCTGGATAACGCCA
CGGAATGATGTCGTCGTGCACAACAATGGTGACTTCTACAGCGCGGAGAATCTCGCTCTCTCCAGGGGAA
GCCGAAGTTTCCAAAAGGTCGTTGATCAAAGCTCGCCGCGTTGTTTCATCAAGCCTTACGGTCACCGTAA
CCAGCAAATCAATATCACTGTGTGGCTTCAGGCCGCCATCCACTGCGGAGCCGTACAAATGTACGGCCAG
CAACGTCGGTTCGAGATGGCGCTCGATGACGCCAACTACCTCTGATAGTTGAGTCGATACTTCGGCGATC
ACCGCTTCCCTCATACTCTTCCTTTTTCAATATTATTGAAGCATTTATCAGGGTTATTGTCTCATGAGCG
GATACATATTTGAATGTATTTAGAAAAATAAACAAATAGCTAGCTCACTCGGTCGCTACGCTCCGGGCGT
GAGACTGCGGCGGGCGCTGCGGACACATACAAAGTTACCCACAGATTCCGTGGATAAGCAGGGGACTAAC
ATGTGAGGCAAAACAGCAGGGCCGCGCCGGTGGCGTTTTTCCATAGGCTCCGCCCTCCTGCCAGAGTTCA
CATAAACAGACGCTTTTCCGGTGCATCTGTGGGAGCCGTGAGGCTCAACCATGAATCTGACAGTACGGGC
GAAACCCGACAGGACTTAAAGATCCCCACCGTTTCCGGCGGGTCGCTCCCTCTTGCGCTCTCCTGTTCCG
ACCCTGCCGTTTACCGGATACCTGTTCCGCCTTTCTCCCTTACGGGAAGTGTGGCGCTTTCTCATAGCTC
ACACACTGGTATCTCGGCTCGGTGTAGGTCGTTCGCTCCAAGCTGGGCTGTAAGCAAGAACTCCCCGTTC
AGCCCGACTGCTGCGCCTTATCCGGTAACTGTTCACTTGAGTCCAACCCGGAAAAGCACGGTAAAACGCC
ACTGGCAGCAGCCATTGGTAACTGGGAGTTCGCAGAGGATTTGTTTAGCTAAACACGCGGTTGCTCTTGA
AGTGTGCGCCAAAGTCCGGCTACACTGGAAGGACAGATTTGGTTGCTGTGCTCTGCGAAAGCCAGTTACC
ACGGTTAAGCAGTTCCCCAACTGACTTAACCTTCGATCAAACCACCTCCCCAGGTGGTTTTTTCGTTTAC
AGGGCAAAAGATTACGCGCAGAAAAAAAGGATCTCAAGAAGATCCTTTGATCTTTTCTACTGAACCGCTC
TAGATTTCAGTGCAATTTATCTCTTCAAATGTAGCACCTGAAGTCAGCCCCATACGATATAAGTTGTAAT
TCTCATGTTAGTCATGCCCCGCGCCCACCGGAAGGAGCTGACTGGGTTGAAGGCTCTCAAGGGCATCGGT
CGAGATCCCGGTGCCTAATGAGTGAGCTAACTTACATTAATTGCGTTGCGCTCACTGCCCGCTTTCCAGT
CGGGAAACCTGTCGTGCCAGCTGCATTAATGAATCGGCCAACGCGCGGGGAGAGGCGGTTTGCGTATTGG
GCGCCAGGGTGGTTTTTCTTTTCACCAGTGAGACGGGCAACAGCTGATTGCCCTTCACCGCCTGGCCCTG
AGAGAGTTGCAGCAAGCGGTCCACGCTGGTTTGCCCCAGCAGGCGAAAATCCTGTTTGATGGTGGTTAAC
GGCGGGATATAACATGAGCTGTCTTCGGTATCGTCGTATCCCACTACCGAGATGTCCGCACCAACGCGCA
GCCCGGACTCGGTAATGGCGCGCATTGCGCCCAGCGCCATCTGATCGTTGGCAACCAGCATCGCAGTGGG
AACGATGCCCTCATTCAGCATTTGCATGGTTTGTTGAAAACCGGACATGGCACTCCAGTCGCCTTCCCGT
TCCGCTATCGGCTGAATTTGATTGCGAGTGAGATATTTATGCCAGCCAGCCAGACGCAGACGCGCCGAGA
CAGAACTTAATGGGCCCGCTAACAGCGCGATTTGCTGGTGACCCAATGCGACCAGATGCTCCACGCCCAG
TCGCGTACCGTCTTCATGGGAGAAAATAATACTGTTGATGGGTGTCTGGTCAGAGACATCAAGAAATAAC
GCCGGAACATTAGTGCAGGCAGCTTCCACAGCAATGGCATCCTGGTCATCCAGCGGATAGTTAATGATCA
GCCCACTGACGCGTTGCGCGAGAAGATTGTGCACCGCCGCTTTACAGGCTTCGACGCCGCTTCGTTCTAC
CATCGACACCACCACGCTGGCACCCAGTTGATCGGCGCGAGATTTAATCGCCGCGACAATTTGCGACGGC
GCGTGCAGGGCCAGACTGGAGGTGGCAACGCCAATCAGCAACGACTGTTTGCCCGCCAGTTGTTGTGCCA
CGCGGTTGGGAATGTAATTCAGCTCCGCCATCGCCGCTTCCACTTTTTCCCGCGTTTTCGCAGAAACGTG
GCTGGCCTGGTTCACCACGCGGGAAACGGTCTGATAAGAGACACCGGCATACTCTGCGACATCGTATAAC

GTTACTGGTTTCACATTCACCACCCTGAATTGACTCTCTTCCGGGCGCTATCATGCCATACCGCGAAAGG
TTTTGCGCCATTCGATGGTGTCCGGGATCTCGACGCTCTCCCTTATGCGACTCCTGCATTAGGAAATTAA
TACGACTCACTATAGGGGAATTGTGAGCGGATAACAATTCCCCTGTAGAAATAATTTTGTTTAACTTTAA
TAAGGAGATATACC

**pB8a**

GGATCCTAACTCGAGTAAGGATCTCCAGGCATCAAATAAAACGAAAGGCTCAGTCGAAAGACTGGGCCTT
TCGTTTTATCTGTTGTTTGTCGGTGAACGCTCTCTACTAGAGTCACACTGGCTCACCTTCGGGTGGGCCT
TTTGCGTTTATACCTAGGCTACAGCCGATAGTCTGGAACAGCGCACTTACGGGTTGCTGCGCAACCCAAG
TGCTACCGGCGCGGCAGCGTGACCCGTGTCGGCGGCTCCAACGGCTCGCCATCGTCCAGAAAACACGGCT
CATCGGGCATCGGCAGGCGCTGCTGCCCGCGCCGTTCCCATTCCTCCGTTTCGGTCAAGGCTGGCAGGTC
TGGTTCCATGCCCGGAATGCCGGGCTGGCTGGGCGGCTCCTCGCCGGGGCCGGTCGGTAGTTGCTGCTCG
CCCGGATACAGGGTCGGGATGCGGCGCAGGTCGCCATGCCCCAACAGCGATTCGTCCTGGTCGTCGTGAT
CAACCACCACGGCGGCACTGAACACCGACAGGCGCAACTGGTCGCGGGGCTGGCCCCACGCCACGCGGTC
ATTGACCACGTAGGCCAACACGGTGCCGGGGCCGTTGAGCTTCACGACGGAGATCCAGCGCTCGGCCACC
AAGTCCTTGACTGCGTATTGGACCGTCCGCAAAGAACGTCCGATGAGCTTGGAAAGTGTCTTCTGGCTGA
CCACCACGGCGTTCTGGTGGCCCATCTGCGCCACGAGGTGATGCAGCAGCATTGCCGCCGTGGGTTTCCT
CGCAATAAGCCCGGCCCACGCCTCATGCGCTTTGCGTTCCGTTTGCACCCAGTGACCGGGCTTGTTCTTG
GCTTGAATGCCGATTTCTCTGGACTGCGTGGCCATGCTTATCTCCATGCGGTAGGGGTGCCGCACGGTTG
CGGCACCATGCGCAATCAGCTGCAACTTTTCGGCAGCGCGACAACAATTATGCGTTGCGTAAAAGTGGCA
GTCAATTACAGATTTTCTTTAACCTACGCAATGAGCTATTGCGGGGGGTGCCGCAATGAGCTGTTGCGTA
CCCCCCTTTTTTAAGTTGTTGATTTTTAAGTCTTTCGCATTTCGCCCTATATCTAGTTCTTTGGTGCCCA
AAGAAGGGCACCCCTGCGGGGTTCCCCCACGCCTTCGGCGCGGCTCCCCCTCCGGCAAAAAGTGGCCCCT
CCGGGGCTTGTTGATCGACTGCGCGGCCTTCGGCCTTGCCCAAGGTGGCGCTGCCCCCTTGGAACCCCCG
CACTCGCCGCCGTGAGGCTCGGGGGGCAGGCGGGCGGGCTTCGCCCTTCGACTGCCCCCACTCGCATAGG
CTTGGGTCGTTCCAGGCGCGTCAAGGCCAAGCCGCTGCGCGGTCGCTGCGCGAGCCTTGACCCGCCTTCC
ACTTGGTGTCCAACCGGCAAGCGAAGCGCGCAGGCCGCAGGCCGGAGGCACTAGTGCTTGGATTCTCACC
AATAAAAAACGCCCGGCGGCAACCGAGCGTTCTGAACAAATCCAGATGGAGTTCTGAGGTCATTACTGGA
TCTATCAACAGGAGTCCAAGCGAGCTCGTAAACTTGGTCTGACAGTTACGCCCCGCCCTGCCACTCATCG
CAGTACTGTTGTAATTCATTAAGCATTCTGCCGACATGGAAGCCATCACAAACGGCATGATGAACCTGAA
TCGCCAGCGGCATCAGCACCTTGTCGCCTTGCGTATAATATTTGCCCATGGTGAAAACGGGGGCGAAGAA
GTTGTCCATATTGGCCACGTTTAAATCAAAACTGGTGAAACTCACCCAGGGATTGGCTGAGACGAAAAAC
ATATTCTCAATAAACCCTTTAGGGAAATAGGCCAGGTTTTCACCGTAACACGCCACATCTTGCGAATATA
TGTGTAGAAACTGCCGGAAATCGTCGTGGTATTCACTCCAGAGCGATGAAAACGTTTCAGTTTGCTCATG
GAAAACGGTGTAACAAGGGTGAACACTATCCCATATCACCAGCTCACCGTCTTTCATTGCCATACGGAAT
TCCGGATGAGCATTCATCAGGCGGGCAAGAATGTGAATAAAGGCCGGATAAAACTTGTGCTTATTTTTCT
TTACGGTCTTTAAAAAGGCCGTAATATCCAGCTGAACGGTCTGGTTATAGGTACATTGAGCAACTGACTG
AAATGCCTCAAAATGTTCTTTACGATGCCATTGGGATATATCAACGGTGGTATATCCAGTGATTTTTTTC
TCCATACTCTTCCTTTTTCAATATTATTGAAGCATTTATCAGGGTTATTGTCTCATGAGCGGATACATAT
TTGAATGTATTTAGAAAAATAAACAAATAGGGGTTCCGCGCACATTTCCCCGAAAAGTGCCACCTGACGT
C

**pET28b-GG (with GFP dropout)**

TGGCGAATGGGACGCGCCCTGTAGCGGCGCATTAAGCGCGGCGGGTGTGGTGGTTACGCGCAGCGTGACC
GCTACACTTGCCAGCGCCCTAGCGCCCGCTCCTTTCGCTTTCTTCCCTTCCTTTCTCGCCACGTTCGCCG
GCTTTCCCCGTCAAGCTCTAAATCGGGGGCTCCCTTTAGGGTTCCGATTTAGTGCTTTACGGCACCTCGA
CCCCAAAAAACTTGATTAGGGTGATGGTTCACGTAGTGGGCCATCGCCCTGATAGACGGTTTTTCGCCCT
TTGACGTTGGAGTCCACGTTCTTTAATAGTGGACTCTTGTTCCAAACTGGAACAACACTCAACCCTATCT
CGGTCTATTCTTTTGATTTATAAGGGATTTTGCCGATTTCGGCCTATTGGTTAAAAAATGAGCTGATTTA
ACAAAAATTTAACGCGAATTTTAACAAAATATTAACGTTTACAATTTCAGGTGGCACTTTTCGGGGAAAT

GTGCGCGGAACCCCTATTTGTTTATTTTTCTAAATACATTCAAATATGTATCCGCTCATGAATTAATTCT
TAGAAAAACTCATCGAGCATCAAATGAAACTGCAATTTATTCATATCAGGATTATCAATACCATATTTTT
GAAAAAGCCGTTTCTGTAATGAAGGAGAAAACTCACCGAGGCAGTTCCATAGGATGGCAAGATCCTGGTA
TCGGTCTGCGATTCCGACTCGTCCAACATCAATACAACCTATTAATTTCCCCTCGTCAAAAATAAGGTTA
TCAAGTGAGAAATCACCATGAGTGACGACTGAATCCGGTGAGAATGGCAAAAGTTTATGCATTTCTTTCC
AGACTTGTTCAACAGGCCAGCCATTACGCTCGTCATCAAAATCACTCGCATCAACCAAACCGTTATTCAT
TCGTGATTGCGCCTGAGCGAGACGAAATACGCGATCGCTGTTAAAAGGACAATTACAAACAGGAATCGAA
TGCAACCGGCGCAGGAACACTGCCAGCGCATCAACAATATTTTCACCTGAATCAGGATATTCTTCTAATA
CCTGGAATGCTGTTTTCCCGGGGATCGCAGTGGTGAGTAACCATGCATCATCAGGAGTACGGATAAAATG
CTTGATGGTCGGAAGAGGCATAAATTCCGTCAGCCAGTTTAGTCTGACCATCTCATCTGTAACATCATTG
GCAACGCTACCTTTGCCATGTTTCAGAAACAACTCTGGCGCATCGGGCTTCCCATACAATCGATAGATTG
TCGCACCTGATTGCCCGACATTATCGCGAGCCCATTTATACCCATATAAATCAGCATCCATGTTGGAATT
TAATCGCGGCCTAGAGCAAGACGTTTCCCGTTGAATATGGCTCATAACACCCCTTGTATTACTGTTTATG
TAAGCAGACAGTTTTATTGTTCATGACCAAAATCCCTTAACGTGAGTTTTCGTTCCACTGAGCGTCAGAC
CCCGTAGAAAAGATCAAAGGATCTTCTTGAGATCCTTTTTTTCTGCGCGTAATCTGCTGCTTGCAAACAA
AAAAACCACCGCTACCAGCGGTGGTTTGTTTGCCGGATCAAGAGCTACCAACTCTTTTTCCGAAGGTAAC
TGGCTTCAGCAGAGCGCAGATACCAAATACTGTCCTTCTAGTGTAGCCGTAGTTAGGCCACCACTTCAAG
AACTCTGTAGCACCGCCTACATACCTCGCTCTGCTAATCCTGTTACCAGTGGCTGCTGCCAGTGGCGATA
AGTCGTGTCTTACCGGGTTGGACTCAAGACGATAGTTACCGGATAAGGCGCAGCGGTCGGGCTGAACGGG
GGGTTCGTGCACACAGCCCAGCTTGGAGCGAACGACCTACACCGAACTGAGATACCTACAGCGTGAGCTA
TGAGAAAGCGCCACGCTTCCCGAAGGGAGAAAGGCGGACAGGTATCCGGTAAGCGGCAGGGTCGGAACAG
GAGAGCGCACGAGGGAGCTTCCAGGGGGAAACGCCTGGTATCTTTATAGTCCTGTCGGGTTTCGCCACCT
CTGACTTGAGCGTCGATTTTTGTGATGCTCGTCAGGGGGGCGGAGCCTATGGAAAAACGCCAGCAACGCG
GCCTTTTTACGGTTCCTGGCCTTTTGCTGGCCTTTTGCTCACATGTTCTTTCCTGCGTTATCCCCTGATT
CTGTGGATAACCGTATTACCGCCTTTGAGTGAGCTGATACCGCTCGCCGCAGCCGAACGACCGAGCGCAG
CGAGTCAGTGAGCGAGGAAGCGGAAGAGCGCCTGATGCGGTATTTTCTCCTTACGCATCTGTGCGGTATT
TCACACCGCATATATGGTGCACTCTCAGTACAATCTGCTCTGATGCCGCATAGTTAAGCCAGTATACACT
CCGCTATCGCTACGTGACTGGGTCATGGCTGCGCCCCGACACCCGCCAACACCCGCTGACGCGCCCTGAC
GGGCTTGTCTGCTCCCGGCATCCGCTTACAGACAAGCTGTGACCGTCTCCGGGAGCTGCATGTGTCAGAG
GTTTTCACCGTCATCACCGAAACGCGCGAGGCAGCTGCGGTAAAGCTCATCAGCGTGGTCGTGAAGCGAT
TCACAGATGTCTGCCTGTTCATCCGCGTCCAGCTCGTTGAGTTTCTCCAGAAGCGTTAATGTCTGGCTTC
TGATAAAGCGGGCCATGTTAAGGGCGGTTTTTTCCTGTTTGGTCACTGATGCCTCCGTGTAAGGGGGATT
TCTGTTCATGGGGGTAATGATACCGATGAAACGAGAGAGGATGCTCACGATACGGGTTACTGATGATGAA
CATGCCCGGTTACTGGAACGTTGTGAGGGTAAACAACTGGCGGTATGGATGCGGCGGGACCAGAGAAAAA
TCACTCAGGGTCAATGCCAGCGCTTCGTTAATACAGATGTAGGTGTTCCACAGGGTAGCCAGCAGCATCC
TGCGATGCAGATCCGGAACATAATGGTGCAGGGCGCTGACTTCCGCGTTTCCAGACTTTACGAAACACGG
AAACCGAAGACCATTCATGTTGTTGCTCAGGTCGCAGACGTTTTGCAGCAGCAGTCGCTTCACGTTCGCT
CGCGTATCGGTGATTCATTCTGCTAACCAGTAAGGCAACCCCGCCAGCCTAGCCGGGTCCTCAACGACAG
GAGCACGATCATGCGCACCCGTGGGGCCGCCATGCCGGCGATAATGGCCTGCTTCTCGCCGAAACGTTTG
GTGGCGGGACCAGTGACGAAGGCTTGAGCGAGGGCGTGCAAGATTCCGAATACCGCAAGCGACAGGCCGA
TCATCGTCGCGCTCCAGCGAAAGCGGTCCTCGCCGAAAATGACCCAGAGCGCTGCCGGCACCTGTCCTAC
GAGTTGCATGATAAAGAAGACAGTCATAAGTGCGGCGACGATAGTCATGCCCCGCGCCCACCGGAAGGAG
CTGACTGGGTTGAAGGCTCTCAAGGGCATCGGTCGAGATCCCGGTGCCTAATGAGTGAGCTAACTTACAT
TAATTGCGTTGCGCTCACTGCCCGCTTTCCAGTCGGGAAACCTGTCGTGCCAGCTGCATTAATGAATCGG
CCAACGCGCGGGGAGAGGCGGTTTGCGTATTGGGCGCCAGGGTGGTTTTTCTTTTCACCAGTGAGACGGG
CAACAGCTGATTGCCCTTCACCGCCTGGCCCTGAGAGAGTTGCAGCAAGCGGTCCACGCTGGTTTGCCCC
AGCAGGCGAAAATCCTGTTTGATGGTGGTTAACGGCGGGATATAACATGAGCTGTCTTCGGTATCGTCGT
ATCCCACTACCGAGATATCCGCACCAACGCGCAGCCCGGACTCGGTAATGGCGCGCATTGCGCCCAGCGC
CATCTGATCGTTGGCAACCAGCATCGCAGTGGGAACGATGCCCTCATTCAGCATTTGCATGGTTTGTTGA
AAACCGGACATGGCACTCCAGTCGCCTTCCCGTTCCGCTATCGGCTGAATTTGATTGCGAGTGAGATATT
TATGCCAGCCAGCCAGACGCAGACGCGCCGAGACAGAACTTAATGGGCCCGCTAACAGCGCGATTTGCTG
GTGACCCAATGCGACCAGATGCTCCACGCCCAGTCGCGTACCGTCTTCATGGGAGAAAATAATACTGTTG

ATGGGTGTCTGGTCAGAGACATCAAGAAATAACGCCGGAACATTAGTGCAGGCAGCTTCCACAGCAATGG
CATCCTGGTCATCCAGCGGATAGTTAATGATCAGCCCACTGACGCGTTGCGCGAGAAGATTGTGCACCGC
CGCTTTACAGGCTTCGACGCCGCTTCGTTCTACCATCGACACCACCACGCTGGCACCCAGTTGATCGGCG
CGAGATTTAATCGCCGCGACAATTTGCGACGGCGCGTGCAGGGCCAGACTGGAGGTGGCAACGCCAATCA
GCAACGACTGTTTGCCCGCCAGTTGTTGTGCCACGCGGTTGGGAATGTAATTCAGCTCCGCCATCGCCGC
TTCCACTTTTTCCCGCGTTTTCGCAGAAACGTGGCTGGCCTGGTTCACCACGCGGGAAACGGTCTGATAA
GAGACACCGGCATACTCTGCGACATCGTATAACGTTACTGGTTTCACATTCACCACCCTGAATTGACTCT
CTTCCGGGCGCTATCATGCCATACCGCGAAAGGTTTTGCGCCATTCGATGGTGTCCGGGATCTCGACGCT
CTCCCTTATGCGACTCCTGCATTAGGAAGCAGCCCAGTAGTAGGTTGAGGCCGTTGAGCACCGCCGCCGC
AAGGAATGGTGCATGCAAGGAGATGGCGCCCAACAGTCCCCCGGCCACGGGGCCTGCCACCATACCCACG
CCGAAACAAGCGCTCATGAGCCCGAAGTGGCGAGCCCGATCTTCCCCATCGGTGATGTCGGCGATATAGG
CGCCAGCAACCGCACCTGTGGCGCCGGTGATGCCGGCCACGATGCGTCCGGCGTAGAGGATCGAGATCTC
GATCCCGCGAAATTAATACGACTCACTATAGGGGAATTGTGAGCGGATAACAATTCCCCTCTAGAAATAA
TTTTGTTTAACTTTAAGAAGGAGATATACCATGGGAGACCTCCCCTATCAGTGATAGAGATTGACATCCC
TATCAGTGATAGAGATACTGAGCACGGATCTTAGCTACTAGAGAAAGAGGAGAAATACTAGATGCGTAAA
GGCGAAGAGCTGTTCACTGGTGTCGTCCCTATTCTGGTGGAACTGGATGGTGATGTCAACGGTCATAAGT
TTTCCGTGCGTGGCGAGGGTGAAGGTGACGCAACTAATGGTAAACTGACGCTGAAGTTCATCTGTACTAC
TGGTAAACTGCCGGTACCTTGGCCGACTCTGGTAACGACGCTGACTTATGGTGTTCAGTGCTTTGCTCGT
TATCCGGACCATATGAAGCAGCATGACTTCTTCAAGTCCGCCATGCCGGAAGGCTATGTGCAGGAACGCA
CGATTTCCTTTAAGGATGACGGCACGTACAAAACGCGTGCGGAAGTGAAATTTGAAGGCGATACCCTGGT
AAACCGCATTGAGCTGAAAGGCATTGACTTTAAAGAAGACGGCAATATCCTGGGCCATAAGCTGGAATAC
AATTTTAACAGCCACAATGTTTACATCACCGCCGATAAACAAAAAAATGGCATTAAAGCGAATTTTAAAA
TTCGCCACAACGTGGAGGATGGATCTGTGCAGCTGGCTGATCACTACCAGCAAAACACTCCAATCGGTGA
TGGTCCTGTTCTGCTGCCAGACAATCACTATCTGAGCACGCAAAGCGTTCTGTCTAAAGATCCGAACGAG
AAACGCGATCATATGGTTCTGCTGGAGTTCGTAACCGCAGCGGGCATCACGCATGGTATGGATGAACTGT
ACAAATGAGGTCTCTTAAGAGCTCCGTCGACAAGCTTGCGGCCGCACTCGAGCACCACCACCACCACCAC
TGAGATCCGGCTGCTAACAAAGCCCGAAAGGAAGCTGAGTTGGCTGCTGCCACCGCTGAGCAATAACTAG
CATAACCCCTTGGGGCCTCTAAACGGGTCTTGAGGGGTTTTTTGCTGAAAGGAGGAACTATATCCGGAT

**pET28a**

TGGCGAATGGGACGCGCCCTGTAGCGGCGCATTAAGCGCGGCGGGTGTGGTGGTTACGCGCAGCGTGACC
GCTACACTTGCCAGCGCCCTAGCGCCCGCTCCTTTCGCTTTCTTCCCTTCCTTTCTCGCCACGTTCGCCG
GCTTTCCCCGTCAAGCTCTAAATCGGGGGCTCCCTTTAGGGTTCCGATTTAGTGCTTTACGGCACCTCGA
CCCCAAAAAACTTGATTAGGGTGATGGTTCACGTAGTGGGCCATCGCCCTGATAGACGGTTTTTCGCCCT
TTGACGTTGGAGTCCACGTTCTTTAATAGTGGACTCTTGTTCCAAACTGGAACAACACTCAACCCTATCT
CGGTCTATTCTTTTGATTTATAAGGGATTTTGCCGATTTCGGCCTATTGGTTAAAAAATGAGCTGATTTA
ACAAAAATTTAACGCGAATTTTAACAAAATATTAACGCTTACAATTTAGGTGGCACTTTTCGGGGAAATG
TGCGCGGAACCCCTATTTGTTTATTTTTCTAAATACATTCAAATATGTATCCGCTCATGAATTAATTCTT
AGAAAAACTCATCGAGCATCAAATGAAACTGCAATTTATTCATATCAGGATTATCAATACCATATTTTTG
AAAAAGCCGTTTCTGTAATGAAGGAGAAAACTCACCGAGGCAGTTCCATAGGATGGCAAGATCCTGGTAT
CGGTCTGCGATTCCGACTCGTCCAACATCAATACAACCTATTAATTTCCCCTCGTCAAAAATAAGGTTAT
CAAGTGAGAAATCACCATGAGTGACGACTGAATCCGGTGAGAATGGCAAAAGTTTATGCATTTCTTTCCA
GACTTGTTCAACAGGCCAGCCATTACGCTCGTCATCAAAATCACTCGCATCAACCAAACCGTTATTCATT
CGTGATTGCGCCTGAGCGAGACGAAATACGCGATCGCTGTTAAAAGGACAATTACAAACAGGAATCGAAT
GCAACCGGCGCAGGAACACTGCCAGCGCATCAACAATATTTTCACCTGAATCAGGATATTCTTCTAATAC
CTGGAATGCTGTTTTCCCGGGGATCGCAGTGGTGAGTAACCATGCATCATCAGGAGTACGGATAAAATGC
TTGATGGTCGGAAGAGGCATAAATTCCGTCAGCCAGTTTAGTCTGACCATCTCATCTGTAACATCATTGG
CAACGCTACCTTTGCCATGTTTCAGAAACAACTCTGGCGCATCGGGCTTCCCATACAATCGATAGATTGT
CGCACCTGATTGCCCGACATTATCGCGAGCCCATTTATACCCATATAAATCAGCATCCATGTTGGAATTT
AATCGCGGCCTAGAGCAAGACGTTTCCCGTTGAATATGGCTCATAACACCCCTTGTATTACTGTTTATGT
AAGCAGACAGTTTTATTGTTCATGACCAAAATCCCTTAACGTGAGTTTTCGTTCCACTGAGCGTCAGACC

CCGTAGAAAAGATCAAAGGATCTTCTTGAGATCCTTTTTTTCTGCGCGTAATCTGCTGCTTGCAAACAAA
AAAACCACCGCTACCAGCGGTGGTTTGTTTGCCGGATCAAGAGCTACCAACTCTTTTTCCGAAGGTAACT
GGCTTCAGCAGAGCGCAGATACCAAATACTGTCCTTCTAGTGTAGCCGTAGTTAGGCCACCACTTCAAGA
ACTCTGTAGCACCGCCTACATACCTCGCTCTGCTAATCCTGTTACCAGTGGCTGCTGCCAGTGGCGATAA
GTCGTGTCTTACCGGGTTGGACTCAAGACGATAGTTACCGGATAAGGCGCAGCGGTCGGGCTGAACGGGG
GGTTCGTGCACACAGCCCAGCTTGGAGCGAACGACCTACACCGAACTGAGATACCTACAGCGTGAGCTAT
GAGAAAGCGCCACGCTTCCCGAAGGGAGAAAGGCGGACAGGTATCCGGTAAGCGGCAGGGTCGGAACAGG
AGAGCGCACGAGGGAGCTTCCAGGGGGAAACGCCTGGTATCTTTATAGTCCTGTCGGGTTTCGCCACCTC
TGACTTGAGCGTCGATTTTTGTGATGCTCGTCAGGGGGGCGGAGCCTATGGAAAAACGCCAGCAACGCGG
CCTTTTTACGGTTCCTGGCCTTTTGCTGGCCTTTTGCTCACATGTTCTTTCCTGCGTTATCCCCTGATTC
TGTGGATAACCGTATTACCGCCTTTGAGTGAGCTGATACCGCTCGCCGCAGCCGAACGACCGAGCGCAGC
GAGTCAGTGAGCGAGGAAGCGGAAGAGCGCCTGATGCGGTATTTTCTCCTTACGCATCTGTGCGGTATTT
CACACCGCATATATGGTGCACTCTCAGTACAATCTGCTCTGATGCCGCATAGTTAAGCCAGTATACACTC
CGCTATCGCTACGTGACTGGGTCATGGCTGCGCCCCGACACCCGCCAACACCCGCTGACGCGCCCTGACG
GGCTTGTCTGCTCCCGGCATCCGCTTACAGACAAGCTGTGACCGTCTCCGGGAGCTGCATGTGTCAGAGG
TTTTCACCGTCATCACCGAAACGCGCGAGGCAGCTGCGGTAAAGCTCATCAGCGTGGTCGTGAAGCGATT
CACAGATGTCTGCCTGTTCATCCGCGTCCAGCTCGTTGAGTTTCTCCAGAAGCGTTAATGTCTGGCTTCT
GATAAAGCGGGCCATGTTAAGGGCGGTTTTTTCCTGTTTGGTCACTGATGCCTCCGTGTAAGGGGGATTT
CTGTTCATGGGGGTAATGATACCGATGAAACGAGAGAGGATGCTCACGATACGGGTTACTGATGATGAAC
ATGCCCGGTTACTGGAACGTTGTGAGGGTAAACAACTGGCGGTATGGATGCGGCGGGACCAGAGAAAAAT
CACTCAGGGTCAATGCCAGCGCTTCGTTAATACAGATGTAGGTGTTCCACAGGGTAGCCAGCAGCATCCT
GCGATGCAGATCCGGAACATAATGGTGCAGGGCGCTGACTTCCGCGTTTCCAGACTTTACGAAACACGGA
AACCGAAGACCATTCATGTTGTTGCTCAGGTCGCAGACGTTTTGCAGCAGCAGTCGCTTCACGTTCGCTC
GCGTATCGGTGATTCATTCTGCTAACCAGTAAGGCAACCCCGCCAGCCTAGCCGGGTCCTCAACGACAGG
AGCACGATCATGCGCACCCGTGGGGCCGCCATGCCGGCGATAATGGCCTGCTTCTCGCCGAAACGTTTGG
TGGCGGGACCAGTGACGAAGGCTTGAGCGAGGGCGTGCAAGATTCCGAATACCGCAAGCGACAGGCCGAT
CATCGTCGCGCTCCAGCGAAAGCGGTCCTCGCCGAAAATGACCCAGAGCGCTGCCGGCACCTGTCCTACG
AGTTGCATGATAAAGAAGACAGTCATAAGTGCGGCGACGATAGTCATGCCCCGCGCCCACCGGAAGGAGC
TGACTGGGTTGAAGGCTCTCAAGGGCATCGGTCGAGATCCCGGTGCCTAATGAGTGAGCTAACTTACATT
AATTGCGTTGCGCTCACTGCCCGCTTTCCAGTCGGGAAACCTGTCGTGCCAGCTGCATTAATGAATCGGC
CAACGCGCGGGGAGAGGCGGTTTGCGTATTGGGCGCCAGGGTGGTTTTTCTTTTCACCAGTGAGACGGGC
AACAGCTGATTGCCCTTCACCGCCTGGCCCTGAGAGAGTTGCAGCAAGCGGTCCACGCTGGTTTGCCCCA
GCAGGCGAAAATCCTGTTTGATGGTGGTTAACGGCGGGATATAACATGAGCTGTCTTCGGTATCGTCGTA
TCCCACTACCGAGATATCCGCACCAACGCGCAGCCCGGACTCGGTAATGGCGCGCATTGCGCCCAGCGCC
ATCTGATCGTTGGCAACCAGCATCGCAGTGGGAACGATGCCCTCATTCAGCATTTGCATGGTTTGTTGAA
AACCGGACATGGCACTCCAGTCGCCTTCCCGTTCCGCTATCGGCTGAATTTGATTGCGAGTGAGATATTT
ATGCCAGCCAGCCAGACGCAGACGCGCCGAGACAGAACTTAATGGGCCCGCTAACAGCGCGATTTGCTGG
TGACCCAATGCGACCAGATGCTCCACGCCCAGTCGCGTACCGTCTTCATGGGAGAAAATAATACTGTTGA
TGGGTGTCTGGTCAGAGACATCAAGAAATAACGCCGGAACATTAGTGCAGGCAGCTTCCACAGCAATGGC
ATCCTGGTCATCCAGCGGATAGTTAATGATCAGCCCACTGACGCGTTGCGCGAGAAGATTGTGCACCGCC
GCTTTACAGGCTTCGACGCCGCTTCGTTCTACCATCGACACCACCACGCTGGCACCCAGTTGATCGGCGC
GAGATTTAATCGCCGCGACAATTTGCGACGGCGCGTGCAGGGCCAGACTGGAGGTGGCAACGCCAATCAG
CAACGACTGTTTGCCCGCCAGTTGTTGTGCCACGCGGTTGGGAATGTAATTCAGCTCCGCCATCGCCGCT
TCCACTTTTTCCCGCGTTTTCGCAGAAACGTGGCTGGCCTGGTTCACCACGCGGGAAACGGTCTGATAAG
AGACACCGGCATACTCTGCGACATCGTATAACGTTACTGGTTTCACATTCACCACCCTGAATTGACTCTC
TTCCGGGCGCTATCATGCCATACCGCGAAAGGTTTTGCGCCATTCGATGGTGTCCGGGATCTCGACGCTC
TCCCTTATGCGACTCCTGCATTAGGAAGCAGCCCAGTAGTAGGTTGAGGCCGTTGAGCACCGCCGCCGCA
AGGAATGGTGCATGCAAGGAGATGGCGCCCAACAGTCCCCCGGCCACGGGGCCTGCCACCATACCCACGC
CGAAACAAGCGCTCATGAGCCCGAAGTGGCGAGCCCGATCTTCCCCATCGGTGATGTCGGCGATATAGGC
GCCAGCAACCGCACCTGTGGCGCCGGTGATGCCGGCCACGATGCGTCCGGCGTAGAGGATCGAGATCTCG
ATCCCGCGAAATTAATACGACTCACTATAGGGGAATTGTGAGCGGATAACAATTCCCCTCTAGAAATAAT
TTTGTTTAACTTTAAGAAGGAGATATACCATGGGCAGCAGCCATCATCATCATCATCACAGCAGCGGCCT

GGTGCCGCGCGGCAGCCATATGGCTAGCATGACTGGTGGACAGCAAATGGGTCGCGGATCCGAATTCGAG
CTCCGTCGACAAGCTTGCGGCCGCACTCGAGCACCACCACCACCACCACTGAGATCCGGCTGCTAACAAA
GCCCGAAAGGAAGCTGAGTTGGCTGCTGCCACCGCTGAGCAATAACTAGCATAACCCCTTGGGGCCTCTA
AACGGGTCTTGAGGGGTTTTTTGCTGAAAGGAGGAACTATATCCGGAT

**pET47b(+)**

ATCCGGATATAGTTCCTCCTTTCAGCAAAAAACCCCTCAAGACCCGTTTAGAGGCCCCAAGGGGTTATGC
TAGTTATTGCTCAGCGGTGGCAGCAGCCAACTCAGCTTCCTTTCGGGCTTTGTTAGCAGCCTAGGTTAA
TTAAGCCTCGAGAGCAGCAGAAGTAGAGCTGTCCATGTGCTGGCGTTCGAATTTAGCAGCAGCGGTTTCT
TTACTACCGCGTGGCACCAGAGCGAGCTCTGCGGCCGCAAGCTTGTCGACGGACGTCGGGCGCGCCAAGG
CCTGTACAGAATTCGGATCCTGGTACCCGGGTCCCTGAAAGAGGACTTCAAGAGCCGCGGAGTGATGGTG
GTGGTGATGTGCCATATGTATATCTCCTTCTTAAAGTTAAACAAAATTATTTCTAGAGGGGAATTGTTAT
CCGCTCACAATTCCCCTATAGTGAGTCGTATTAATTTCGCGGGATCGAGATCGATCTCGATCCTCTACGC
CGGACGCATCGTGGCCGGCATCACCGGCGCCACAGGTGCGGTTGCTGGCGCCTATATCGCCGACATCACC
GATGGGGAAGATCGGGCTCGCCACTTCGGGCTCATGAGCGCTTGTTTCGGCGTGGGTATGGTGGCAGGCC
CCGTGGCCGGGGGACTGTTGGGCGCCATCTCCTTGCATGCACCATTCCTTGCGGCGGCGGTGCTCAACGG
CCTCAACCTACTACTGGGCTGCTTCCTAATGCAGGAGTCGCATAAGGGAGAGCGTCGAGATCCCGGACAC
CATCGAATGGCGCAAAACCTTTCGCGGTATGGCATGATAGCGCCCGGAAGAGAGTCAATTCAGGGTGGTG
AATGTGAAACCAGTAACGTTATACGATGTCGCAGAGTATGCCGGTGTCTCTTATCAGACCGTTTCCCGCG
TGGTGAACCAGGCCAGCCACGTTTCTGCGAAAACGCGGGAAAAAGTGGAAGCGGCGATGGCGGAGCTGAA
TTACATTCCCAACCGCGTGGCACAACAACTGGCGGGCAAACAGTCGTTGCTGATTGGCGTTGCCACCTCC
AGTCTGGCCCTGCACGCGCCGTCGCAAATTGTCGCGGCGATTAAATCTCGCGCCGATCAACTGGGTGCCA
GCGTGGTGGTGTCGATGGTAGAACGAAGCGGCGTCGAAGCCTGTAAAGCGGCGGTGCACAATCTTCTCGC
GCAACGCGTCAGTGGGCTGATCATTAACTATCCGCTGGATGACCAGGATGCCATTGCTGTGGAAGCTGCC
TGCACTAATGTTCCGGCGTTATTTCTTGATGTCTCTGACCAGACACCCATCAACAGTATTATTTTCTCCC
ATGAAGACGGTACGCGACTGGGCGTGGAGCATCTGGTCGCATTGGGTCACCAGCAAATCGCGCTGTTAGC
GGGCCCATTAAGTTCTGTCTCGGCGCGTCTGCGTCTGGCTGGCTGGCATAAATATCTCACTCGCAATCAA
ATTCAGCCGATAGCGGAACGGGAAGGCGACTGGAGTGCCATGTCCGGTTTTCAACAAACCATGCAAATGC
TGAATGAGGGCATCGTTCCCACTGCGATGCTGGTTGCCAACGATCAGATGGCGCTGGGCGCAATGCGCGC
CATTACCGAGTCCGGGCTGCGCGTTGGTGCGGACATCTCGGTAGTGGGATACGACGATACCGAAGACAGC
TCATGTTATATCCCGCCGTTAACCACCATCAAACAGGATTTTCGCCTGCTGGGGCAAACCAGCGTGGACC
GCTTGCTGCAACTCTCTCAGGGCCAGGCGGTGAAGGGCAATCAGCTGTTGCCCGTCTCACTGGTGAAAAG
AAAAACCACCCTGGCGCCCAATACGCAAACCGCCTCTCCCCGCGCGTTGGCCGATTCATTAATGCAGCTG
GCACGACAGGTTTCCCGACTGGAAAGCGGGCAGTGAGCGCAACGCAATTAATGTAAGTTAGCTCACTCAT
TAGGCACCGGGATCTCGACCGATGCCCTTGAGAGCCTTCAACCCAGTCAGCTCCTTCCGGTGGGCGCGGG
GCATGACTAGCATGATCGTGCTCCTGTCGTTGAGGACCCGGCTAGGCTGGCGGGGTTGCCTTACTGGTTA
GCAGAATGAATCACCGATACGCGAGCGAACGTGAAGCGACTGCTGCTGCAAAACGTCTGCGACCTGAGCA
ACAACATGAATGGTCTTCGGTTTCCGTGTTTCGTAAAGTCTGGAAACGCGGAAGTCAGCGCCCTGCACCA
TTATGTTCCGGATCTGCATCGCAGGATGCTGCTGGCTACCCTGTGGAACACCTACATCTGTATTAACGAA
GCGCTGGCATTGACCCTGAGTGATTTTTCTCTGGTCCCGCCGCATCCATACCGCCAGTTGTTTACCCTCA
CAACGTTCCAGTAACCGGGCATGTTCATCATCAGTAACCCGTATCGTGAGCATCCTCTCTCGTTTCATCG
GTATCATTACCCCCATGAACAGAAATCCCCCTTACACGGAGGCATCAGTGACCAAACAGGAAAAAACCGC
CCTTAACATGGCCCGCTTTATCAGAAGCCAGACATTAACGCTTCTGGAGAAACTCAACGAGCTGGACGCG
GATGAACAGGCAGACATCTGTGAATCGCTTCACGACCACGCTGATGAGCTTTACCGCAGCTGCCTCGCGC
GTTTCGGTGATGACGGTGAAAACCTCTGACACATGCAGCTCCCGGAGACGGTCACAGCTTGTCTGTAAGC
GGATGCCGGGAGCAGACAAGCCCGTCAGGGCGCGTCAGCGGGTGTTGGCGGGTGTCGGGGCGCAGCCATG
ACCCAGTCACGTAGCGATAGCGGAGTGTATACTGGCTTAACTATGCGGCATCAGAGCAGATTGTACTGAG
AGTGCACCATATATGCGGTGTGAAATACCGCACAGATGCGTAAGGAGAAAATACCGCATCAGGCGCTCTT
CCGCTTCCTCGCTCACTGACTCGCTGCGCTCGGTCGTTCGGCTGCGGCGAGCGGTATCAGCTCACTCAAA
GGCGGTAATACGGTTATCCACAGAATCAGGGGATAACGCAGGAAAGAACATGTGAGCAAAAGGCCAGCAA
AAGGCCAGGAACCGTAAAAAGGCCGCGTTGCTGGCGTTTTTCCATAGGCTCCGCCCCCCTGACGAGCATC

ACAAAAATCGACGCTCAAGTCAGAGGTGGCGAAACCCGACAGGACTATAAAGATACCAGGCGTTTCCCCC
TGGAAGCTCCCTCGTGCGCTCTCCTGTTCCGACCCTGCCGCTTACCGGATACCTGTCCGCCTTTCTCCCT
TCGGGAAGCGTGGCGCTTTCTCATAGCTCACGCTGTAGGTATCTCAGTTCGGTGTAGGTCGTTCGCTCCA
AGCTGGGCTGTGTGCACGAACCCCCCGTTCAGCCCGACCGCTGCGCCTTATCCGGTAACTATCGTCTTGA
GTCCAACCCGGTAAGACACGACTTATCGCCACTGGCAGCAGCCACTGGTAACAGGATTAGCAGAGCGAGG
TATGTAGGCGGTGCTACAGAGTTCTTGAAGTGGTGGCCTAACTACGGCTACACTAGAAGGACAGTATTTG
GTATCTGCGCTCTGCTGAAGCCAGTTACCTTCGGAAAAAGAGTTGGTAGCTCTTGATCCGGCAAACAAAC
CACCGCTGGTAGCGGTGGTTTTTTTGTTTGCAAGCAGCAGATTACGCGCAGAAAAAAAGGATCTCAAGAA
GATCCTTTGATCTTTTCTACGGGGTCTGACGCTCAGTGGAACGAAAACTCACGTTAAGGGATTTTGGTCA
TGAACAATAAAACTGTCTGCTTACATAAACAGTAATACAAGGGGTGTTATGAGCCATATTCAACGGGAAA
CGTCTTGCTCTAGGCCGCGATTAAATTCCAACATGGATGCTGATTTATATGGGTATAAATGGGCTCGCGA
TAATGTCGGGCAATCAGGTGCGACAATCTATCGATTGTATGGGAAGCCCGATGCGCCAGAGTTGTTTCTG
AAACATGGCAAAGGTAGCGTTGCCAATGATGTTACAGATGAGATGGTCAGACTAAACTGGCTGACGGAAT
TTATGCCTCTTCCGACCATCAAGCATTTTATCCGTACTCCTGATGATGCATGGTTACTCACCACTGCGAT
CCCCGGCAAAACAGCATTCCAGGTATTAGAAGAATATCCTGATTCAGGTGAAAATATTGTTGATGCGCTG
GCAGTGTTCCTGCGCCGGTTGCATTCGATTCCTGTTTGTAATTGTCCTTTTAACAGTGATCGCGTATTTC
GTCTCGCTCAGGCGCAATCACGAATGAATAACGGTTTGGTTGATGCGAGTGATTTTGATGACGAGCGTAA
TGGCTGGCCTGTTGAACAAGTCTGGAAAGAAATGCATAAACTTTTGCCATTCTCACCGGATTCAGTCGTC
ACTCATGGTGATTTCTCACTTGATAACCTTATTTTTGACGAGGGGAAATTAATAGGTTGTATTGATGTTG
GACGAGTCGGAATCGCAGACCGATACCAGGATCTTGCCATCCTATGGAACTGCCTCGGTGAGTTTTCTCC
TTCATTACAGAAACGGCTTTTTCAAAAATATGGTATTGATAATCCTGATATGAATAAATTGCAGTTTCAT
TTGATGCTCGATGAGTTTTTCTAAGAATTAATTCATGAGCGGATACATATTTGAATGTATTTAGAAAAAT
AAACAAATAGGGGTTCCGCGCACATTTCCCCGAAAAGTGCCACCTGAAATTGTAAACGTTAATATTTTGT
TAAAATTCGCGTTAAATTTTTGTTAAATCAGCTCATTTTTTAACCAATAGGCCGAAATCGGCAAAATCCC
TTATAAATCAAAAGAATAGACCGAGATAGGGTTGAGTGTTGTTCCAGTTTGGAACAAGAGTCCACTATTA
AAGAACGTGGACTCCAACGTCAAAGGGCGAAAAACCGTCTATCAGGGCGATGGCCCACTACGTGAACCAT
CACCCTAATCAAGTTTTTTGGGGTCGAGGTGCCGTAAAGCACTAAATCGGAACCCTAAAGGGAGCCCCCG
ATTTAGAGCTTGACGGGGAAAGCCGGCGAACGTGGCGAGAAAGGAAGGGAAGAAAGCGAAAGGAGCGGGC
GCTAGGGCGCTGGCAAGTGTAGCGGTCACGCTGCGCGTAACCACCACACCCGCCGCGCTTAATGCGCCGC
TACAGGGCGCGTCCCATTCGCCA

**TxTl T7 A1**

ATCCCCAGGCATCAAATAAAACGAAAGGCTCAGTCGAAAGACTGGGCCTTTCGTTTTATCTGTTGTTTGT
CGGTGAACGCTCTCTACTAGAGTCACACTGGCTCACCTTCGGGTGGGCCTTTCTGCGTTTATAGCTGAGC
ATGAGACGGAAATCTGCTCGTCAGTGGTGCTCACACTGACGAATCATGTACAGATCATACCGATGACTGC
CTGGCGACTCACAACTAAGCAAGACAGCCGGAACCAGCGCCGGCGAACACCACTGCATATATGGCATATC
ACAACAGTCCAACTAGTGCACTGCAGTACATTATTCTTAGAAAAACTCATCGAGCATCAAATGAAACTGC
AATTTATTCATATCAGGATTATCAATACCATATTTTTGAAAAAGCCGTTTCTGTAATGAAGGAGAAAACT
CACCGAGGCAGTTCCATAGGATGGCAAGATCCTGGTATCGGTCTGCGATTCCGACTCGTCCAACATCAAT
ACAACCTATTAATTTCCCCTCGTCAAAAATAAGGTTATCAAGTGAGAAATCACCATGAGTGACGACTGAA
TCCGGTGAGAATGGCAAAAGTTTATGCATTTCTTTCCAGACTTGTTCAACAGGCCAGCCATTACGCTCGT
CATCAAAATCACTCGCATCAACCAAACCGTTATTCATTCGTGATTGCGCCTGAGCGAGACGAAATACGCG
ATCGCTGTTAAAAGGACAATTACAAACAGGAATCGAATGCAACCGGCGCAGGAACACTGCCAGCGCATCA
ACAATATTTTCACCTGAATCAGGATATTCTTCTAATACCTGGAATGCTGTTTTCCCGGGGATCGCAGTGG
TGAGTAACCATGCATCATCAGGAGTACGGATAAAATGCTTGATGGTCGGAAGAGGCATAAATTCCGTCAG
CCAGTTTAGTCTGACCATCTCATCTGTAACATCATTGGCAACGCTACCTTTGCCATGTTTCAGAAACAAC
TCTGGCGCATCGGGCTTCCCATACAATCGATAGATTGTCGCACCTGATTGCCCGACATTATCGCGAGCCC
ATTTATACCCATATAAATCAGCATCCATGTTGGAATTTAATCGCGGCCTAGAGCAAGACGTTTCCCGTTG
AATATGGCTCATAACACCCCTTGTATTACTGTTTATGTAAGCAGACAGTTTTATTGGAGTTTTTTCCATA
GGCTCCGCCCCCCTGACAAGCATCACGAAATCTGACGCTCAAATCAGTGGTGGCGAAACCCGACAGGACT
ATAAAGATACCAGGCGTTTCCCCCTGGCGGCTCCCTCGTGCGCTCTCCTGTTCCTGCCTTTCGGTTTACC

GGTGTCATTCCGCTGTTATGGCCGCGTTTGTCTCATTCCACGCCTGACACTCAGTTCCGGGTAGGCAGTT
CGCTCCAAGCTGGACTGTATGCACGAACCCCCCGTTCAGTCCGACCGCTGCGCCTTATCCGGTAACTATC
GTCTTGAGTCCAACCCGGAAAGACATGCAAAAGCACCACTGGCAGCAGCCACTGGTAATTGATTTAGAGG
AGTTAGTCTTGAAGTCATGCGCCGGTTAAGGCTAAACTGAAAGGACAAGTTTTGGTGACTGCGCTCCTCC
AAGCCAGTTACCTCGGTTCAAAGAGTTGGTAGCTCAGAGAACCTTCGAAAAACCGCCCTGCAAGGCGGTT
TTTTCGTTTTCAGAGCAAGAGATTACGCGCAGACCAAAACGATCTCAAGAAGATCATCTTATTCCCTGAA
TTCGCATCTAGACTGATGAGACGTGGTAGAGCCACAAACAGCCGGTACAAGCAACGATCTCCAGGACCAT
CTGAATCATGCGCGGATGACACGAACTCACGACGGCGATCACAGACATTAACCCACAGTACAGACACTGC
GACAACGTGGCAATTCGTCGCAATACAACGGAGGCCGTTGAGCACCGCCGCCGCAAGGAATGGTGCATGC
AAGGAGATGGCGCCCAACAGTCCCCCGGCCACGGGGCCTGCCACCATACCCACGCCGAAACAAGCGCTCA
TGAGCCCGAAGTGGCGAGCCCGATCTTCCCCATCGGTGATGTCGGCGATATAGGCGCCAGCAACCGCACC
TGTGGCGCCGGTGATGCCGGCCACGATGCGTCCGGCGTAGAGGATCGAGATCTCGATCCCGCGAAATTAA
TACGACTCACTATAGGGGAATTGTGAGCGGATAACAATTCCCCTCTAGAAATAATTTTGTTTAACTTTAA
GAAGGAGATATAT

**TxTl T7 A2**

ATCCCCAGGCATCAAATAAAACGAAAGGCTCAGTCGAAAGACTGGGCCTTTCGTTTTATCTGTTGTTTGT
CGGTGAACGCTCTCTACTAGAGTCACACTGGCTCACCTTCGGGTGGGCCTTTCTGCGTTTATAGCTGCCA
ATGAGACGACGGGGTCATCACGGCTCATCATGCGCCCAACAAATGTGTGCCATACACGCTCGGATGACTG
CCTGATGACCGCACTGACTGGGGACAGCCGATCCACCTAAGCCTGTGAGAGAAGCAGACACCCGACAGAT
CAAGGCAGTTAACTAGTGCACTGCAGTACATTATTCTTAGAAAAACTCATCGAGCATCAAATGAAACTGC
AATTTATTCATATCAGGATTATCAATACCATATTTTTGAAAAAGCCGTTTCTGTAATGAAGGAGAAAACT
CACCGAGGCAGTTCCATAGGATGGCAAGATCCTGGTATCGGTCTGCGATTCCGACTCGTCCAACATCAAT
ACAACCTATTAATTTCCCCTCGTCAAAAATAAGGTTATCAAGTGAGAAATCACCATGAGTGACGACTGAA
TCCGGTGAGAATGGCAAAAGTTTATGCATTTCTTTCCAGACTTGTTCAACAGGCCAGCCATTACGCTCGT
CATCAAAATCACTCGCATCAACCAAACCGTTATTCATTCGTGATTGCGCCTGAGCGAGACGAAATACGCG
ATCGCTGTTAAAAGGACAATTACAAACAGGAATCGAATGCAACCGGCGCAGGAACACTGCCAGCGCATCA
ACAATATTTTCACCTGAATCAGGATATTCTTCTAATACCTGGAATGCTGTTTTCCCGGGGATCGCAGTGG
TGAGTAACCATGCATCATCAGGAGTACGGATAAAATGCTTGATGGTCGGAAGAGGCATAAATTCCGTCAG
CCAGTTTAGTCTGACCATCTCATCTGTAACATCATTGGCAACGCTACCTTTGCCATGTTTCAGAAACAAC
TCTGGCGCATCGGGCTTCCCATACAATCGATAGATTGTCGCACCTGATTGCCCGACATTATCGCGAGCCC
ATTTATACCCATATAAATCAGCATCCATGTTGGAATTTAATCGCGGCCTAGAGCAAGACGTTTCCCGTTG
AATATGGCTCATAACACCCCTTGTATTACTGTTTATGTAAGCAGACAGTTTTATTGGAGTTTTTTCCATA
GGCTCCGCCCCCCTGACAAGCATCACGAAATCTGACGCTCAAATCAGTGGTGGCGAAACCCGACAGGACT
ATAAAGATACCAGGCGTTTCCCCCTGGCGGCTCCCTCGTGCGCTCTCCTGTTCCTGCCTTTCGGTTTACC
GGTGTCATTCCGCTGTTATGGCCGCGTTTGTCTCATTCCACGCCTGACACTCAGTTCCGGGTAGGCAGTT
CGCTCCAAGCTGGACTGTATGCACGAACCCCCCGTTCAGTCCGACCGCTGCGCCTTATCCGGTAACTATC
GTCTTGAGTCCAACCCGGAAAGACATGCAAAAGCACCACTGGCAGCAGCCACTGGTAATTGATTTAGAGG
AGTTAGTCTTGAAGTCATGCGCCGGTTAAGGCTAAACTGAAAGGACAAGTTTTGGTGACTGCGCTCCTCC
AAGCCAGTTACCTCGGTTCAAAGAGTTGGTAGCTCAGAGAACCTTCGAAAAACCGCCCTGCAAGGCGGTT
TTTTCGTTTTCAGAGCAAGAGATTACGCGCAGACCAAAACGATCTCAAGAAGATCATCTTATTCCCTGAA
TTCGCATCTAGACTGATGAGACGTGGTAGAGCCACAAACAGCCGGTACAAGCAACGATCTCCAGGACCAT
CTGAATCATGCGCGGATGACACGAACTCACGACGGCGATCACAGACATTAACCCACAGTACAGACACTGC
GACAACGTGGCAATTCGTCGCAATACAACGGAGGCCGTTGAGCACCGCCGCCGCAAGGAATGGTGCATGC
AAGGAGATGGCGCCCAACAGTCCCCCGGCCACGGGGCCTGCCACCATACCCACGCCGAAACAAGCGCTCA
TGAGCCCGAAGTGGCGAGCCCGATCTTCCCCATCGGTGATGTCGGCGATATAGGCGCCAGCAACCGCACC
TGTGGCGCCGGTGATGCCGGCCACGATGCGTCCGGCGTAGAGGATCGAGATCTCGATCCCGCGAAATTAA
TACGACTCACTATAGGGGAATTGTGAGCGGATAACAATTCCCCTCTAGAAATAATTTTGTTTAACTTTAA
GAAGGAGATATAT

**TxTl T500 A3**

CTCGAGGAATTCGACTCAATTAGTTCAGTCAGTTTCAGGATATTAGTCATCTCTACATTGATTATGAGTA
TTCAGAAATTCCTTAAATATTCTGACAAATGCTCTTTCCCTAAACTCCCCCCATAAAAAAACCCGCCGAA
GCGGGTTTTTACGTTATTTGCGGATTAACGATTACTCGTTATCAGAACCGCCCAGACCTGCGTTCAGCAG
TTCTGCCAGGCTGGCAGATGCGTCTTCCGAATTGATCCGTCGACCAAAGCCCGCCGAAAGGCGGGCTTTT
CTGTGCCGGCATGATAAGCTGTCAAACATGAGAATTACAACTTATATCGTATGGGGCTGACTTCAGGTGC
TACATTTGAAGAGATAAATTGCACTGAAATCTAGAAATATTTTATCTGATTAATAAGATGATCTTCTTGA
GATCGTTTTGGTCTGCGCGTAATCTCTTGCTCTGAAAACGAAAAAACCGCCTTGCAGGGCGGTTTTTCGA
AGGTTCTCTGAGCTACCAACTCTTTGAACCGAGGTAACTGGCTTGGAGGAGCGCAGTCACCAAAACTTGT
CCTTTCAGTTTAGCCTTAACCGGCGCATGACTTCAAGACTAACTCCTCTAAATCAATTACCAGTGGCTGC
TGCCAGTGGTGCTTTTGCATGTCTTTCCGGGTTGGACTCAAGACGATAGTTACCGGATAAGGCGCAGCGG
TCGGACTGAACGGGGGGTTCGTGCATACAGTCCAGCTTGGAGCGAACTGCCTACCCGGAACTGAGTGTCA
GGCGTGGAATGAGACAAACGCGGCCATAACAGCGGAATGACACCGGTAAACCGAAAGGCAGGAACAGGAG
AGCGCACGAGGGAGCCGCCAGGGGAAACGCCTGGTATCTTTATAGTCCTGTCGGGTTTCGCCACCACTGA
TTTGAGCGTCAGATTTCGTGATGCTTGTCAGGGGGGCGGAGCCTATGGAAAAACGGCTTTGCCGCGGCCC
TCTCACTTCCCTGTTAAGTATCTTCCTGGCATCTTCCAGGAAATCTCCGCCCCGTTCGTAAGCCATTTCC
GCTCGCCGCAGTCGAACGACCGAGCGTAGCGAGTCAGTGAGCGAGGAAGCGGAATATATCCTGTATCACA
TATTCTGCTGACGCACCGGTGCAGCCTTTTTTCTCCTGCCACATGAAGCACTTCACTGACACCCTCATCA
GTGCCAACATAGTAAGCCAGTATACACTCCGCTAGGGTCATGAGATTATCAAAAAGGATCTTCACCTAGA
TCCTTTTAAATTAAAAATGAAGTTTTAAATCAATCTAAAGTATATATGAGTAAACTTGGTCTGACAGTTA
CCAATGCTTAATCAGTGAGGCACCTATCTCAGCGATCTGTCTATTTCGTTCATCCATAGTTGCCTGACTC
CCCGTCGTGTAGATAACTACGATACGGGAGGGCTTACCATCTGGCCCCAGTGCTGCAATGATACCGCGAG
ACCCACGCTCACCGGCTCCAGATTTATCAGCAATAAACCAGCCAGCCGGAAGGGCCGAGCGCAGAAGTGG
TCCTGCAACTTTATCCGCCTCCATCCAGTCTATTAATTGTTGCCGGGAAGCTAGAGTAAGTAGTTCGCCA
GTTAATAGTTTGCGCAACGTTGTTGCCATTGCTACAGGCATCGTGGTGTCACGCTCGTCGTTTGGTATGG
CTTCATTCAGCTCCGGTTCCCAACGATCAAGGCGAGTTACATGATCCCCCATGTTGTGCAAAAAAGCGGT
TAGCTCCTTCGGTCCTCCGATCGTTGTCAGAAGTAAGTTGGCCGCAGTGTTATCACTCATGGTTATGGCA
GCACTGCATAATTCTCTTACTGTCATGCCATCCGTAAGATGCTTTTCTGTGACTGGTGAGTACTCAACCA
AGTCATTCTGAGAATAGTGTATGCGGCGACCGAGTTGCTCTTGCCCGGCGTCAATACGGGATAATACCGC
GCCACATAGCAGAACTTTAAAAGTGCTCATCATTGGAAAACGTTCTTCGGGGCGAAAACTCTCAAGGATC
TTACCGCTGTTGAGATCCAGTTCGATGTAACCCACTCGTGCACCCAACTGATCTTCAGCATCTTTTACTT
TCACCAGCGTTTCTGGGTGAGCAAAAACAGGAAGGCAAAATGCCGCAAAAAAGGGAATAAGGGCGACACG
GAAATGTTGAATACTCATACTCTTCCTTTTTCAATATTATTGAAGCATTTATCAGGGTTATTGTCTCATG
AGCGGATACATATTTGAATGTATTTAGAAAAATAAACAAATAGGGGTTCCGCGCACATTTCCCCGAAAAG
TGCCACCTGACGTCTAAGAAACCATTATTATCATGACATTAACCTATAAAAATAGGCGTATCACGAGGCC
CTTTCGTCTTCAAGAATTCTGGCGAATCCTCTGACCAGCCAGAAAACGACCTTTCTGTGGTGAAACCGGA
TGCTGCAATTCAGAGCGGCAGCAAGTGGGGGACAGCAGAAGACCTGACCGCCGCAGAGTGGATGTTTGAC
ATGGTGAAGACTATCGCACCATCAGCCAGAAAACCGAATTTTGCTGGGTGGGCTAACGATATCCGCCTGA
TGCGTGAACGTGACGGACGTAACCACCGCGACATGTGTGTGCTGTTCCGCTGGGCATGCCAGGACAACTT
CTGGTCCGGTAACGTGCTGAGCTAACACCGTGCGTGTTGACAATTTTACCTCTGGCGGTGATAATGGTTG
CAGCTAGCAATAATTTTGTTTAACTTTAAGAAGGAGGATCCAA

**FKBP12-1.1**

ATGGGCGGAGTGCAGGTGGAAACCATCTCCCCAGGAGACGGGCGCACCTTCCCCAAGCGCGGCCAGACCT
GCGTGGTGCACATTACCGGGATGCTTGAAGATGGAAAGAAATTTATTTCCTCCCGGGACAGAAACAAGCC
CTTTAAGTTTATGCTAGGCAAGCAGCGTGTGCTGCGAGGCTGGGAAGAAGGGGTTGCCCAGATGAGTGTG
GGTCAGAGAGCCAAACTGACTATATCTCCAGATTATGCCTTTGGTGCCACTGGGCACCCAGGCATCATCC
CACCACATGCCACTCTCGTCTTCGATGTGGAGCTTCTAAAACTGGAA

**FRB-1.1**

ATGGGTGTGGCCATCCTCTGGCATGAGATGTGGCATGAAGGCGCGGAAGAGGCAGCGCGTTTGTACCGTG
GGGAAAGGAACGTGAAAGGCATGTTTGAGGTGCTGGAGCCCTTGCATGCTATGATGGAACGGGGCCCCCA
GACTCTGAAGGAAACATCCTTTAATCAGGCCTATGGTCGAGATTTAATGGAGGCCCAAGAGTGGTGCAGG
AAGTACATGAAATCAGGGAATGTCAAGGACCTCTGGCAAGCCATGCTGCTCTATGCGCATGTGCGTGATC
GAATCTCA

**RapF-1.1**

ATGGGCAGCAGCAGCAGCATTGGCGAAAAGATTAACGAATGGTATATGTACATACGCCGATTCAGCATAC
CCGATGCAGCGTATTTGGCGTTTGAAATCGCGCAAGAGCTGGATCAAATGGAAGAAGATCAAGACCTTCA
TTTGTACTATTCACTGATGCTGTTTCGGGCGTATCTAATGGCCGAGTACCTTGAACCGTTAGAAAAAATG
AGGATTGAGGAACAGCCGAGACTGTCTGATCTGCTGCTTGAGATTGATAAAAAA

**RapF-1.2**

ATGGGCAGCAGCAGCAGCATTGGCGAAAAGATTAACGAATGGTATATGTACATACGCCGATTCAGCGTGC
CCGATGCAGCGTATTTGGGCTTTGAAATCAGCCAAGAGCTGGATCAAATGGAAGAAGATCAAGACCTTCA
TTTGTACTATTCACTGATGCTGTTTCGGGCGTATCTAATGCGTGAGTACCTTGAACCGTTAGAAAAAATG
AGGATTGAGGAACAGCCGAGACTGTCTGATCTGCTGCTTGAGATTGATAAAAAA

**RapF-1.3**

ATGGGCAGCAGCAGCAGCATTGGCGAAAAGATTAACGAATGGTATATGTACATACGCCGATTCAGCGTGC
CCGCGGCAGCGTATTTGGGCTTTGAAATCAGCCAAGAGCTGGATCAAATGGAAGAAGATCAAGACCTTCA
TTTGTACTATTCACTGATGCTGTTTCGGGCGTATCTAATGCGTGAGTACCTTGAACCGTTAGAAAAAATG
AGGATTGAGGAACAGCCGAGACTGTCTGATCTGCTGCTTGAGATTGATAAAAAA

**RapF-1.4**

ATGGGCAGCAGCAGCAGCATTGGCGAAAAGATTAACGAATTTTATATGTACATACGCCGATTCAGCATAC
CCGATGCAGCGTATTTGGCGTTTGAAATCGCGCAAGAGCTGGATCAAATGGAAGAAGATCAAGACCTTCA
TTTGTACTATTCACTGATGCTGTTTCGGGCGTATCTAATGGCCGAGTACCTTGAACCGTTAGAAAAAATG
AGGATTGAGGAACAGCCGAGACTGTCTGATCTGCTGCTTGAGATTGATAAAAAA

**ComA-1.1**

ATGGGTTCCTCTCAAAAAGAACAAGATGTGCTCACACCTAGAGAATGCCTGATTCTTCAAGAAGTTGAAA
AGGGATTTACAAACCAAGAAATCGCAGATGCCCTTCATTTACGTAAGAGCGCGATTGAAGCGAGCTTGAC
ATCGATTTTCAATAAGCTGAATGTCGGTTCACGGACGGAAGCGGTTTTGATTGCGAAATCAGACGGTGTA
CTT

**ComA-1.2**

ATGGGTTCCTCTCAAAAAGAACAAGATGTGCTCACACCTAGAGAATGCCTGATTCTTCAAGAAGTTGAAA
AGGGATTTACAAACCAAGAAATCGCAGATGCCCTTCACATGCGTAAGAGCGCGATTGAAGCGAGCTTGAC
ATCGATTTTCAATAAGCTGAATGTCGGTTCACGGACGGAAGCGGTTTTGATTGCGAAATCAGACGGTGTA
CTT

**ComA-1.3**

ATGGGTTCCTCTCAAAAAGAACAAGATGTGCTCACACCTAGAGAATGCCTGATTCTTCAAGAAGTTGAAA
AGGGATTTACAAACCAAGAAATCGCAGATGCCCTTCACATGCGTAAGAGCGCGATTGAAGCGAGCTTGAC
ATCGATTTTCGCGAAGCTGAATGTCGGTTCACGGACGGAAGCGGTTTTGATTGCGAAATCAGACGGTGTA
CTT

**ComA-1.4**

ATGGGTTCCTCTCAAAAAGAACAAGATGTGCTCACACCTAGAGAATGCCTGATTCTTCAAGAAGTTGAAA
AGGGATTTACAAACCAAGAAATCGCAGATGCCCTTCATTTACGTAAGAGCGCGATTGAAATGAGCTTGAC
ATCGATTTTCAATAAGCTGAATGTCGGTTCACGGACGGAAGCGGTTTTGATTGCGAAATCAGACGGTGTA
CTT

**AR-1.1**

SDLGRKLLEAARAGQDDEVRILMANGADVNAADNTGTTPLHLAAYSGHLEIVEVLLKHGADVDASDVFGF
TPLILAALWGHLEIVEVLLKNGADVNAMGSDGWTPLHAAAYFGYLEIVEVLLKHGADVNAQDKRGKTAFD
ISIDNGNEDLAEILQKLN

**AR-1.2**

SDLGRKLLEAARAGQDDEVRILMANGADVNAADNTGTTPLHLAAYSGHLEIVEVLLKHGADVDASDVFGL
TPLILAALWGHLEIVEVLLKNGADVNAMDSDGWTPLHAAAYFGYLEIVEVLLKHGADVNAQDKRGKTAFD
VSIDNGNEDLAEILQKLN

**AR-1.3**

SDLGRKLLEAARAGQDDEVRILMANGADVNAADNTGTTPLHLAAYSGHLEIVEVLLKHGADVDASDVFGF
TPLLLAALWGHLEIVEVLLKNGADVNAMGSDGWTPLHAAAWFGYLEIVEVLLKHGADVNAQDKRGKTAFD
ESIDNGNEDLAEILQKLN

**AR-1.4**

SDLGRKLLEAARAGQDDEVRILMANGADVNAADNTGTTPLHLAAYSGHLEIVEVLLKHGADVDASDVFGM
TPLFLAALWGHLEIVEVLLKNGADVNAMTSDGWTPLHAAAYFGYLEIVEVLLKHGADVNAQDKRGKTAFD
VSIDNGNEDLAEILQKLN

**AR-2.5**

SDLGRKLLEAARAGQDDEVRILMANGADVNAADNTGTTPLHLAAYSGHLEIVEVLLKHGADVDASDVFGF
TPLLLAALWGHLEIVEVLLKHGEDVNAMGSDGWTPLHAAAWFGYLEIVEVLLKHGADVNAQDKRGKTAFD
ESIDNGNEDLAEILQKLN

**AR-2.6**

SDLGRKLLEAARAGQDDEVRILMANGADVNAADNTGTTPLHLAAYSGHLEIVEVLLKHGADVDASDVFGF
TPLGLAALWGHLEIVEVLLKHGEDVNAMGSDGWTPLHAAAWFGYLEIVEVLLKHGADVNAQDKRGKTAFD
ESIDNGNEDLAEILQKLN

**AR-2.7**

SDLGRKLLEAARAGQDDEVRILMANGADVNAADNTGTTPLHLAAYSGHLEIVEVLLKHGADVDASDVFGF
TPLGLAALWGHLEIVEVLLKHGEDVNAMGSDGWTPLHAAAKFGYLEIVEVLLKHGADVNAQDKRGKTPFD
LAIDNGNEDIAEVLQKAA

**AR-3.8**

SDLGRKLLEAARAGQDDEVRILMANGADVNAADNTGTTPLHLAAYSGHLEIVEVLLKHGADVDASDVFGF
TPLGLAALWGHLEIVEVLLKHGEDVNAMGSDGWTPLHAAAKFGYLEIVEVLLKHGADVNAQDKKGLTPFD
LAILNGNEDIAEVLQKAA

**MBP-1.1**

KIEEGKLVIWINGDKGYNGLAEVGKKFEKDTGIKVTVEHPDKLEEKFPQVAATGDGPDIIFWAHDRFGGY
AQSGLLAEITPDKAFQDKLYPFTWDAVRYNGKLIAYPIAVEALSLIYNKDLLPNPPKTWEEIFALDKELK
AKGKSALMFNLQEPYFTWPLIAADGGYAFKYENGKYDIKDVGVDNAGAKAGLTFLVALIAAKAMNADTDY
SIAEAAFNKGETAMTINGPWAWSNIDTSKVNYGVTVLPTFKGQPSKPFVGVLSAGINAASPNKELAKEFL
ENYLLTDEGLEAVNKDKPLGAVALKSYEEELAKDPRIAATMENAQKGEIMPNIPQMSAFWYAVRTAVINA
ASG

**MBP-1.2**

KIEEGKLVIWINGDKGYNGLAEVGKKFEKDTGIKVTVEHPDKLEEKFPQVAATGDGPDIIFWAHDRFGGY
AQSGLLAEITPDKAFQDKLYPFTWDAVRYNGKLIAYPIAVEALSLIYNKDLLPNPPKTWEEIFALDKELK
AKGKSALMFNLQEPYFTWPLIAADGGYAFKYENGKYDIKDVGVDNAGAKAGLTFLVALIKAKHMNADTDY
SIAEAAFNKGETAMTINGPWAWSNIDTSKVNYGVTVLPTFKGQPSKPFVGVLSAGINAASPNKELAKEFL
ENYLLTDEGLEAVNKDKPLGAVALKSYEEELAKDPRIAATMENAQKGEIMPNIPQMSAFWYAVRTAVINA
ASG

**MBP-1.3**

KIEEGKLVIWINGDKGYNGLAEVGKKFEKDTGIKVTVEHPDKLEEKFPQVAATGDGPDIIFWAHDRFGGY
AQSGLLAEITPDKAFQDKLYPFTWDAVRYNGKLIAYPIAVEALSLIYNKDLLPNPPKTWEEIFALDKELK
AKGKSALMFNLQEPYFTWPLIAADGGYAFKYENGKYDIKDVGVDNAGAKAGLTRLVYLIAAKAMNADTDY
SIAEAAFNKGETAMTINGPWAWSNIDTSKVNYGVTVLPTFKGQPSKPFVGVLSAGINAASPNKELAKEFL
ENYLLTDEGLEAVNKDKPLGAVALKSYEEELAKDPRIAATMENAQKGEIMPNIPQMSAFWYAVRTAVINA
ASG

**MBP-1.4**

KIEEGKLVIWINGDKGYNGLAEVGKKFEKDTGIKVTVEHPDKLEEKFPQVAATGDGPDIIFWAHDRFGGY
AQSGLLAEITPDKAFQDKLYPFTWDAVRYNGKLIAYPIAVEALSLIYNKDLLPNPPKTWEEIFALDKELK
AKGKSALMFNLQEPYFTWPLIAADGGYAFKYENGKYDIKDVGVDNAGAKAGLTFLVALIKAKAMNADTDY
SIAEAAFNKGETAMTINGPWAWSNIDTSKVNYGVTVLPTFKGQPSKPFVGVLSAGINAASPNKELAKEFL
ENYLLTDEGLEAVNKDKPLGAVALKSYEEELAKDPRIAATMENAQKGEIMPNIPQMSAFWYAVRTAVINA
ASG

**MBP-2.5**

KIEEGKLVIWINGDKGYNGLAEVGKKFEKDTGIKVTVEHPDKLEEKFPQVAATGDGPDIIFWAHDRFGGY
AQSGLLAEITPDKAFQDKLYPFTWDAVRYNGKLIAYPIAVEALSLIYNKDLLPNPPKTWEEIFALDKELK
AKGKSALMFNLQEPYFTWPLIAADGGYAFKYENGKYDIKDVGVDNAGAKAGLTALVYLIAAKAMNADTDY
SIAEAAFNKGETAMTINGPWAWSNIDTSKVNYGVTVLPTFKGQPSKPFVGVLSAGINAASPNKELAKEFL
ENYLLTDEGLEAVNKDKPLGAVALKSYEEELAKDPRIAATMENAQKGEIMPNIPQMSAFWYAVRTAVINA
ASGRQTVDEALKDAQTRITK

**MBP-3.6**

KIEEGKLVIWINGDKGYNGLAEVGKKFEKDTGIKVTVEHPDKLEEKFPQVAATGDGPDIIFWAHDRFGGY
AQSGLLAEITPDKAFQDKLYPFTWDAVRYNGKLIAYPIAVEALSLIYNKDLLPNPPKTWEEIFALDKELK
AKGKSALMFNLQEPYFTWPLIAADGGYAFKYENGKYDIKDVGVDNAGAKAGLTALVALIAAKAMNADTDY
SIAEAAFNKGETAMTINGPWAWSNIDTSKVNYGVTVLPTFKGQPSKPFVGVLSAGINAASPNKELAKEFL
ENYLLTDEGLEAVNKDKPLGAVALKSYEEELAKDPRIAATMENAQKGEIMPNIPQMSAFWYAVRTAVINA
ASGRQTVDEALKDAQTRITK

**ERG12**

ATGTCATTACCGTTCTTAACTTCTGCACCGGGAAAGGTTATTATTTTTGGTGAACACTCTGCTGTGTACA
ACAAGCCTGCCGTCGCTGCTAGTGTGTCTGCGTTGAGAACCTACCTGCTAATAAGCGAGTCATCTGCACC
AGATACTATTGAATTGGACTTCCCGGACATTAGCTTTAATCATAAGTGGTCCATCAATGATTTCAATGCC
ATCACCGAGGATCAAGTAAACTCCCAAAAATTGGCCAAGGCTCAACAAGCCACCGATGGCTTGTCTCAGG
AACTCGTTAGTCTTTTGGACCCGTTGTTAGCTCAACTATCCGAATCCGCCCACTACCATGCAGCGTTTTG
TTTCCTGTATATGTTTGTTTGCCTATGCCCCCATGCCAAGAATATTAAGTTTTCTTTAAAGTCTACTTTA
CCCATCGGTGCTGGGTTGGGCTCAAGCGCCTCTATTTCTGTATCACTGGCCTTAGCTATGGCCTACTTGG
GGGGGTTAATAGGATCTAATGACTTGGAAAAGCTGTCAGAAAACGATAAGCATATAGTGAATCAATGGGC
CTTCATAGGTGAAAAGTGTGCTCACGGTACCCCTTCAGGAATAGATAACGCTGTGGCCACTTATGGTAAT
GCCCTGCTATTTGAAAAAGACTCACATAATGGAACAATAAACACAAACAATTTTAAGTTCTTAGATGATT
TCCCAGCCATTCCAATGATCCTAACCTATACTAGAATCCCAAGGTCTACAAAAGACCTTGTTGCTCGCGT
TCGTGTGTTGGTCACCGAGAAATTTCCTGAAGTTATGAAGCCAATTCTAGATGCCATGGGTGAATGTGCC
CTACAAGGCTTAGAGATCATGACTAAGTTAAGTAAATGTAAAGGCACCGATGACGAGGCTGTAGAAACTA
ATAATGAACTGTATGAACAACTATTGGAATTGATAAGAATAAATCATGGACTGCTTGTCTCAATCGGTGT
TTCTCATCCTGGATTAGAACTTATTAAAAATCTGAGCGATGATTTGAGAATTGGCTCCACAAAACTTACC
GGTGCTGGTGGCGGCGGTTGCTCTTTGACTTTGTTACGAAGAGACATTACTCAAGAGCAAATTGACAGCT
TCAAAAAGAAATTGCAAGATGATTTTAGTTACGAGACATTTGAAACAGACTTGGGTGGGACTGGCTGCTG
TTTGTTAAGCGCAAAAAATTTGAATAAAGACCTTAAAATCAAATCCCTAGTATTCCAATTATTTGAAAAT
AAAACTACCACAAAGCAACAAATTGACGATCTATTATTGCCAGGAAACACGAATTTACCATGGACTTCA

**ERG8**

ATGTCAGAGTTGAGAGCCTTCAGTGCCCCAGGGAAAGCGTTACTAGCTGGTGGATATTTAGTTTTAGATA
CAAAATATGAAGCATTTGTAGTCGGATTATCGGCAAGAATGCATGCTGTAGCCCATCCTTACGGTTCATT
GCAAGGGTCTGATAAGTTTGAAGTGCGTGTGAAAAGTAAACAATTTAAAGATGGGGAGTGGCTGTACCAT
ATAAGTCCTAAAAGTGGCTTCATTCCTGTTTCGATAGGCGGATCTAAGAACCCTTTCATTGAAAAAGTTA
TCGCTAACGTATTTAGCTACTTTAAACCTAACATGGACGACTACTGCAATAGAAACTTGTTCGTTATTGA
TATTTTCTCTGATGATGCCTACCATTCTCAGGAGGATAGCGTTACCGAACATCGTGGCAACAGAAGATTG
AGTTTTCATTCGCACAGAATTGAAGAAGTTCCCAAAACAGGGCTGGGCTCCTCGGCAGGTTTAGTCACAG
TTTTAACTACAGCTTTGGCCTCCTTTTTTGTATCGGACCTGGAAAATAATGTAGACAAATATAGAGAAGT
TATTCATAATTTAGCACAAGTTGCTCATTGTCAAGCTCAGGGTAAAATTGGAAGCGGGTTTGATGTAGCG
GCGGCAGCATATGGATCTATCAGATATAGAAGATTCCCACCCGCATTAATCTCTAATTTGCCAGATATTG
GAAGTGCTACTTACGGCAGTAAACTGGCGCATTTGGTTGATGAAGAAGACTGGAATATTACGATTAAAAG

TAACCATTTACCTTCGGGATTAACTTTATGGATGGGCGATATTAAGAATGGTTCAGAAACAGTAAAACTG
GTCCAGAAGGTAAAAAATTGGTATGATTCGCATATGCCAGAAAGCTTGAAAATATATACAGAACTCGATC
ATGCAAATTCTAGATTTATGGATGGACTATCTAAACTAGATCGCTTACACGAGACTCATGACGATTACAG
CGATCAGATATTTGAGTCTCTTGAGAGGAATGACTGTACCTGTCAAAAGTATCCTGAAATCACAGAAGTT
AGAGATGCAGTTGCCACAATTAGACGTTCCTTTAGAAAAATAACTAAAGAATCTGGTGCCGATATCGAAC
CTCCCGTACAAACTAGCTTATTGGATGATTGCCAGACCTTAAAAGGAGTTCTTACTTGCTTAATACCTGG
TGCTGGTGGTTATGACGCCATTGCAGTGATTACTAAGCAAGATGTTGATCTTAGGGCTCAAACCGCTAAT
GACAAAAGATTTTCTAAGGTTCAATGGCTGGATGTAACTCAGGCTGACTGGGGTGTTAGGAAAGAAAAAG
ATCCGGAAACTTATCTTGATAAA

**MVD1**

ATGACCGTTTACACAGCATCCGTTACCGCACCCGTCAACATCGCAACCCTTAAGTATTGGGGGAAAAGGG
ACACGAAGTTGAATCTGCCCACCAATTCGTCCATATCAGTGACTTTATCGCAAGATGACCTCAGAACGTT
GACCTCTGCGGCTACTGCACCTGAGTTTGAACGCGACACTTTGTGGTTAAATGGAGAACCACACAGCATC
GACAATGAAAGAACTCAAAATTGTCTGCGCGACCTACGCCAATTAAGAAAGGAAATGGAATCGAAGGACG
CCTCATTGCCCACATTATCTCAATGGAAACTCCACATTGTCTCCGAAAATAACTTTCCTACAGCAGCTGG
TTTAGCTTCCTCCGCTGCTGGCTTTGCTGCATTGGTCTCTGCAATTGCTAAGTTATACCAATTACCACAG
TCAACTTCAGAAATATCTAGAATAGCAAGAAAGGGGTCTGGTTCAGCTTGTAGATCGTTGTTTGGCGGAT
ACGTGGCCTGGGAAATGGGAAAAGCTGAAGATGGTCATGATTCCATGGCAGTACAAATCGCAGACAGCTC
TGACTGGCCTCAGATGAAAGCTTGTGTCCTAGTTGTCAGCGATATTAAAAAGGATGTGAGTTCCACTCAG
GGTATGCAATTGACCGTGGCAACCTCCGAACTATTTAAAGAAAGAATTGAACATGTCGTACCAAAGAGAT
TTGAAGTCATGCGTAAAGCCATTGTTGAAAAAGATTTCGCCACCTTTGCAAAGGAAACAATGATGGATTC
CAACTCTTTCCATGCCACATGTTTGGACTCTTTCCCTCCAATATTCTACATGAATGACACTTCCAAGCGT
ATCATCAGTTGGTGCCACACCATTAATCAGTTTTACGGAGAAACAATCGTTGCATACACGTTTGATGCAG
GTCCAAATGCTGTGTTGTACTACTTAGCTGAAAATGAGTCGAAACTCTTTGCATTTATCTATAAATTGTT
TGGCTCTGTTCCTGGATGGGACAAGAAATTTACTACTGAGCAGCTTGAGGCTTTCAACCATCAATTTGAA
TCATCTAACTTTACTGCACGTGAATTGGATCTTGAGTTGCAAAAGGATGTTGCCAGAGTGATTTTAACTC
AAGTCGGTTCAGGCCCACAAGAAACAAACGAATCTTTGATTGACGCAAAGACTGGTCTACCAAAGGAA

**idi**

ATGATAATGCAAACGGAACACGTCATTTTATTGAATGCACAGGGAGTTCCCACGGGTACGCTGGAAAAGT
ATGCCGCACACACGGCAGACACCCGCTTACATCTCGCGTTCTCCAGTTGGCTGTTTAATGCCAAAGGACA
ATTATTAGTTACCCGCCGCGCACTGAGCAAAAAAGCATGGCCTGGCGTGTGGACTAACTCGGTTTGTGGG
CACCCACAACTGGGAGAAAGCAACGAAGACGCAGTGATCCGCCGTTGCCGTTATGAGCTTGGCGTGGAAA
TTACGCCTCCTGAATCTATCTATCCTGACTTTCGCTACCGCGCCACCGATCCGAGTGGCATTGTGGAAAA
TGAAGTGTGTCCGGTATTTGCCGCACGCACCACTAGTGCGTTACAGATCAATGATGATGAAGTGATGGAT
TATCAATGGTGTGATTTAGCAGATGTATTACACGGTATTGATGCCACGCCGTGGGCGTTCAGTCCGTGGA
TGGTGATGCAGGCGACAAATCGCGAAGCCAGAAAACGATTATCTGCATTTACCCAGCTTAAA

**ispA**

ATGGACTTTCCGCAGCAACTCGAAGCCTGCGTTAAGCAGGCCAACCAGGCGCTGAGCCGTTTTATCGCCC
CACTGCCCTTTCAGAACACTCCCGTGGTCGAAACCATGCAGTATGGCGCATTATTAGGTGGTAAGCGCCT
GCGACCTTTCCTGGTTTATGCCACCGGTCATATGTTCGGCGTTAGCACAAACACGCTGGACGCACCCGCT
GCCGCCGTTGAGTGTATCCACGCTTACTCATTAATTCATGATGATTTACCGGCAATGGATGATGACGATC
TGCGTCGCGGTTTGCCAACCTGCCATGTGAAGTTTGGCGAAGCAAACGCGATTCTCGCTGGCGACGCTTT
ACAAACGCTGGCGTTCTCGATTTTAAGCGATGCCGATATGCCGGAAGTGTCGGACCGCGACAGAATTTCG
ATGATTTCTGAACTGGCGAGCGCCAGTGGTATTGCCGGAATGTGCGGTGGTCAGGCATTAGATTTAGACG
CGGAAGGCAAACACGTACCTCTGGACGCGCTTGAGCGTATTCATCGTCATAAAACCGGCGCATTGATTCG
CGCCGCCGTTCGCCTTGGTGCATTAAGCGCCGGAGATAAAGGACGTCGTGCTCTGCCGGTACTCGACAAG

TATGCAGAGAGCATCGGCCTTGCCTTCCAGGTTCAGGATGACATCCTGGATGTGGTGGGAGATACTGCAA
CGTTGGGAAAACGCCAGGGTGCCGACCAGCAACTTGGTAAAAGTACCTACCCTGCACTTCTGGGCCTTGA
GCAAGCCCGGAAGAAAGCCCGGGATCTGATCGACGATGCCCGTCAGTCGCTGAAACAACTGGCTGAACAG
TCACTCGATACCTCGGCACTGGAAGCGCTAGCGGACTACATCATCCAGCGTAATAAA

**ADS**

ATGGCCCTGACCGAAGAGAAACCGATCCGCCCGATCGCTAACTTCCCGCCGTCTATCTGGGGTGACCAGT
TCCTGATCTACGAAAAGCAGGTTGAGCAGGGTGTTGAACAGATCGTAAACGACCTGAAGAAAGAAGTTCG
TCAGCTGCTGAAAGAAGCTCTGGACATCCCGATGAAACACGCTAACCTGCTGAAACTGATCGACGAGATC
CAGCGTCTGGGTATCCCGTACCACTTCGAACGCGAAATCGACCACGCACTGCAGTGCATCTACGAAACCT
ACGGCGACAACTGGAACGGCGACCGTTCTTCTCTGTGGTTTCGTCTGATGCGTAAACAGGGCTACTACGT
TACCTGTGACGTTTTTAACAACTACAAGGACAAGAACGGTGCTTTCAAACAGTCTCTGGCTAACGACGTT
GAAGGCCTGCTGGAACTGTACGAAGCGACCTCCATGCGTGTACCGGGTGAAATCATCCTGGAGGACGCGC
TGGGTTTCACCCGTTCTCGTCTGTCCATTATGACTAAAGACGCTTTCTCTACTAACCCGGCTCTGTTCAC
CGAAATCCAGCGTGCTCTGAAACAGCCGCTGTGGAAACGTCTGCCGCGTATCGAAGCAGCACAGTACATT
CCGTTTTACCAGCAGCAGGACTCTCACAACAAGACCCTGCTGAAACTGGCTAAGCTGGAGTTCAACCTGC
TGCAGTCTCTGCACAAAGAAGAACTGTCTCACGTTTGTAAGTGGTGGAAGGCATTTGACATCAAGAAAAA
CGCGCCGTGCCTGCGTGACCGTATCGTTGAATGTTACTTCTGGGGTCTGGGTTCTGGTTATGAACCACAG
TACTCCCGTGCACGTGTGTTCTTCACTAAAGCTGTAGCTGTTATCACCCTGATCGATGACACTTACGATG
CTTACGGCACCTACGAAGAACTGAAGATTTTTACTGAAGCTGTAGAACGCTGGTCTATCACTTGCCTGGA
CACTCTGCCGGAGTACATGAAACCGATCTACAAACTGTTCATGGATACCTACACCGAAATGGAGGAGTTC
CTGGCAAAAGAAGGCCGTACCGACCTGTTCAACTGCGGTAAAGAGTTTGTTAAAGAGTTCGTACGTAACC
TGATGGTTGAAGCTAAATGGGCTAACGAAGGCCATATCCCGACTACCGAAGAACATGACCCGGTTGTTAT
CATCACCGGCGGTGCAAACCTGCTGACCACCACTTGCTATCTGGGTATGTCCGACATCTTTACCAAGGAA
TCTGTTGAATGGGCTGTTTCTGCACCGCCGCTGTTCCGTTACTCCGGTATTCTGGGTCGTCGTCTGAACG
ACCTGATGACCCACAAAGCAGAGCAGGAACGTAAACACTCTTCCTCCTCTCTGGAATCCTACATGAAGGA
ATATAACGTTAACGAGGAGTACGCACAGACTCTGATCTATAAAGAAGTTGAAGACGTATGGAAAGACATC
AACCGTGAATACCTGACTACTAAAAACATCCCGCGCCCGCTGCTGATGGCAGTAATCTACCTGTGCCAGT
TCCTGGAAGTACAGTACGCTGGTAAAGATAACTTCACTCGCATGGGCGACGAATACAAACACCTGATCAA
ATCCCTGCTGGTTTACCCGATGTCCATC

**nDHFR**

ATGGTTCGACCATTGAACTGCATCGTCGCCGTGTCCCAAAATATGGGGATTGGCAAGAACGGAGACCTAC
CCTGGCCTCCGCTCAGGAACGAGTTCAAGTACTTCCAAAGAATGACCACAACCTCTTCAGTGGAAGGTAA
ACAGAATCTGGTGATTATGGGTAGGAAAACCTGGTTCTCCATTCCTGAGAAGAATCGACCTTTAAAGGAC
AGAATTAATATAGTTCTCAGTAGAGAACTCAAAGAACCACCACGAGGAGCTCATTTTCTTGCCAAAAGTT
TGGATGATGCCTTAAGACTTATTGAACAACCGGAATTGGGTACC

**cDHFR**

ATGAGTAAAGTAGACATGGTTTGGATAGTCGGAGGCAGTTCTGTTTACCAGGAAGCCATGAATCAACCAG
GCCACCTCAGACTCTTTGTGACAAGGATCATGCAGGAATTTGAAAGTGACACGTTTTTCCCAGAAATTGA
TTTGGGGAAATATAAACTTCTCCCAGAATACCCAGGCGTCCTCTCTGAGGTCCAGGAGGAAAAAGGCATC
AAGTATAAGTTTGAAGTCTACGAGAAGAAAGAC

**nDHFR fusion linker**

GGGAGCAGTGCTAGCGGAACTTCTAGCACTAGTTCCGGAATT

**cDHFR fusion linker**

GGTGGCTCTGGCAGTGGAGCTAGCACT

**LgBIT**

ATGGTCTTCACACTCGAAGATTTCGTTGGGGACTGGGAACAGACAGCCGCCTACAACCTGGACCAAGTCC
TTGAACAGGGAGGTGTGTCCAGTTTGCTGCAGAATCTCGCCGTGTCCGTAACTCCGATCCAAAGGATTGT
CCGGAGCGGTGAAAATGCCCTGAAGATCGACATCCATGTCATCATCCCGTATGAAGGTCTGAGCGCCGAC
CAAATGGCCCAGATCGAAGAGGTGTTTAAGGTGGTGTACCCTGTGGATGATCATCACTTTAAGGTGATCC
TGCCCTATGGCACACTGGTAATCGACGGGGTTACGCCGAACATGCTGAACTATTTCGGACGGCCGTATGA
AGGCATCGCCGTGTTCGACGGCAAAAAGATCACTGTAACAGGGACCCTGTGGAACGGCAACAAAATTATC
GACGAGCGCCTGATCACCCCCGACGGCTCCATGCTGTTCCGAGTAACCATCAACAGC

**SmBIT**

ATGGTGACCGGCTACCGGCTGTTCGAGGAGATTCTG

**LgBIT and ddFP fusion linker**

GGTAGCGGCAGCGGCAG

**SmBIT fusion linker**

GGTAGCGGCAGCGGCAGGGTAGCGGCTTCT

**ddRFPb**

ATGGTGAGCAAGGGCGAGGAGACCATCAAAGAGTTCATGCGCTTCAAGGTGCGCATGGAGGGCTCCATGA
ACGGCCACGAGTTCGAGATCGAGGGCGAGGGCGAGGGCCGCCCCTACGAGGGCACCCAGACCGCCAAGCT
GAAGGTGACCAAGGGCGGCCCCCTGCCCTTCGCCTGGGACATCCTGTCCCCCCAGTTCATGTACGGCTCC
GAGGCGTACGTGAGGCACCCCGCCGACATCCCCGATTACAAGAAGCTGCCCTTCCCCGAGGGCTTCAAGT
GGGAGCGCGTGATGAACTTCGAAGACGGCGGTCTGGTGACCGTTACCCAGGACTCCTCCCTGCAGGACGG
CACGCTGATCTGCAAGGTGAAGATGCGCGGCACCAACTTCCCCCCCGACGGCCCCGTAATGCAGAAGAAG
ACCATGGGCTGGGAGGCCTCCACCGAGATGCTGTACCCCGAAGACGGCGTGCTGAAGGGCCATAGCTATC
AGGCCCTGAAGCTGAAGGACGGCGGCCACTACCTGGTGGAGTTCGAGACCATCTACATGGCCAAGAAGCC
CGTGCAACTGCCCGGCGATTACTGTGTGGACACCAAGCTGGACATCACCTCCCACAACGAGGACTACACC
ATCGTGGAACAGTACGGGCGCTCCGAGGGCCGCCACCGTCTGGGCATGGACGAGCGGTACAAG

**ddGFPa**

ATGGCGAGCAAGAGCGAGGAGGTCATCAAAGAGTTCATGCGCTTCAAGGTGCGCTTGGAGGGCTCCATGA
ACGGCCACGAGTTCGAGATCGAGGGCGAGGGCGAGGGCCGCCCTTACGAGGGCACCCAGACCGCCAAGCT
GAAGGTGACCAAGGGCGGCCCCCTGCCCTTCGCCTGGGACATCCTGTCCCCCCTGATCATGTACGGCTCC
AAGATGTACGTGAAGCACCCCGCCGACGTCCCCGATTACATGAAGCTGTCCTTCCCCGAGGGCTTCAAGT
GGGAGCGCGTGATGCACTTCGAGGACGGCGGTCTGGTGACCGCGACACAGGACTCCTCCCTGCAGGACGG
CACGCTGATCTACAAGGTGAAGATGCGCGGCACCAACTTCCCCCCCGACGGCCCCGTAATGCAGAAGAAG
ACCTTGGGCTGGGATTATGCCACCGAGCGCCTGTACCCCGAAGAAGGCGTGCTGAAGGGCGAGCTCCTGG
GGCGCCTGAAGCTGAAGGACGGCGGCCTCAACCTGGTGGAGTCCAAGACCATCTACATGGCCAAGAAGCC
CGTGCAACTGCCCGGCTACTACTTCGTGGACACCAAGCTGGACATCACCTCCCACAACGAGGACTACACC
ATCGTGGAACAGTACGAGCGCTCCGAGGGCCGCCACCACCTGGGGCATGGTACAGGTAGCACAGGCAGC

**T7 RNA polymerase**

ATGAACACGATTAACATCGCTAAGAACGACTTCTCTGACATCGAACTGGCTGCTATCCCGTTCAACACTC
TGGCTGACCATTACGGTGAGCGTTTAGCTCGCGAACAGTTGGCCCTTGAGCATGAGTCTTACGAGATGGG
TGAAGCACGCTTCCGCAAGATGTTTGAGCGTCAACTTAAAGCTGGTGAGGTTGCGGATAACGCTGCCGCC
AAGCCTCTCATCACTACCCTACTCCCTAAGATGATTGCACGCATCAACGACTGGTTTGAGGAAGTGAAAG
CTAAGCGCGGCAAGCGCCCGACAGCCTTCCAGTTCCTGCAAGAAATCAAGCCGGAAGCCGTAGCGTACAT
CACCATTAAGACCACTCTGGCTTGCCTAACCAGTGCTGACAATACAACCGTTCAGGCTGTAGCAAGCGCA
ATCGGTCGGGCCATTGAGGACGAGGCTCGCTTCGGTCGTATCCGTGACCTTGAAGCTAAGCACTTCAAGA
AAAACGTTGAGGAACAACTCAACAAGCGCGTAGGGCACGTCTACAAGAAAGCATTTATGCAAGTTGTCGA
GGCTGACATGCTCTCTAAGGGTCTACTCGGTGGCGAGGCGTGGTCTTCGTGGCATAAGGAAGACTCTATT
CATGTAGGAGTACGCTGCATCGAGATGCTCATTGAGTCAACCGGAATGGTTAGCTTACACCGCCAAAATG
CTGGCGTAGTAGGTCAAGACTCTGAGACTATCGAACTCGCACCTGAATACGCTGAGGCTATCGCAACCCG
TGCAGGTGCGCTGGCTGGCATCTCTCCGATGTTCCAACCTTGCGTAGTTCCTCCTAAGCCGTGGACTGGC
ATTACTGGTGGTGGCTATTGGGCTAACGGTCGTCGTCCTCTGGCGCTGGTGCGTACTCACAGTAAGAAAG
CACTGATGCGCTACGAAGACGTTTACATGCCTGAGGTGTACAAAGCGATTAACATTGCGCAAAACACCGC
ATGGAAAATCAACAAGAAAGTCCTAGCGGTCGCCAACGTAATCACCAAGTGGAAGCATTGTCCGGTCGAG
GACATCCCTGCGATTGAGCGTGAAGAACTCCCGATGAAACCGGAAGACATCGACATGAATCCTGAGGCTC
TCACCGCGTGGAAACGTGCTGCCGCTGCTGTGTACCGCAAGGACAAGGCTCGCAAGTCTCGCCGTATCAG
CCTTGAGTTCATGCTTGAGCAAGCCAATAAGTTTGCTAACCATAAGGCCATCTGGTTCCCTTACAACATG
GACTGGCGCGGTCGTGTTTACGCTGTGTCAATGTTCAACCCGCAAGGTAACGATATGACCAAAGGACTGC
TTACGCTGGCGAAAGGTAAACCAATCGGTAAGGAAGGTTACTACTGGCTGAAAATCCACGGTGCAAACTG
TGCGGGTGTCGATAAGGTTCCGTTCCCTGAGCGCATCAAGTTCATTGAGGAAAACCACGAGAACATCATG
GCTTGCGCTAAGTCTCCACTGGAGAACACTTGGTGGGCTGAGCAAGATTCTCCGTTCTGCTTCCTTGCGT
TCTGCTTTGAGTACGCTGGGGTACAGCACCACGGCCTGAGCTATAACTGCTCCCTTCCGCTGGCGTTTGA
CGGGTCTTGCTCTGGCATCCAGCACTTCTCCGCGATGCTCCGAGATGAGGTAGGTGGTCGCGCGGTTAAC
TTGCTTCCTAGTGAAACCGTTCAGGACATCTACGGGATTGTTGCTAAGAAAGTCAACGAGATTCTACAAG
CAGACGCAATCAATGGGACCGATAACGAAGTAGTTACCGTGACCGATGAGAACACTGGTGAAATCTCTGA
GAAAGTCAAGCTGGGCACTAAGGCACTGGCTGGTCAATGGCTGGCTTACGGTGTTACTCGCAGTGTGACT
AAGCGTTCAGTCATGACGCTGGCTTACGGGTCCAAAGAGTTCGGCTTCCGTCAACAAGTGCTGGAAGATA
CCATTCAGCCAGCTATTGATTCCGGCAAGGGTCTGATGTTCACTCAGCCGAATCAGGCTGCTGGATACAT
GGCTAAGCTGATTTGGGAATCTGTGAGCGTGACGGTGGTAGCTGCGGTTGAAGCAATGAACTGGCTTAAG
TCTGCTGCTAAGCTGCTGGCTGCTGAGGTCAAAGATAAGAAGACTGGAGAGATTCTTCGCAAGCGTTGCG
CTGTGCATTGGGTAACTCCTGATGGTTTCCCTGTGTGGCAGGAATACAAGAAGCCTATTCAGACGCGCTT
GAACCTGATGTTCCTCGGTCAGTTCCGCTTACAGCCTACCATTAACACCAACAAAGATAGCGAGATTGAT
GCACACAAACAGGAGTCTGGTATCGCTCCTAACTTTGTACACAGCCAAGACGGTAGCCACCTTCGTAAGA
CTGTAGTGTGGGCACACGAGAAGTACGGAATCGAATCTTTTGCACTGATTCACGACTCCTTCGGTACCAT
TCCGGCTGACGCTGCGAACCTGTTCAAAGCAGTGCGCGAAACTATGGTTGACACATATGAGTCTTGTGAT
GTACTGGCTGATTTCTACGACCAGTTCGCTGACCAGTTGCACGAGTCTCAATTGGACAAAATGCCAGCAC
TTCCGGCTAAAGGTAACTTGAACCTCCGTGACATCTTAGAGTCGGACTTCGCGTTCGCG

**Supplementary Notes and References**

1. Adapted from: Mandell, D.J., Backbone Flexibility in Computational Design, PhD Dissertation, UCSF, ProQuest Dissertations Publishing, 3412217, (2010).
2. A. Zanghellini, L. Jiang, A. M. Wollacott, G. Cheng, J. Meiler, E. A. Althoff, D. Rothlisberger, D. Baker, New algorithms and an in silico benchmark for computational enzyme design. *Protein Sci* **15**, 2785-2794 (2006).
3. R. L. Dunbrack, Jr., F. E. Cohen, Bayesian statistical analysis of protein side-chain rotamer preferences. *Protein Sci* **6**, 1661-1681 (1997).
4. I. W. Davis, D. Baker, RosettaLigand docking with full ligand and receptor flexibility. *J Mol Biol* **385**, 381-392 (2009).
5. D. Rothlisberger, O. Khersonsky, A. M. Wollacott, L. Jiang, J. DeChancie, J. Betker, J. L. Gallaher, E. A. Althoff, A. Zanghellini, O. Dym, S. Albeck, K. N. Houk, D. S. Tawfik, D. Baker, Kemp elimination catalysts by computational enzyme design. *Nature* **453**, 190-195 (2008).
6. L. Jiang, E. A. Althoff, F. R. Clemente, L. Doyle, D. Rothlisberger, A. Zanghellini, J. L. Gallaher, J. L. Betker, F. Tanaka, C. F. Barbas, 3rd, D. Hilvert, K. N. Houk, B. L. Stoddard, D. Baker, De novo computational design of retro-aldol enzymes. *Science* **319**, 1387-1391 (2008).
7. D. J. Mandell, E. A. Coutsias, T. Kortemme, Sub-angstrom accuracy in protein loop reconstruction by robotics-inspired conformational sampling. *Nat Methods* **6**, 551-552 (2009).
8. M. Babor, D. J. Mandell, T. Kortemme, Assessment of flexible backbone protein design methods for sequence library prediction in the therapeutic antibody Herceptin-HER2 interface. *Protein Sci* **20**, 1082-1089 (2011).
9. E. L. Humphris, T. Kortemme, Prediction of protein-protein interface sequence diversity using flexible backbone computational protein design. *Structure* **16**, 1777-1788 (2008).
10. N. Ollikainen, R. M. de Jong, T. Kortemme, Coupling Protein Side-Chain and Backbone Flexibility Improves the Re-design of Protein-Ligand Specificity. *PLoS Comput Biol* **11**, e1004335 (2015).
11. C. A. Smith, T. Kortemme, Backrub-like backbone simulation recapitulates natural protein conformational variability and improves mutant side-chain prediction. *J Mol Biol* **380**, 742-756 (2008).
12. P. Conway, M. D. Tyka, F. DiMaio, D. E. Konerding, D. Baker, Relaxation of backbone bond geometry improves protein energy landscape modeling. *Protein Sci* **23**, 47-55 (2014).
13. J. N. Pelletier, F. X. Campbell-Valois, S. W. Michnick, Oligomerization domain-directed reassembly of active dihydrofolate reductase from rationally designed fragments. *Proc Natl Acad Sci U S A* **95**, 12141-12146 (1998).
14. D. M. Hoover, J. Lubkowski, DNAWorks: an automated method for designing oligonucleotides for PCR-based gene synthesis. *Nucleic Acids Res* **30**, e43 (2002).
15. C. Engler, R. Kandzia, S. Marillonnet, A one pot, one step, precision cloning method with high throughput capability. *PLoS One* **3**, e3647 (2008).
16. M. E. Lee, W. C. DeLoache, B. Cervantes, J. E. Dueber, A Highly Characterized Yeast Toolkit for Modular, Multipart Assembly. *ACS Synth Biol* **4**, 975-986 (2015).

17. Z. Z. Sun, C. A. Hayes, J. Shin, F. Caschera, R. M. Murray, V. Noireaux, Protocols for implementing an Escherichia coli based TX-TL cell-free expression system for synthetic biology. *J Vis Exp*, e50762 (2013).
18. A. S. Dixon, M. K. Schwinn, M. P. Hall, K. Zimmerman, P. Otto, T. H. Lubben, B. L. Butler, B. F. Binkowski, T. Machleidt, T. A. Kirkland, M. G. Wood, C. T. Eggers, L. P. Encell, K. V. Wood, NanoLuc Complementation Reporter Optimized for Accurate Measurement of Protein Interactions in Cells. *ACS Chem Biol* **11**, 400-408 (2016).
19. H. K. Binz, P. Amstutz, A. Kohl, M. T. Stumpp, C. Briand, P. Forrer, M. G. Grutter, A. Pluckthun, High-affinity binders selected from designed ankyrin repeat protein libraries. *Nat Biotechnol* **22**, 575-582 (2004).
20. G. Winter, xia2: an expert system for macromolecular crystallography data reduction. *Journal of Applied Crystallography* **43**, 186-190 (2010).
21. W. Kabsch, Xds. *Acta Crystallogr D Biol Crystallogr* **66**, 125-132 (2010).
22. P. Evans, Scaling and assessment of data quality. *Acta Crystallogr D Biol Crystallogr* **62**, 72-82 (2006).
23. J. E. Padilla, T. O. Yeates, A statistic for local intensity differences: robustness to anisotropy and pseudo-centering and utility for detecting twinning. *Acta Crystallogr D Biol Crystallogr* **59**, 1124-1130 (2003).
24. A. J. McCoy, R. W. Grosse-Kunstleve, P. D. Adams, M. D. Winn, L. C. Storoni, R. J. Read, Phaser crystallographic software. *J Appl Crystallogr* **40**, 658-674 (2007).
25. P. D. Adams, P. V. Afonine, G. Bunkoczi, V. B. Chen, I. W. Davis, N. Echols, J. J. Headd, L. W. Hung, G. J. Kapral, R. W. Grosse-Kunstleve, A. J. McCoy, N. W. Moriarty, R. Oeffner, R. J. Read, D. C. Richardson, J. S. Richardson, T. C. Terwilliger, P. H. Zwart, PHENIX: a comprehensive Python-based system for macromolecular structure solution. *Acta Crystallogr D Biol Crystallogr* **66**, 213-221 (2010).
26. K. A. Kantardjieff, B. Rupp, Matthews coefficient probabilities: Improved estimates for unit cell contents of proteins, DNA, and protein-nucleic acid complex crystals. *Protein Sci* **12**, 1865-1871 (2003).
27. C. X. Weichenberger, B. Rupp, Ten years of probabilistic estimates of biocrystal solvent content: new insights via nonparametric kernel density estimate. *Acta Crystallogr D Biol Crystallogr* **70**, 1579-1588 (2014).
28. P. V. Afonine, R. W. Grosse-Kunstleve, N. Echols, J. J. Headd, N. W. Moriarty, M. Mustyakimov, T. C. Terwilliger, A. Urzhumtsev, P. H. Zwart, P. D. Adams, Towards automated crystallographic structure refinement with phenix.refine. *Acta Crystallogr D Biol Crystallogr* **68**, 352-367 (2012).
29. M. C. Thompson, Identifying and Overcoming Crystal Pathologies: Disorder and Twinning. *Methods Mol Biol* **1607**, 185-217 (2017).
30. P. Emsley, B. Lohkamp, W. G. Scott, K. Cowtan, Features and development of Coot. *Acta Crystallogr D Biol Crystallogr* **66**, 486-501 (2010).
31. N. W. Moriarty, R. W. Grosse-Kunstleve, P. D. Adams, electronic Ligand Builder and Optimization Workbench (eLBOW): a tool for ligand coordinate and restraint generation. *Acta Crystallogr D Biol Crystallogr* **65**, 1074-1080 (2009).
32. F. C. Bernstein, T. F. Koetzle, G. J. Williams, E. F. Meyer, Jr., M. D. Brice, J. R. Rodgers, O. Kennard, T. Shimanouchi, M. Tasumi, The Protein Data Bank: a computer-based archival file for macromolecular structures. *J Mol Biol* **112**, 535-542 (1977).

33. M. A. Kramer, S. K. Wetzel, A. Pluckthun, P. R. Mittl, M. G. Grutter, Structural determinants for improved stability of designed ankyrin repeat proteins with a redesigned C-capping module. *J Mol Biol* **404**, 381-391 (2010).
34. P. A. Karplus, K. Diederichs, Linking crystallographic model and data quality. *Science* **336**, 1030-1033 (2012).
35. A. T. Brunger, Free R value: cross-validation in crystallography. *Methods Enzymol* **277**, 366-396 (1997).
36. V. B. Chen, W. B. Arendall, 3rd, J. J. Headd, D. A. Keedy, R. M. Immormino, G. J. Kapral, L. W. Murray, J. S. Richardson, D. C. Richardson, MolProbity: all-atom structure validation for macromolecular crystallography. *Acta Crystallogr D Biol Crystallogr* **66**, 12-21 (2010).
